# A chemical screen based on an interruption of zebrafish gastrulation identifies the HTR2C inhibitor Pizotifen as a suppressor of EMT-mediated metastasis

**DOI:** 10.1101/2021.03.04.434001

**Authors:** Joji Nakayama, Lora Tan, Boon Cher Goh, Shu Wang, Hideki Makinoshima, Zhiyuan Gong

## Abstract

Metastasis is responsible for approximately 90% of cancer-associated mortality but few models exist that allow for rapid and effective screening of anti-metastasis drugs. Current mouse models of metastasis are too expensive and time consuming to use for rapid and high-throughput screening. Therefore, we created a unique screening concept utilizing conserved mechanisms between zebrafish gastrulation and cancer metastasis for identification of potential anti-metastatic drugs. We hypothesized that small chemicals that interrupt zebrafish gastrulation might also suppress metastatic progression of cancer cells and developed a phenotype-based chemical screen to test the hypothesis. The screen used epiboly, the first morphogenetic movement in gastrulation, as a marker and enabled 100 chemicals to be tested in five hours. The screen tested 1280 FDA-approved drugs and identified Pizotifen, an antagonist for serotonin receptor 2C (HTR2C) as an epiboly-interrupting drug. Pharmacologic and genetic inhibition of HTR2C suppressed metastatic progression in a mouse model. Blocking HTR2C with Pizotifen restored epithelial properties to metastatic cells through inhibition of Wnt-signaling. In contrast, HTR2C induced epithelial to mesenchymal transition (EMT) through activation of Wnt-signaling and promoted metastatic dissemination of human cancer cells in a zebrafish xenotransplantation model. Taken together, our concept offers a novel platform for discovery of anti-metastasis drugs.

## Introduction

Metastasis, a leading contributor to the morbidity of cancer patients, occurs through multiple steps: invasion, intravasation, extravasation, colonization, and metastatic tumor formation (1–3). The physical translocation of cancer cells is an initial step of metastasis and molecular mechanisms of it involve cell motility, the breakdown of local basement membrane, loss of cell polarity, acquisition of stem cell-like properties, and EMT (4–6). These cell-biological phenomena are also observed during vertebrate gastrulation in that evolutionarily conserved morphogenetic movements of epiboly, internalization, convergence, and extension progress (7). In zebrafish, the first morphogenetic movement, epiboly, is initiated at approximately 4 hours post fertilization (hpf) to move cells from the animal pole to eventually engulf the entire yolk cell by 10 hpf (8, 9). The embryonic cell movements are governed by the molecular mechanisms that are partially shared in metastatic cell dissemination.

At least fifty common genes were shown to be involved in both metastasis and gastrulation progression: Knockdown of these genes in Xenopus or zebrafish induced gastrulation defects; conversely, overexpression of these genes conferred metastatic potential on cancer cells and knockdown of these genes suppressed metastasis (Table S1). This evidence led us to hypothesize that small molecules that interrupt zebrafish gastrulation may suppress metastatic progression of human cancer cells.

Here we report a unique screening concept based on the hypothesis. Pizotifen, an antagonist for HTR2C, was identified from the screen as a “hit” that interrupted zebrafish gastrulation. A mouse model of metastasis confirmed pharmacological and genetic inhibition of HTR2C suppressed metastatic progression. Moreover, HTR2C induced EMT and promoted metastatic dissemination of non-metastatic cancer cells in a zebrafish xenotransplantation model. These results demonstrated that this concept could offer a novel high-throughput platform for discovery of anti-metastasis drugs and can be converted to a chemical genetic screening platform.

## Results

### Small molecules interrupting epiboly of zebrafish have a potential to suppress metastatic progression of human cancer cells

Before performing a screening assay, we conducted preliminary experiments to test the hypothesis. First, we examined whether hindering the molecular function of reported genes, whose knockdown induced gastrulation defects in zebrafish, might suppress cell motility and invasion of cancer cells. We chose protein arginine methyltransferase 1 (PRMT1) and cytochrome P450 family 11 (CYP11A1), both of whose knockdown induced gastrulation defects in zebrafish but whose involvement in metastatic progression is unclear (10, 11). Elevated expression of PRMT1 and CYP11A1 were observed in highly metastatic human breast cancer cell lines and knockdown of these genes through RNA interference suppressed the motility and invasion of MDA-MB-231 cells without affecting their viability (Figure S1A-C).

Next, we conducted an inverse examination of whether chemicals which were reported to suppress metastatic dissemination of cancer cells could interrupt epiboly progression of zebrafish embryos. Niclosamide and Vinpocetine are reported to suppress metastatic progression (12, 13) (14, 15). Either Niclosamide or Vinpocetine-treated zebrafish embryos showed complete arrest at very early stages or severe delay in epiboly progression, respectively (Figure S1D).

These results suggest that epiboly could serve as a marker for this screening assay and epiboly-interrupting drugs that are identified through this screening could have the potential to suppress metastatic progression of human cancer cells.

### 132 FDA-approved drugs induced delayed in epiboly of zebrafish embryos

We screened 1,280 FDA, EMA or other agencies-approved drugs (Prestwick, Inc) in our zebrafish assay. The screening showed that 0.9% (12/1280) of the drugs, including Actimycin A and Tolcapone, induced severe or complete arrest of embryonic cell movement when embryos were treated with 10μM. 5.2% (66/1280) of the drugs, such as Dicumarol, Racecadotril, Pizotifen and S(-) Eticlopride hydrochloride, induced either delayed epiboly or interrupted epiboly of the embryos. 93.3% (1194/1280) of drugs has no effect on epiboly progression of the embryos. 0.6% (8/1280) of drugs induced a toxic lethality. Epiboly progression was affected more severely when embryos were treated with 50μM; 1.7% (22/1280) of the drugs induced severe or complete arrest of it. 8.6% (110/1280) of the drugs induced either delayed epiboly or interrupt epiboly of the embryos. 4.3% (55/1280) of drugs induced a toxic lethality (Figure 1A and 1B, Table S2). Among the epiboly-interrupting drugs, several drugs have already been reported to inhibit metastasis-related molecular mechanisms: Adrenosterone or Zardaverine, which target HSD11β1 or PDE3 and 4, respectively, are reported to inhibit EMT (16, 17); Racecadotril, which targets enkephalinase, is reported to confer metastatic potential on colon cancer cell (18); and Disulfiram, which targets ALDH, is reported to confer stem-like properties on metastatic cancer cells (19). This evidence suggests that epiboly-interrupting drugs have the potential for suppressing metastasis of human cancer cells.

**Figure 1.**
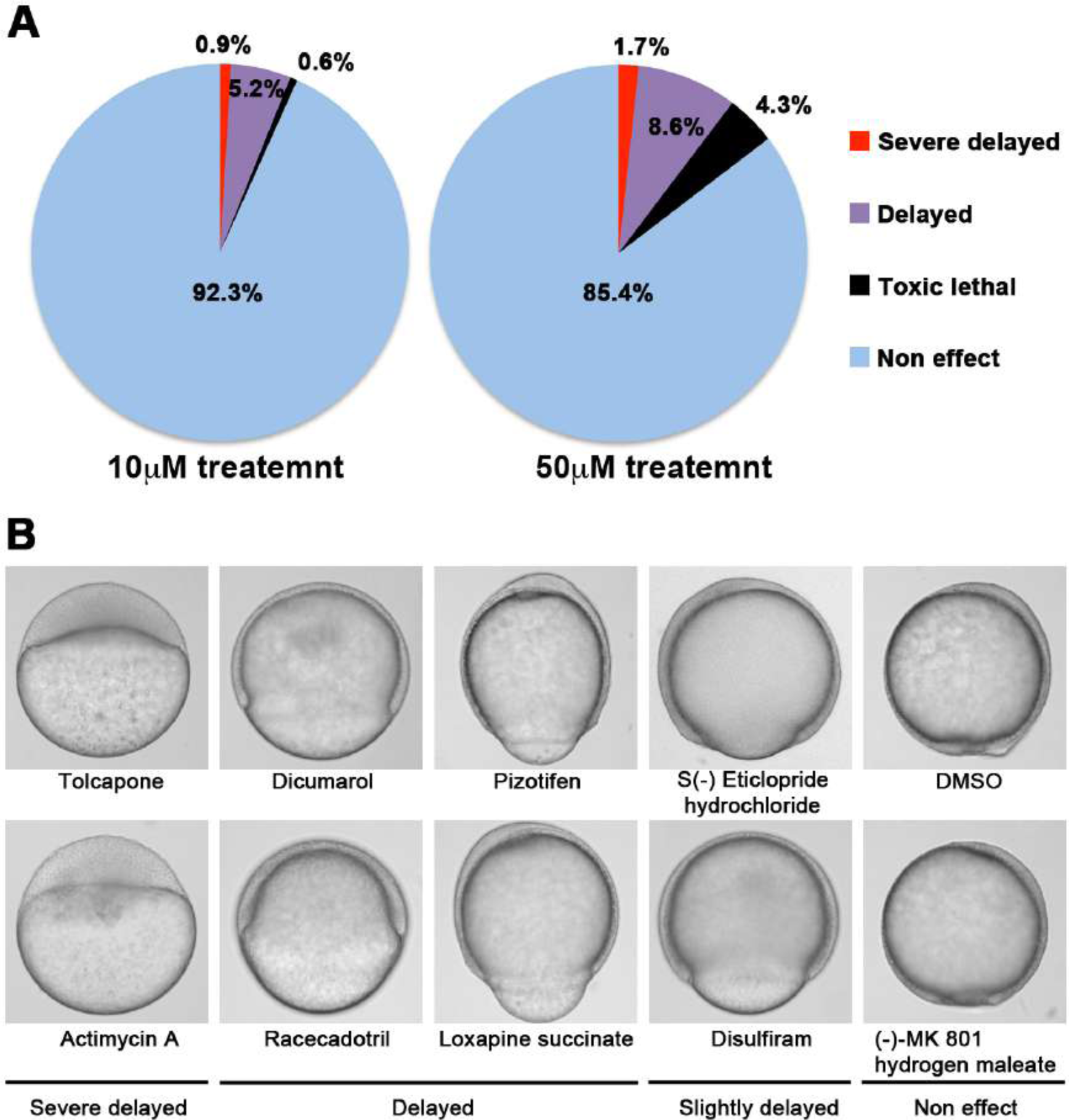
A chemical screen for identification of epiboly-interrupting drugs. (A) Cumulative results of the chemical screen in which each drug was used at either 10µM (left) or 50µM (right) concentrations. 1,280 FDA, EMA or other agencies-approved drugs were subjected to this screening. Positive “hit” drugs were those that interrupted epiboly progression. (B) Representative samples of the embryos that were treated with indicated drugs.

### Identified drugs suppressed cell motility and invasion of human cancer cells

It has been reported that zebrafish have orthologues to 86% of 1318 human drug targets (20). However, it was not known whether the epiboly-interrupting drugs could suppress metastatic dissemination of human cancer cells. To test this, we subjected the 78 epiboly-interrupting drugs that showed a suppressor effect on epiboly progression at a 10μM concentration to *in vitro* experiments using a human cancer cell line. The experiments examined whether the drugs could suppress cell motility and invasion of MDA-MB-231 cells through a Boyden chamber. Before conducting the experiment, we investigated whether these drugs might effect viability of MDA-MB-231 cells using an MTT assay.

Sixteen of the 78 drugs strongly effected cell viability at concentrations less than 1μM and were not used in the cell motility experiments. The remaining 62 drugs were assayed in Boyden chamber motility experiments. Twenty of the 62 drugs inhibited cell motility and invasion of MDA-MB-231 cells without effecting cell viability. Among the 20 drugs, Hexachlorophene and Nitazoxanide were removed since the primary targets of the drugs, D-lactate dehydrogenase and pyruvate ferredoxin oxidoreductase are not expressed in mammalian cells. With the exception of Ipriflavone, whose target is still unclear, the known primary targets of the remaining 17 drugs are reported to be expressed by mammalian cells (Figure 2A and Table 1).

**Figure 2.**
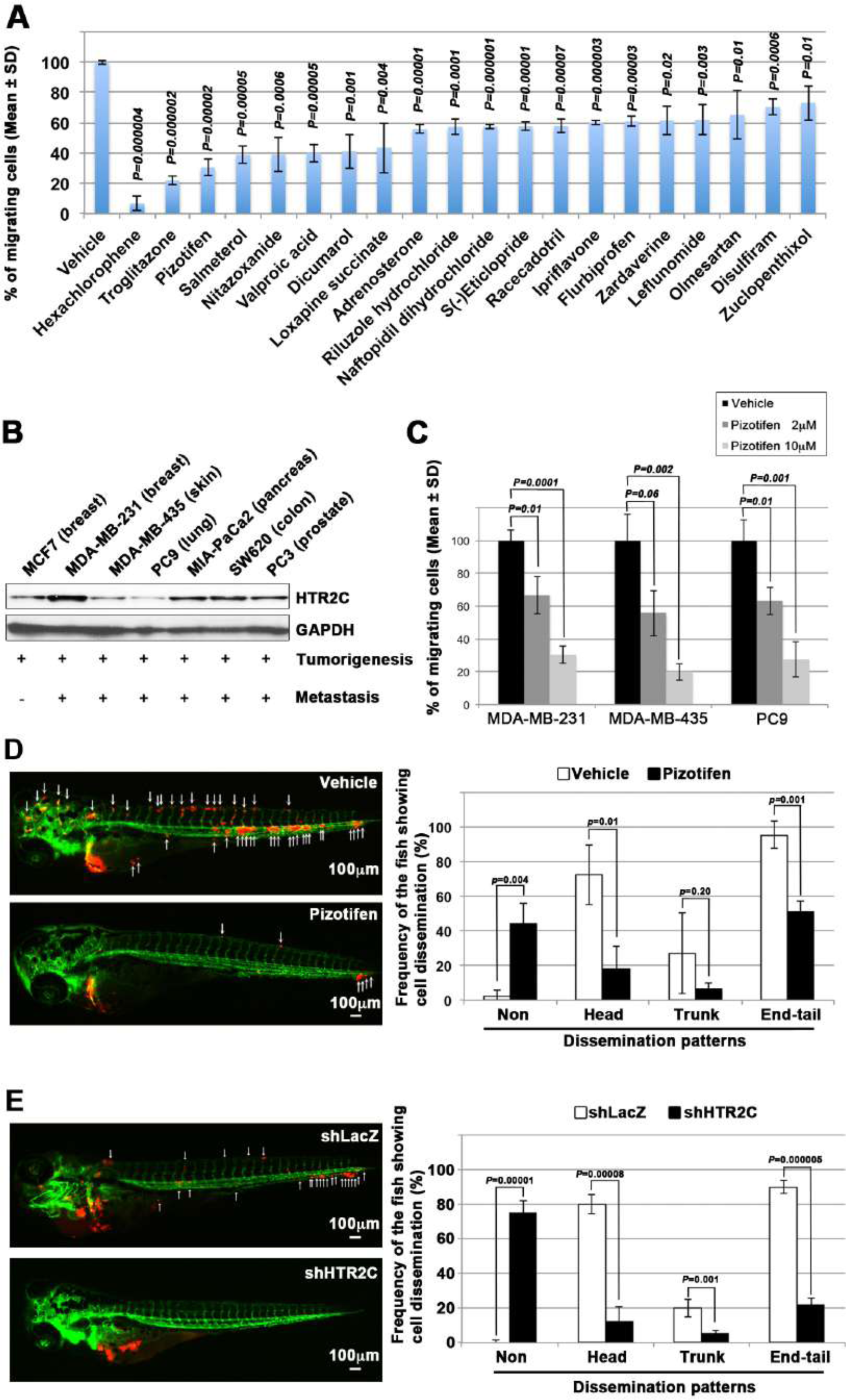
Pizotifen, one of epiboly-interrupting drugs suppressed metastatic dissemination of human cancer cells lines in vivo and vitro. (A) Effect of the epiboly-interrupting drugs on cell motility and invasion of MBA-MB-231 cells. MBA-MB-231 cells were treated with vehicle or each of the epiboly-interrupting drugs and then subjected to Boyden chamber assays. Fetal bovine serum (1%v/v) was used as the chemoattractant in both assays. Each experiment was performed at least twice. (B) Western blot analysis of HTR2C levels (top) in a non-metastatic human cancer cell line, MCF7 (breast) and highly metastatic human cancer cell lines, MDA-MB-231 (breast), MDA-MB-435 (melanoma), PC9 (lung), MIA-PaCa2 (pancreas), PC3 (prostate) and SW620 (colon); GAPDH loading control is shown (bottom). (C) Effect of pizotifen on cell motility and invasion of MBA-MB-231, MDA-MB-435 and PC9 cells. Either vehicle or pizotifen treated the cells were subjected to Boyden chamber assays. Fetal bovine serum (1%v/v) was used as the chemoattractant in both assays. Each experiment was performed at least twice. (D) and (E) Representative images of dissemination of 231R, shLacZ 231R or shHTR2C 231R cells in zebrafish xenotransplantation model. The fish larvae that were inoculated with 231R cells, were treated with either vehicle (top left) or the drug (lower left) (D). The fish larvae that were inoculated with either shLacZ 231R or shHTR2C 231R cells (lower left) (E). White arrows head indicate disseminated 231R cells. The images were shown in 4x magnification. Scale bar, 100µm. The mean frequencies of the fish showing head, trunk, or end-tail dissemination were counted (graph on right). Each value is indicated as the mean ± SEM of two independent experiments. Statistical analysis was determined by Student’s t test. See also Figure S2 and S3, Table S3-S5.

**Table 1.** Primary targets of the identified drugs

| The identified drugs | Primary targets of the identified drugs |
| --- | --- |
| Hexachlorophene | D-lactate dehydrogenase (D-LDH), not expressed in mammalian cells |
| Troglitazone | Agonist for Peroxisome proliferator-activated receptor $\alpha$ and $\gamma$ (PPARα and $\gamma$) |
| Pizotifen malate | 5-hydroxytryptamine receptor 2C (HTR2C) |
| Salmeterol | Adrenergic receptor beta 2 (ADRB2) |
| Nitazoxanide | Pyruvate ferredoxin oxidoreductase (PFOR), not expressed in mammalian cells |
| Valproic acid | Histone deacetylases (HDACs) |
| Dicumarol | NAD(P)H dehydrogenase quinone 1 (NQO1) |
| Loxapine succinate | Dopamine receptor D2 and D4 (DRD2 and DRD4) |
| Adrenosterone | Hydroxysteroid (11-beta) dehydrogenase 1 (HSD11β1) |
| Riluzole hydrochloride | Glutamate R and Voltage-dependent Na<sup>+</sup> channel |
| Naftopidil dihydrochloride | 5-hydroxytryptamine receptor 1A (HTR1A) and α1-adrenergic receptor (AR) |
| S(-)Eticlopride hydrochloride | Dopamine receptor D2 (DRD2) |
| Racecadotril | Membrane metallo-endopeptidase (MME) |
| Ipriflavone | Unknow |
| Flurbiprofen | Cyclooxygenase 1 and 2 (Cox1 and 2) |
| Zardaverine | Phosphodiesterase III/IV (PDE3/4) |
| Leflunomide | Dihydroorotate dehydrogenase (DHODH) |
| Olmesartan | Angiotensin II receptor alpha |
| Disulfiram | Aldehyde dehydrogenase (ALDH)<br>Dopamine β-hydroxylase (DBH) |
| Zuclopenthixol dihydrochloride | Dopamine receptors D1 and D2 (DRD1 and 2) |

We confirmed if highly metastatic human cancer cell lines expressed the genes that code for these targets using Western blotting analyses. Among the genes, serotonin receptor 2C (HTR2C), which is a primary target of Pizotifen, was highly expressed in only metastatic cell lines (Figure 2B). Pizotifen suppressed cell motility and invasion of several highly metastatic human cancer cell lines in a dose-dependent manner (Figure 2C). Similarly, Dopamine receptor D2 (DRD2), which is a primary target of S(-) Eticlopride hydrochloride, was highly expressed in only metastatic cell lines, and the drug suppressed cell motility and invasion of these cells in a dose-dependent manner (Figure S2).

These results indicate that a number of the epiboly-interrupting drugs also have suppressor effects on cell motility and invasion of highly metastatic human cancer cells.

### Pizotifen suppressed metastatic dissemination of human cancer cells in a zebrafish xenotransplantation model

While a number of the epiboly-interrupting drugs suppressed cell motility and invasion of human cell lines *in vitro,* it was still unclear whether the drugs could suppress metastatic dissemination of cancer cells in vivo. Therefore, we examined whether the identified drugs could suppress metastatic dissemination of these human cancer cells in a zebrafish xenotransplantation model. Pizotifen was selected to test since HTR2C was overexpressed only in highly metastatic cell lines supporting the hypothesis that it could be a novel target for blocking metastatic dissemination of cancer cells (Figure 2B). Red fluorescent protein-labelled MDA-MB-231 (231R) cells were injected into the duct of Cuvier of *Tg (kdrl:eGFP)* zebrafish at 2 dpf and then maintained in the presence of either vehicle or Pizotifen. Twenty-four hours post-injection, the numbers of fish showing metastatic dissemination of 231R cells were measured via fluorescence microscopy. In this model, the dissemination patterns were generally divided into three categories: (i) head dissemination, in which disseminated 231R cells exist in the vessel of the head part; (ii) trunk dissemination, in which the cells were observed in the vessel dilating from the trunk to the tail; (iii) end-tail dissemination, in which the cells were observed in the vessel of the end-tail part (16).

Three independent experiments revealed that the frequencies of fish in the drug-treated group showing head, trunk, or end-tail dissemination, significantly decreased to 55.3±7.5%, 28.5±5.0% or 43.5±19.1% when compared with those in the vehicle-treated group; 95.8±5.8%, 47.1±7.7% or 82.6±12.7%. Conversely, the frequency of the fish in the drug-treated group not showing any dissemination, significantly increased to 45.4±0.5% when compared with those in the vehicle-treated group; 2.0±2.9% (Figure 2D and Table S3).

Similar effects were observed in another xenograft experiments using an RFP-labelled human pancreatic cancer cell line, MIA-PaCa-2 (MP2R). In the drug treated group, the frequencies of the fish showing head, trunk or end-tail dissemination, significantly decreased to 15.3±6.7%, 6.2±1.3%, or 41.1±1.5%; conversely, the frequency of the fish not showing any dissemination significantly increased to 46.3±8.9% when compared with those in the vehicle-treated group; 74.5±11.1%, 18.9±14.9%, 77.0±9.0%, or 17.2±0.7% (Figure S3 and Table S4).

To eliminate the possibility that the metastasis suppressing effects of Pizotifen might result from off-target effects of the drug, we conducted validation experiments to determine whether knockdown of HTR2C would show the same effects. Sub-clones of 231R cells that expressed shRNA targeting either LacZ or HTR2C were injected into the fish at 2 dpf and the fish were maintained in the absence of drug. In the fish that were inoculated with shHTR2C 231R cells, the frequencies of the fish showing head, trunk, and end-tail dissemination, significantly decreased to 6.7±4.9%, 6.7±0.7%, or 20.0±16.5%; conversely, the frequency of the fish not showing any dissemination, significantly increased to 80.0±4.4% when compared with those that were inoculated with shLacZ 231R cells; 80.0±27.1%, 20.0±4.5%, 90.0± 7.7%, or 0% (Figure 2E and Table S5).

These results indicate that pharmacological and genetic inhibition of HTR2C suppressed metastatic dissemination of human cancer cells in vivo.

### Pizotifen suppressed metastasis progression of a mouse model of metastasis

We examined the metastasis-suppressor effect of Pizotifen in a mouse model of metastasis (21). Luciferase-expressing 4T1 murine mammary carcinoma cells were inoculated into the mammary fat pads (MFP) of female BALB/c mice. On day two post inoculation, the mice were randomly assigned to two groups and one group received once daily intraperitoneal injections of 10mg/kg Pizotifen while the other group received a vehicle injection. Bioluminescence imaging and tumor measurement revealed that the sizes of the primary tumors in Pizotifen-treated mice were equal to those in the vehicle-treated mice at the time of resection on day 10 post inoculation. Immunofluorescent staining also demonstrated that the percentage of Ki67 positive cells in the resected primary tumors of Pizotifen-treated mice were the same as those of vehicle-treated mice (Figure 3A-C), additionally, both groups showed less than 1% cleaved caspase 3 positive cells (data not shown). Therefore, no anti-tumor effect of Pizotifen was observed on the primary tumor. After 70 days from inoculation, bioluminescence imaging detected light emitted in the lungs, livers and lymph nodes of vehicle-treated mice but not those of Pizotifen-treated mice (Figure 3C). Vehicle-treated mice formed 5 to 50 metastatic nodules per lung in all 10 mice analyzed; conversely, Pizotifen-treated mice (n=10) formed 0 to 5 nodules per lung in all 10 mice analyzed (Figure 3D). Histological analyses confirmed that metastatic lesions in the lungs were detected in all vehicle-treated mice; conversely, they were detected in only 2 of 10 Pizotifen-treated mice and the rest of the mice showed metastatic colony formations around the bronchiole of the lung. In addition, 4 of 10 vehicle-treated mice exhibited metastasis in the liver and the rest showed metastatic colony formation around the portal tract of the liver. In contrast, none of 10 Pizotifen-treated mice showed liver metastases and only half of the 10 mice showed metastatic colony formation around the portal tract (Figure 3E). These results indicate that Pizotifen can suppress metastasis progression without affecting primary tumor growth.

**Figure 3.**
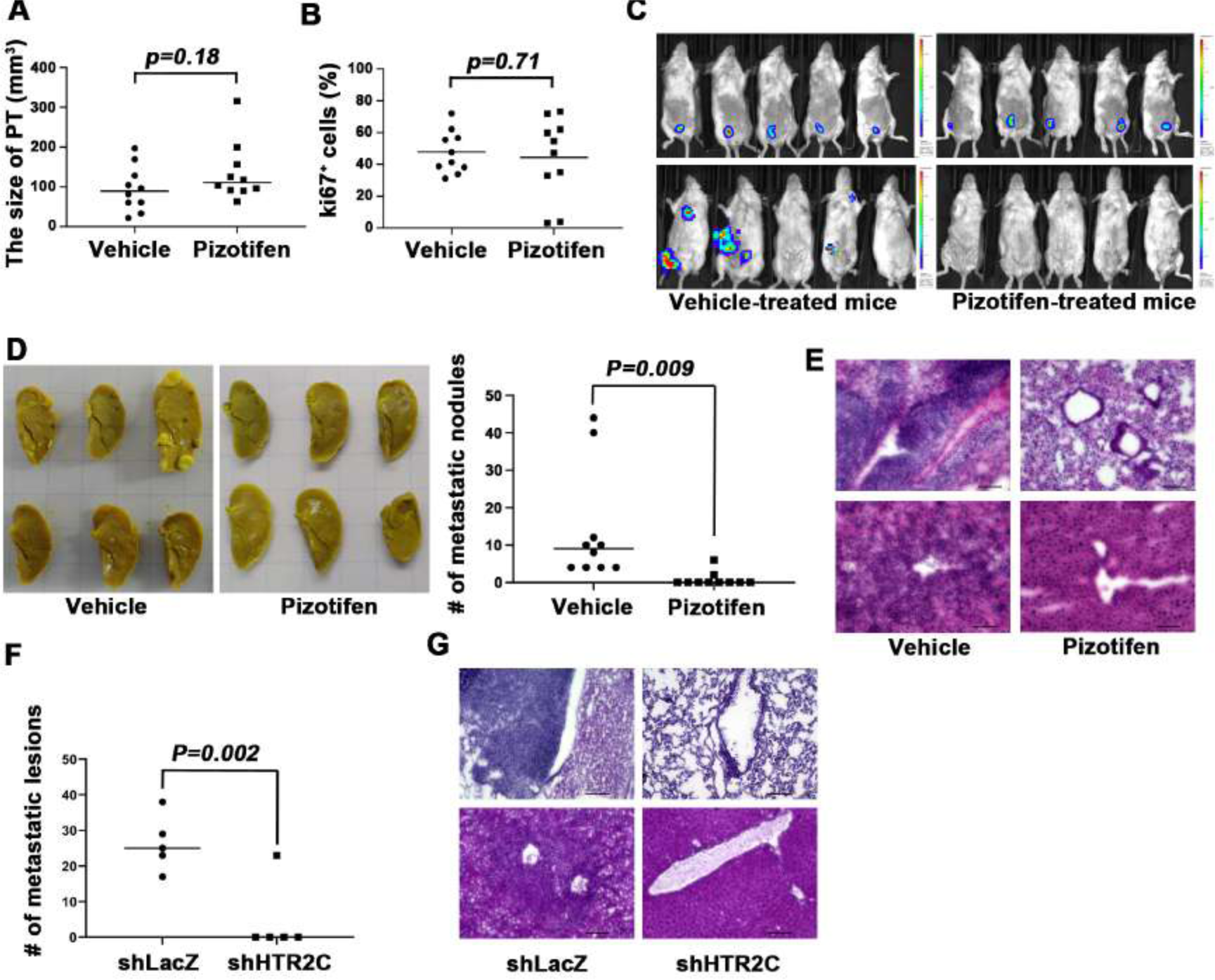
Pizotifen suppressed metastatic progression in a mouse model of metastasis. (A) Mean volumes (n=10 per group) of 4T1 primary tumors formed in the mammary fat pad of either vehicle or Pizotifen-treated mice at day 10 post injection. (B) Ki67 expression level in 4T1 primary tumors formed in the mammary fat pad of either vehicle or Pizotifen-treated mice at day 10 post injection. The mean expression levels of Ki67 (n=10 mice per group) were determined and were calculated as the mean ration of Ki67 positive cells to DAPI area. (C) Representative images of primary tumors on day 10 post injection (top panels) and metastatic burden on day 70 post injection (bottom panels) taken using an IVIS Imaging System. (D) Representative images of the lungs from either vehicle (top) or Pizotifen-treated mice (bottom) at 70 days past tumor inoculation. Number of metastatic nodules in the lung of either vehicle or Pizotifen-treated mice (right). (E) Representative H&E staining of the lung (top) and liver (bottom) from either vehicle or Pizotifen-treated mice. (F) The mean number of metastatic lesions in step sections of the lungs from the mice that were inoculated with 4T1-12B cells expressing shRNA targeting for either LacZ or HTR2C. (G) Representative H&E staining of the lung and liver from the mice that were inoculated with 4T1-12B cells expressing shRNA targeting for either LacZ or HTR2C. Each value is indicated as the mean ± SEM. Statistical analysis was determined by Student’s *t* test.

To eliminate the possibility that the metastasis suppressing effects of Pizotifen might result from off-target effects, we conducted validation experiments to determine whether knockdown of HTR2C would show the same effects. The basic experimental process followed the experimental design described above except that sub-clones of 4T1 cells that expressed shRNA targeting either LacZ or HTR2C were injected into the MFP of female BALB/c mice and the mice were maintained without drug. Histological analyses revealed that all of the mice (n=5) that were inoculated with 4T1 cells expressing shRNA targeting LacZ showed metastases in the lungs. The mean number of metastatic lesions in a lung was 26.4±7.8. In contrast, only one of the mice (n=5) were inoculated with 4T1 cells expressing shRNA targeting HTR2C showed metastases in the lungs and the rest of the mice showed metastatic colony formation around the bronchiole of the lung. The mean number of metastatic lesions in the lung significantly decreased to 10% of those of mice that were inoculated with 4T1 cells expressing shRNA targeting LacZ (Figure 3F-H).

Taken together, pharmacological and genetic inhibition of HTR2C showed an anti-metastatic effect in the 4T1 model system.

### HTR2C promoted EMT-mediated metastatic dissemination of human cancer cells

Although pharmacological and genetic inhibition of HTR2C inhibited metastasis progression, a role for HTR2C on metastatic progression has not been reported.

Therefore, we examined whether HTR2C could confer metastatic properties on poorly metastatic cells.

Firstly, we established a stable sub-clone of MCF7 human breast cancer cells expressing either vector control or HTR2C. Vector control expressing MCF7 cells maintained highly organized cell-cell adhesion and cell polarity; however, HTR2C-expressing MCF7 cells led to loss of cell-cell contact and cell scattering. The cobblestone-like appearance of these cells was replaced by a spindle-like, fibroblastic morphology.

Western blotting and immunofluorescence (IF) analyses revealed that HTR2C-expressing MCF7 cells showed loss of E-cadherin and EpCAM, and elevated expressions of N-cadherin, vimentin and an EMT-inducible transcriptional factor Zeb1. Similar effects were validated through another experiment using an immortal keratinocyte cell line, HaCaT cells, in that HTR2C-expressing HaCaT cells also showed loss of cell-cell contact and cell scattering with loss of epithelial markers and gain of mesenchymal markers (Figure 4A-C). Therefore, both the morphological and molecular changes in the HTR2C-expressing MCF7 and HaCaT cells demonstrated that these cells had undergone an EMT.

**Figure 4.**
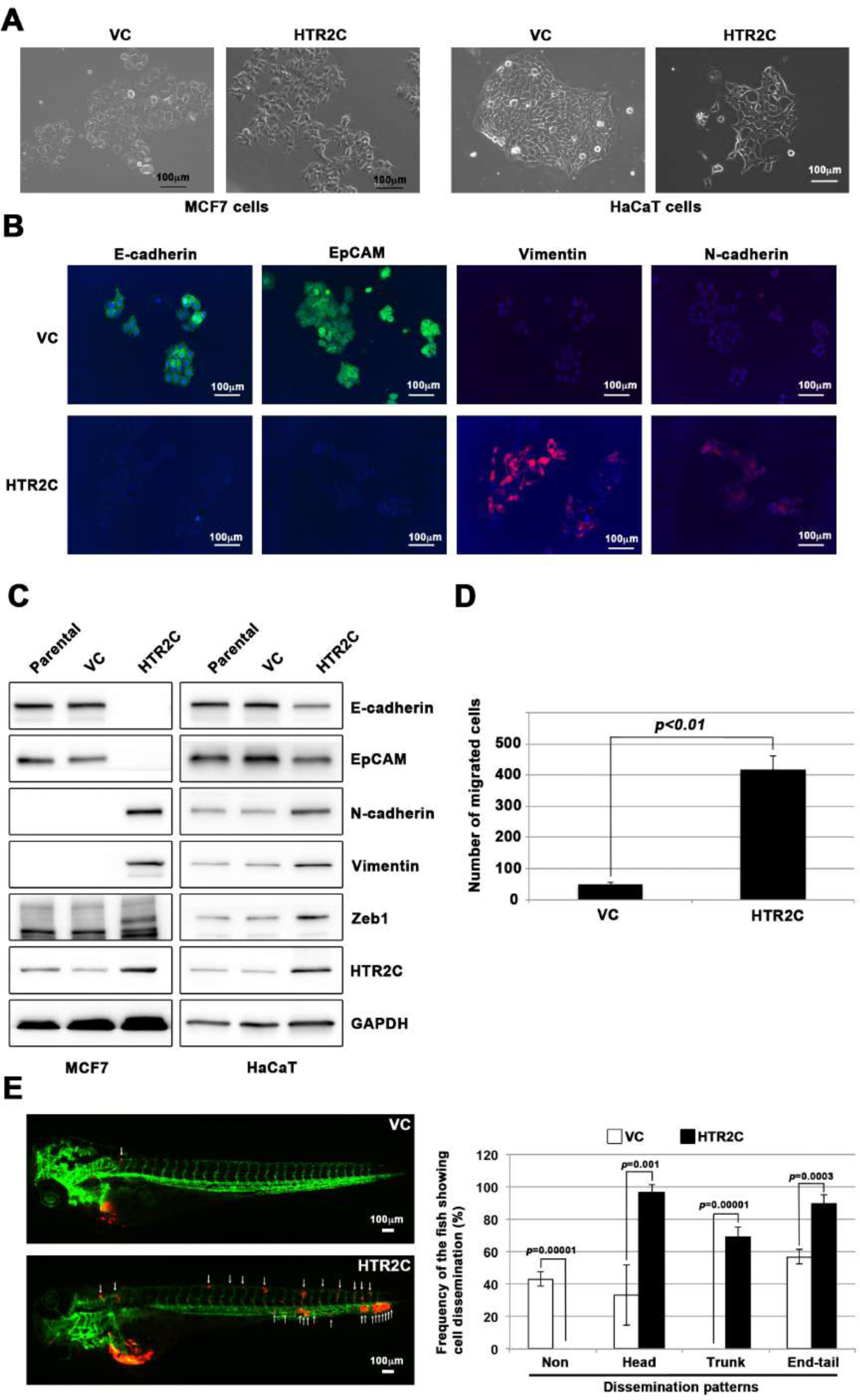
HTR2C induced EMT-mediated metastatic dissemination of human cancer cells. (A) The morphologies of the MCF7 and HaCaT cells expressing either the control vector or HTR2C were revealed by phase contrast microscopy. (B) Immunofluorescence staining of E-cadherin, EpCAM, Vimentin, and N-cadherin expressions in the MCF7 cells from Figure 4A. (C) Expression of E-cadherin, EpCAM, Vimentin, N-cadherin, and HTR2C was examined by western-blotting in the MCF7 and HaCaT cells; GAPDH loading control is shown (bottom). (D) Effect of HTR2C on cell motility and invasion of MCF7 cells. MCF7 cells were subjected to Boyden chamber assays. Fetal bovine serum (1%v/v) was used as the chemoattractant in both assays. Each experiment was performed at least twice. (E) Representative images of dissemination patterns of MCF7 cells expressing either the control vector (top left) or HTR2C (lower left) in a zebrafish xenotransplantation model. White arrows head indicate disseminated MCF7 cells. The mean frequencies of the fish showing head, trunk, or end-tail dissemination tabulated (right). Each value is indicated as the mean ± SEM of two independent experiments. Statistical analysis was determined by Student’s *t* test. See also Table S6.

Next, we examined whether HTR2C-driven EMT could promote metastatic dissemination of human cancer cells. Boyden chamber assay revealed that HTR2C expressing MCF7 cells showed an increased cell motility and invasion compared with vector control-expressing MCF7 cells in vitro (Figure 4D). Moreover, we conducted in vivo examination of whether HTR2C expression could promote metastatic dissemination of human cancer cells in a zebrafish xenotransplantation model. Red fluorescence protein-labelled MCF7 cells expressing either vector control or HTR2C were injected into the duct of Cuvier of *Tg* (*kdrl:eGFP*) zebrafish at 2 dpf. Twenty-four hours post injection, the frequencies of the fish showing metastatic dissemination of the inoculated cells were measured using fluorescence microscopy. In the fish that were inoculated with HTR2C expressing MCF7 cells, the frequencies of the fish showing head, trunk, and end-tail dissemination significantly increased to 96.7±4.7%, 68.8±6.4%, or 89.5±3.4%; conversely, the frequency of the fish not showing any dissemination decreased to 0% when compared with those in the fish that were inoculated with vector control expressing MCF7 cells; 33.1±18.5%, 0%, 56.9± 4.4%, or 43% (Figure 4E and Table S6).

These results indicated that HTR2C promoted metastatic dissemination of cancer cells though induction of EMT, and suggest that the screen can easily be converted to a chemical genetic screening platform.

### Pizotifen induced mesenchymal to epithelial transition through inhibition of Wnt-signaling

Finally, we elucidated the mechanism of action of how Pizotifen suppressed metastasis, especially metastatic dissemination of cancer cells. Our results showed that HTR2C induced EMT and that pharmacological and genetic inhibition of HTR2C suppressed metastatic dissemination of MDA-MB-231 cells that had already transitioned to mesenchymal-like traits via EMT. Therefore, we speculated that blocking HTR2C with Pizotifen might inhibit the molecular mechanisms which follow EMT induction. We firstly investigated the expressions of epithelial and mesenchymal markers in Pizotifen-treated MDA-MB-231 cells since the activation of an EMT program needs to be transient and reversible, and transition from a fully mesenchymal phenotype to a epithelial-mesenchymal hybrid state or a fully epithelial phenotype is associated with malignant phenotypes (22). IF and FACS analyses revealed 20% of Pizotifen-treated MDA-MB-231 cells restored E-cadherin expression. Also, western blotting analysis demonstrated that 4T1 primary tumors from Pizotifen-treated mice has elevated E-cadherin expression compared with tumors from vehicle-treated mice (Figure 5A-C). However, mesenchymal markers did not change between vehicle and Pizotifen-treated MDA-MB-231 cells (data not shown). We further analyzed E-cadherin positive (E-cad^+^) cells in Pizotifen-treated MDA-MB-231 cells. The E-cad^+^ cells showed elevated expressions of epithelial markers KRT14 and KRT19; and decreased expression of mesenchymal makers vimentin, MMP1, MMP3, and S100A4. Recent research reports that an EMT program needs to be transient and reversible and that a mesenchymal phenotype in cancer cells is achieved by constitutive ectopic expression of Zeb1. In accordance with the research, the E-cad^+^ cells and 4T1 primary tumors from Pizotifen-treated mice had decreased Zeb1 expression compared with vehicle-treated cells and tumors from vehicle-treated mice (Figure 5D). In contrast, HTR2C-expressing MCF7 and HuMEC cells expressed Zeb1 but not vehicle control MCF7 and HuMEC cells (Figure 4C). These results indicate that HTR2C-mediated signaling induced EMT through up-regulation of Zeb1 and blocking HTR2C with Pizotifen induced mesenchymal to epithelial transition through downregulation of Zeb1.

**Figure 5.**
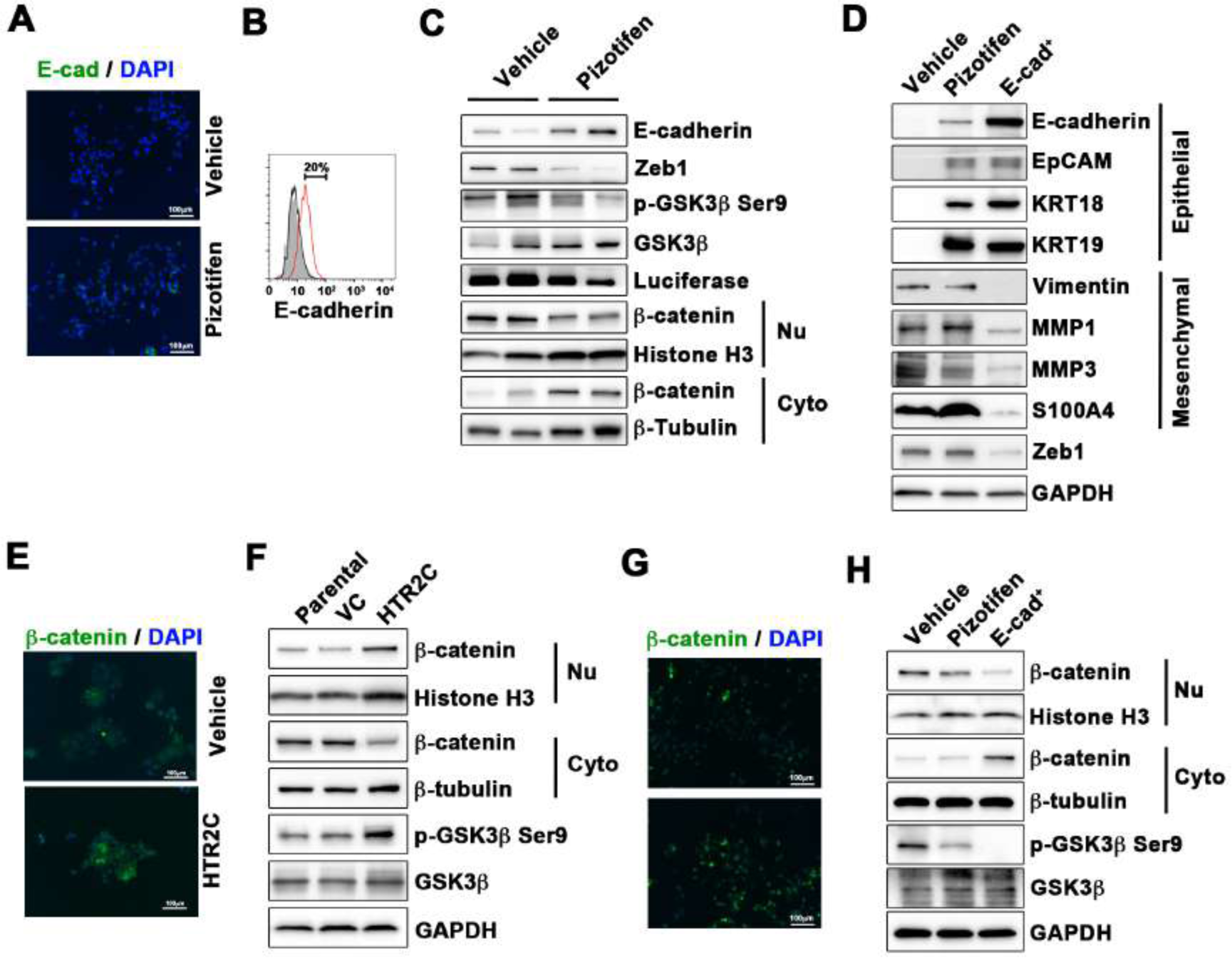
Pizotifen restored mesenchymal-like traits of MDA-MB-231 cells into epithelial traits through blocking nuclear accumulation of β-catenin. (A) IF staining of E-cadherin in either vehicle or Pizotifen-treated MDA-MB-231 cells. (B) Surface expression of E-cadherin in either vehicle (black) or Pizotifen (red)-treated MDA-MB-231 cells by FACS analysis. Non-stained controls are shown in gray. (C) Protein expressions levels of E-cadherin, Zeb1, and β-catenin in the cytoplasm and nucleus of 4T1 primary tumors from either vehicle or Pizotifen-treated mice are shown; Luciferase, Histone H3, and β-tubulin are used as loading control for whole cell, nuclear or cytoplasmic lysate, respectively. (D) Protein expression levels of epithelial and mesenchymal markers and Zeb1 in either vehicle or Pizotifen-treated MDA-MB-231 cells or E-cadherin positive (E-cad^+^) cells in Pizotifen-treated MDA-MB-231 cells are shown. (E) Immunofluorescence staining of β-catenin in the MCF7 cells expressing either vector control or HTR2C. (F) Expressions of β-catenin in the cytoplasm and nucleus of MCF7 cells. (G) IF staining of β-catenin in either vehicle or Pizotifen-treated MDA-MB-231 cells. (H) Expressions of β-catenin in the cytoplasm and nucleus of MDA-MB-231 cells and the E-cad^+^ cells.

We further investigated the mechanism of action of how blocking HTR2C with Pizotifen induced down-regulation of Zeb1. In embryogenesis, serotonin-mediated signaling is required for Wnt-dependent specification of the superficial mesoderm during gastrulation (23). In cancer cells, overexpression of HTR1D is associated with Wnt-signaling which enables induction of EMT (24, 25). This evidence led to a hypothesis that HTR2C-mediated signaling might turn on transcriptional activity of β-catenin and that might induce up-regulation of EMT-TFs. IF analyses revealed β-catenin was accumulated in the nucleus of HTR2C-expressing MCF7 cells but it was located in the cytoplasm of vector control-expressing cells (Figure 5E). Nuclear accumulation of β-catenin in HTR2C-expressing MCF7 cells was confirmed by western blot (Figure 5F). In contrast, Pizotifen-treated MDA-MB-231 cells showed β-catenin located in the cytoplasm of the cells. Vehicle-treated cells showed β-catenin accumulated in the nucleus of the cells. (Figure 5G), and western blotting analysis confirmed that it was located in the cytoplasm of Pizotifen-treated MDA-MB-231 cells (Figure 5H). Furthermore, immunohistochemistry and western blotting analyses showed that β-catenin accumulated in the nucleus, and phospho-GSKβ and Zeb1 expression were decreased in 4T1 primary tumors from Pizotifen-treated mice compared with vehicle-treated mice (Figure 5C).

These results indicated that HTR2C would regulate transcriptional activity of β-catenin and Pizotifen could inhibit it.

Taken together, we conclude that blocking HTR2C with Pizotifen restored epithelial properties to metastatic cells (MDA-MB-231 and 4T1 cells) through a decrease of transcriptional activity of β-catenin and that suppressed metastatic progression of the cells.

## Discussion

Reducing or eliminating mortality associated with metastatic disease is a key goal of medical oncology, but few models exist that allow for rapid, effective screening of novel compounds that target the metastatic dissemination of cancer cells. Based on accumulated evidence that at least fifty genes play an essential role in governing both metastasis and gastrulation progression (Table 1S), we hypothesized that small molecule inhibitors that interrupt gastrulation of zebrafish embryos might suppress metastatic progression of human cancer cells. We created a unique screening concept utilizing gastrulation of zebrafish embryos to test the hypothesis. Our results clearly confirmed our hypothesis: 25.6% (20/76 drugs) of epiboly-interrupting drugs could also suppress cell motility and invasion of highly metastatic human cell lines in vitro. In particular, Pizotifen which is an antagonist for serotonin receptor 2C and one of the epiboly-interrupting drugs, could suppress metastasis in a mouse model (Figure 3A-E). Thus, this screen could offer a novel platform for discovery of anti-metastasis drugs.

There are at least two advantages to the screen described herein. One is that the screen can easily be converted to a chemical genetic screening platform. Indeed, we have provided the first evidence that HTR2C, which is a primary target of Pizotifen, induced EMT and promoted metastatic dissemination of cancer cells (Figure 4A-E). In this research, 1,280 FDA approval drugs were screened, this is less than a few percent of all of druggable targets (approximately 100 targets) in the human proteome in the body. If chemical genetic screening using specific inhibitor libraries were conducted, more genes that contribute to metastasis and gastrulation could be identified. The second advantage is that the screen enables one researcher to test 100 drugs in 5 hours with using optical microscopy, drugs, and zebrafish embryos. That indicates this screen is not only highly efficient, low-cost, and low-labor but also enables researchers who do not have high throughput screening instruments to conduct drug screening for anti-metastasis drugs.

## Acknowledgments

We sincerely appreciate Dr. Joshua Collins (NIH/NIDCR) and Dr. Diane Palmieri (NIH/NCI) for helping this research. We thank Dr. Herrick (Albany medical collage) for providing pCMV-h5TH2C-VSV with us. This study was funded by grants from National Medical Research Council of Singapore (R-154000547511) and Ministry of Education of Singapore (R-154000A23112) to Z.G.

## Author Contributions

Design research; J.N. Conducting experiments; J.N. and L.T. Analyzing data: J.N. Writing the paper; J.N. and Z.G. Funding Acquisition; Z.G, S.W. B.C.G., H.M. and Supervision; Z.G.

## Materials and Methods

### Zebrafish embryo screening

Zebrafish embryos at two cell stage were collected at 20 mins after their fertilization. Each drug was added to a well of a 24-well plate containing approximately 20 zebrafish embryos per well in either 10 µM or 50 µM final concentration when the embryos reached the sphere stage. Chemical treatment was initiated at 4 hours post-fertilization (hpf) and approximately 20 embryos were treated with two different concentrations for each compound tested. The treatment was ended at 9 hpf when vehicle (DMSO) treated embryos as control reach 80-90% completion of the epiboly stage. The compounds which induced delay (<50% epiboly) in epiboly were selected as hit compounds for in vitro testing using highly metastatic human cancer cell lines. The study protocol was approved by the Institutional Animal Care and Use Committee of the National University of Singapore (protocol number: R16-1068).

### Reagents

FDA, EMA and other agencies approved chemical libraries was purchased from Prestwick Chemical (Illkirch, France). Pizotifen and S(-) Eticlopride hydrochloride were purchased from Sigma-Aldrich (St. Louis, MO).

### Cell culture and cell viability assay

MCF7, MDA-MB-231, MDA-MB-435, MIA-PaCa2, PC3, SW620, PC9 and HaCaT cells were obtained from American Type Culture Collection (ATCC, Manassas, VA). Luciferase-expressing 4T1 (4T1-12B) cells were provided from Dr. Gary Sahagian (Tufts University, Boston, MA). All culture methods followed the supplier’s instruction. Cell viability assay was performed as previously described (16).

### Plasmid

A DNA fragment coding for HTR2C was amplified by PCR with primers containing restriction enzyme recognition sequences. The HTR2C coding fragment was amplified from hsp70l:mCherry-T2A-CreERT2 plasmid (17).

### Immunoblotting

Western blotting was performed as described previously (16). Anti-PRMT1, anti-CYP11A1, anti-E-cadherin, anti-EpCAM, anti-Vimentin, anti-N-cadherin, anti-Zeb1, anti-Histone H3, anti-*α*-tubulin and anti-GAPDH antibodies were purchased from Cell signaling Technology. Anti-HTR2C and anti-DRD2 antibodies were purchased form Abcam. Anti-phospho-GSK3β (Ser9), anti-GSK3β, Anti-KRT18, anti-KRT19, anti-MMP1, anti-MMP2, anti-S100A4, anti-Luciferase, and anti-β-catenin antibodies were purchased from Santa Cruz.

### Flow cytometry

Cells were stained with FITC-conjugated E-cadherin antibody (Biolegend, San Diego, CA). Flow cytometry was performed as described (26) and analyzed with FlowJo software (TreeStar, Ashland, OR).

### shRNA mediated gene knockdown

The short hairpin RNA (shRNA)-expressing lentivirus vectors were constructed using pLVX-shRNA1 vector (Clontech). PRMT1-shRNA_#3–targeting sequence is GTGTTCCAGTATCTCTGATTA; PRMT1-shRNA_#4–targeting sequence is TTGACTCCTACGCACACTTTG. CYP11A1-shRNA_#4–targeting sequence is GCGATTCATTGATGCCATCTA; CYP11A1-shRNA_#4–targeting sequence is GAAATCCAACACCTCAGCGAT. Human HTR2C-shRNA–targeting sequence is TCATGCACCTCTGCGCTATAT. Mouse HTR2C-shRNA–targeting sequence is CTTCATACCGCTGACGATTAT. LacZ-shRNA– targeting sequence is CTACACAAATCAGCGATT.

### Immunofluorescence

Immunofluorescence microscopy assay was performed by previously described (16). Goat anti-mouse and goat anti-rabbit immunoglobulin G (IgG) antibodies conjugated to Alexa Fluor 488 (Life Technologies) and diluted at 1:100 were used. Nuclei were visualized by the addition of 2μg/ml of 4’, 6-diamidino-2-phenylindole (DAPI) and photographed at 100x magnification by a fluorescent microscope BZ-X700 (KEYENCE, Japan).

### Boyden chamber cell motility and invasion assay

These assays were performed by previously described (16). In Boyden chamber assay, either 3x10^5^ MDA-MB-231, 1x10^6^ MDA-MB-435, or 5x10^5^ PC9 cells were applied to each well in the upper chamber.

### Zebrafish xenotransplantation model

*Tg(kdrl:eGFP)* zebrafish was provided by Dr. Stainier (Max Planck Institute for Heart and Lung Research). Embryos that were derived from the line were maintained in E3 medium containing 200μM 1-phenyl-2-thiourea (PTU). Approximately 100-400 Red fluorescence protein (RFP)-labelled MBA-MB-231 or MIA-Paca2 cells were injected into the duct of Cuvier of the zebrafish at 2dpf. The fish were randomly assigned to two groups. One group was maintained in the presence of pizotifen-containing E3 medium and the other group was maintained in vehicle-containing E3 medium.

### Spontaneous metastasis mouse model

4T1-12B cells (2x10^4^) were injected into the #4 mammary fat pad while the mice were anesthetized. To monitor tumor growth and metastases, mice were imaged biweekly by IVIS Imaging System (ParkinElmer). The primary tumor was resected 10 days after inoculation. The study protocol (protocol number: BRC IACUC #110612) was approved by A*STAR (Agency for Science, Technology and Research, Singapore).

### Histological Analysis

All OCT embedded primary tumors, lungs, and livers of mice from the spontaneous metastasis 4T1 model were sectioned on a cryostat. Eight micron sections were taken at five hundred micron intervals through the entirety of the livers and lungs. Sections were subsequently stained with hematoxylin and eosin. Metastatic lesions were counted under a microscope in each section for both lungs and livers.

### Statistics

Data were analyzed by Student’s t test; p < 0.05 was considered significant.

## Supplemental Information

**Figure S1.**
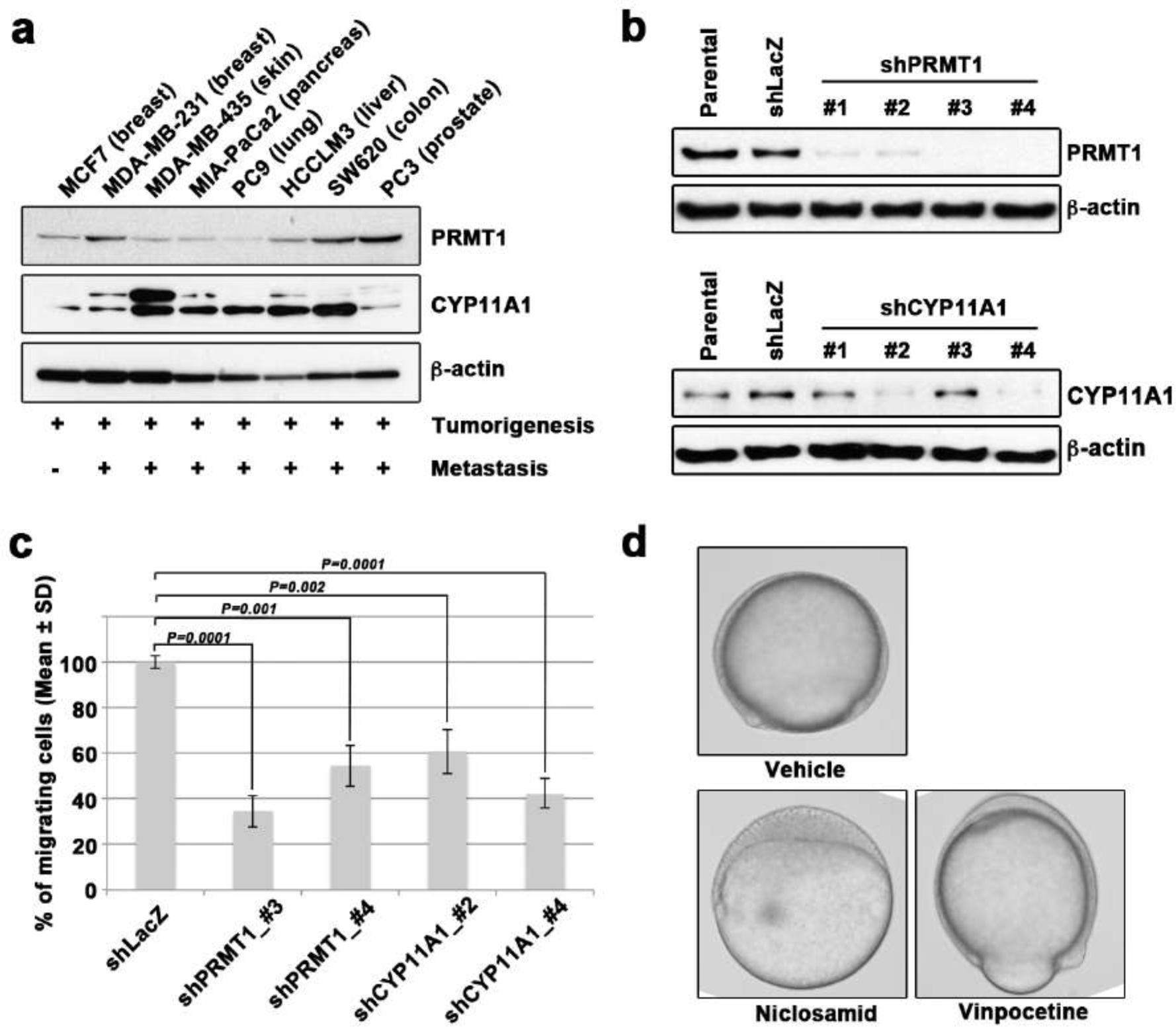
Molecular mechanisms of epiboly in zebrafish overlap with those in cancer metastasis. (A) Western blot analysis of PRMT1 (upper) and CYP11A1 (middle) protein levels in non-metastatic human cancer cell line (MCF7) and highly metastatic human cancer cell lines (MDA-MB-231, MDA-MB-435, MIA-PaCa2, PC9, HCCLM3, PC3 and SW620); β-actin loading control is shown (bottom). (B) Knockdown of PRMT1 or CYP11A1 in MDA-MB-231 cells. MBA-MB-231 cells were transfected with a control shRNA targeting LacZ, and one of four independent shRNAs targeting PRMT1 (clone #1 to #4) or one of two independent shRNAs targeting CYP11A1 (clone #1 to #4). Reduced PRMT1 and CYP11A1 expression, determined by western blot, in sub-cell lines of MDA-MB-231 cells expressing PRMT1 shRNA (clone #3 and #4 or CYP11A1 (clone #2 and #4), compared with controls (parental cell line MDA-MB-231 and control shRNA cells); β-actin levels shown as a loading control. (C) Effect of shRNAs targeting either PRMT1 or CYP11A1 on cell motility and invasion of MBA-MB-231 cells. Parental MDA-MB-231 cells and four sub-cell lines of MDA-MB-231 cells that were transfected with either shRNA targeting either LacZ, two independent shRNAs targeting PRMT1 (clone #3 and #4) or two independent shRNAs targeting CYP11A1 (clone #2 and #4), were subjected to Boyden chamber assays. (D) Zebrafish embryos treated with either vehicle (DMSO), 10μM Niclosamide, or 50µM vinpocetine. Approximately 20 embryos were treated with either DMSO as a vehicle control, niclosamide, or vinpocetine. The treatment was started at 4 hpf when all of embryos reached sphere stage and ended at 9 hpf when control embryos reached 80-90% epiboly stage. Each experiment was performed at least twice. Statistical analysis was determined by Student’s *t* test.

**Figure S2.**
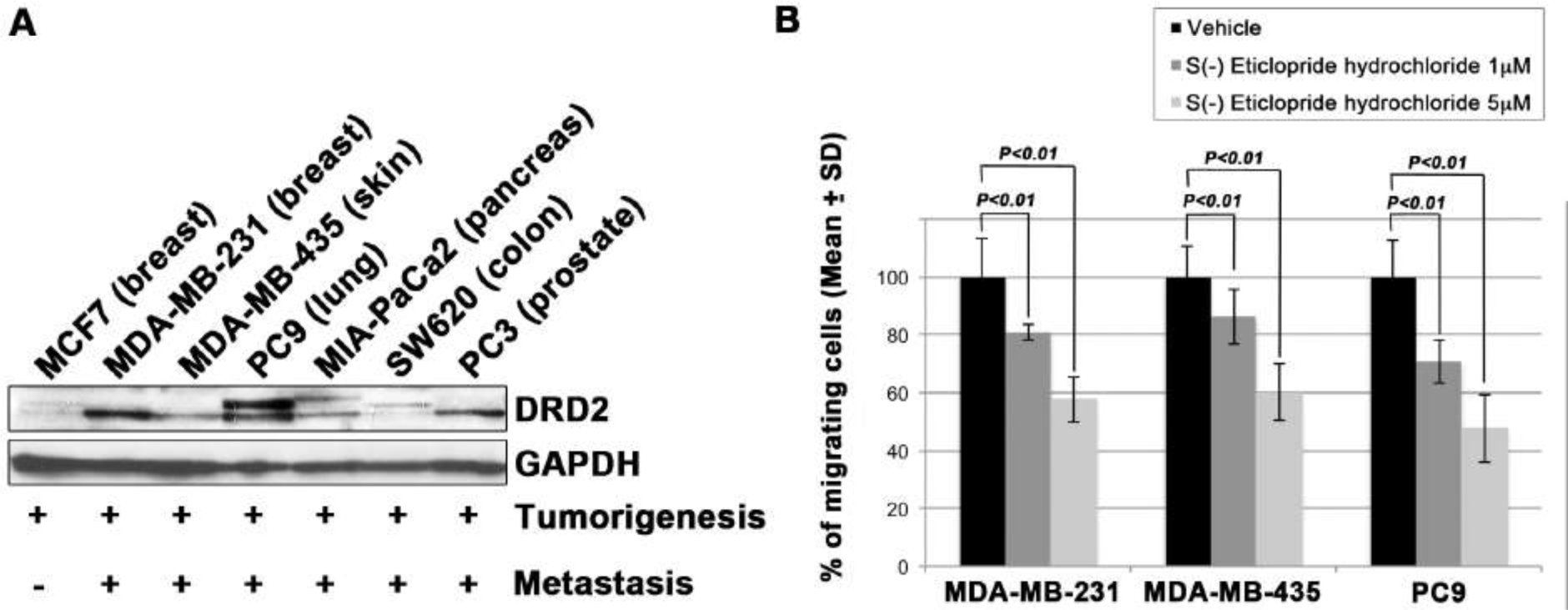
S (-) Eticlopride hydrochloride suppressed cell motility and invasion of human cancer cells. Related to Figure 2. (A) Western blot analysis of DRD2 levels in non-metastatic human cancer cell line, MCF7 (breast) and highly metastatic human cancer cell lines, MDA-MB-231 (breast), MDA-MB-435 (melanoma), MIA-PaCa2 (pancreas), PC3 (prostate) and SW620 (colon); GAPDH loading control is shown (bottom). (B) Effect of S (-) Eticlopride hydrochloride on cell motility and invasion of MBA-MB-231, MDA-MB-435, and PC9 cells. Either vehicle or pizotifen treated cells were subjected to Boyden chamber assays. Fetal bovine serum (1%v/v) was used as the chemoattractant in both assays. Each experiment was performed at least twice. Statistical analysis was determined by Student’s *t* test.

**Figure S3.**
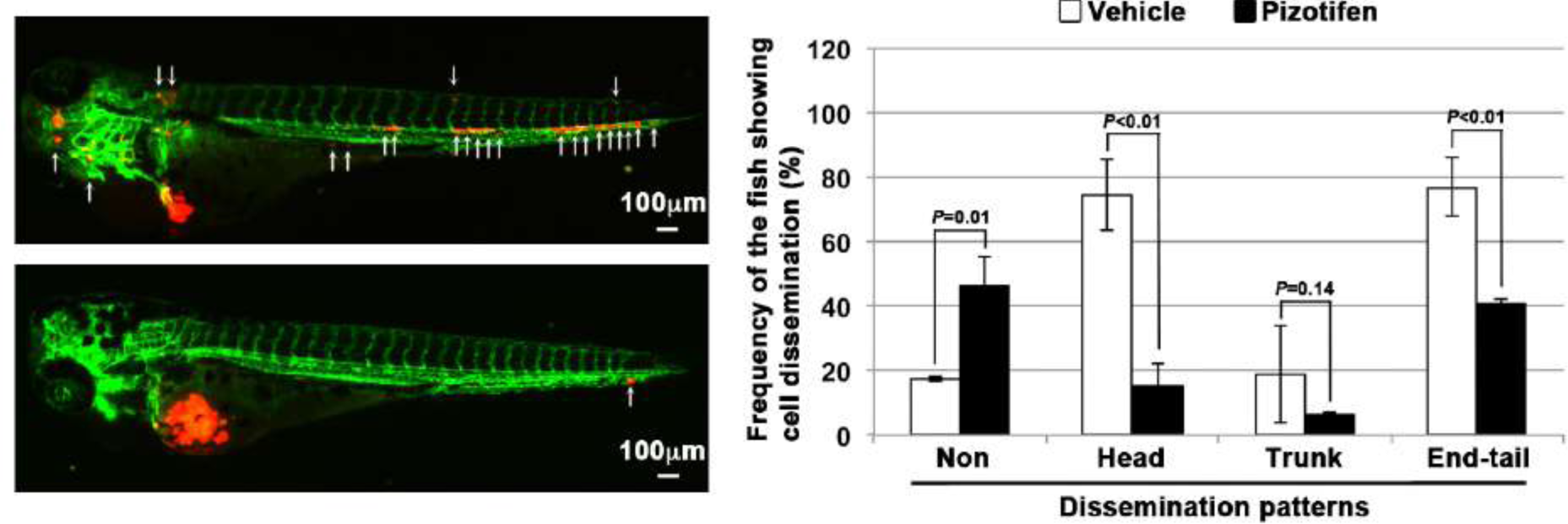
Pizotifen suppressed metastatic dissemination of human pancreatic cancer cells in a zebrafish xenotransplantation model. Related to Figure 2. Representative images of dissemination of MIA-PaCa2 cells in zebrafish xenotransplantation model. The fish were inoculated with MIA-PaCa2 cells, and treated with either vehicle (top left) or drug (lower left). White arrows heads indicate disseminated MIA-PaCa2 cells. The images were shown in 4x magnification. Scale bar, 100μm. The mean frequencies of the fish showing head, trunk, or end-tail dissemination were tabulated (right). Each value is indicated as the mean ± SEM of two independent experiments. Statistical analysis was determined by Student’s *t* test.

**Table S1.**
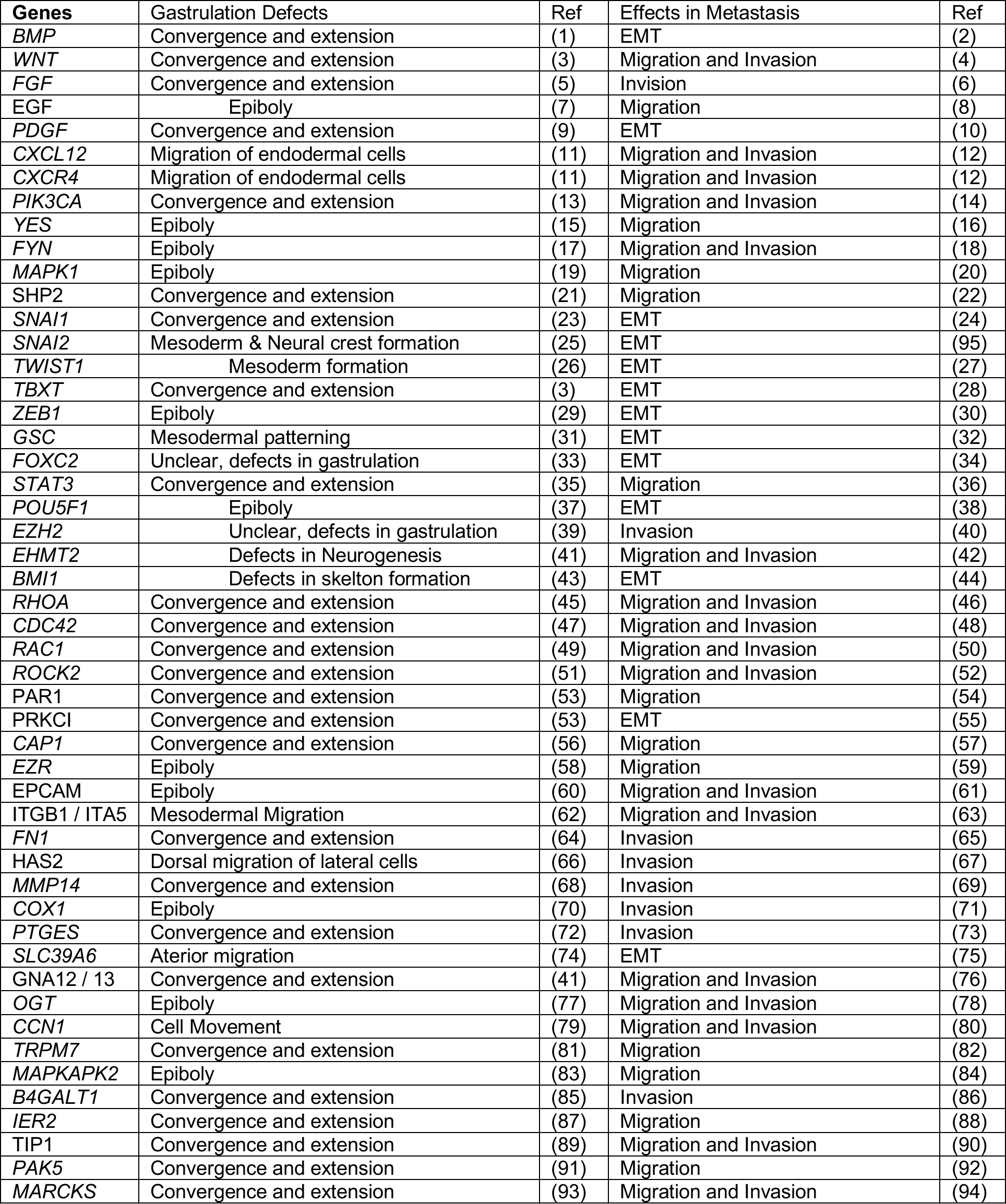
A list of the genes that are involved between gastrulation and metastasis progression. A list of the fifty genes that play essential role in governing both metastasis and gastrulation progression. The gastrulation defects in Xenopus or zebrafish that are induced by knockdown of each of these genes, were indicated. The molecular mechanism in metastasis that are inhibited by knockdown of each of the same genes, were indicated.

| Genes | Gastrulation Defects | Ref | Effects in Metastasis | Ref |
| --- | --- | --- | --- | --- |
| *BMP* | Convergence and extension | (1) | EMT | (2) |
| *WNT* | Convergence and extension | (3) | Migration and Invasion | (4) |
| *FGF* | Convergence and extension | (5) | Invasion | (6) |
| *EGF* | Epiboly | (7) | Migration | (8) |
| *PDGF* | Convergence and extension | (9) | EMT | (10) |
| *CXCL12* | Migration of endodermal cells | (11) | Migration and Invasion | (12) |
| *CXCR4* | Migration of endodermal cells | (11) | Migration and Invasion | (12) |
| *PIK3CA* | Convergence and extension | (13) | Migration and Invasion | (14) |
| *YES* | Epiboly | (15) | Migration | (16) |
| *FYN* | Epiboly | (17) | Migration and Invasion | (18) |
| *MAPK1* | Epiboly | (19) | Migration | (20) |
| *SHP2* | Convergence and extension | (21) | Migration | (22) |
| *SNAI1* | Convergence and extension | (23) | EMT | (24) |
| *SNAI2* | Mesoderm & Neural crest formation | (25) | EMT | (95) |
| *TWIST1* | Mesoderm formation | (26) | EMT | (27) |
| *TBXT* | Convergence and extension | (3) | EMT | (28) |
| *ZEB1* | Epiboly | (29) | EMT | (30) |
| *GSC* | Mesodermal patterning | (31) | EMT | (32) |
| *FOXC2* | Unclear, defects in gastrulation | (33) | EMT | (34) |
| *STAT3* | Convergence and extension | (35) | Migration | (36) |
| *POU5F1* | Epiboly | (37) | EMT | (38) |
| *EZH2* | Unclear, defects in gastrulation | (39) | Invasion | (40) |
| *EHMT2* | Defects in Neurogenesis | (41) | Migration and Invasion | (42) |
| *BMI1* | Defects in skelton formation | (43) | EMT | (44) |
| *RHOA* | Convergence and extension | (45) | Migration and Invasion | (46) |
| *CDC42* | Convergence and extension | (47) | Migration and Invasion | (48) |
| *RAC1* | Convergence and extension | (49) | Migration and Invasion | (50) |
| *ROCK2* | Convergence and extension | (51) | Migration and Invasion | (52) |
| *PAR1* | Convergence and extension | (53) | Migration | (54) |
| *PRKCI* | Convergence and extension | (53) | EMT | (55) |
| *CAP1* | Convergence and extension | (56) | Migration | (57) |
| *EZR* | Epiboly | (58) | Migration | (59) |
| *EPCAM* | Epiboly | (60) | Migration and Invasion | (61) |
| *ITGB1 / ITA5* | Mesodermal Migration | (62) | Migration and Invasion | (63) |
| *FN1* | Convergence and extension | (64) | Invasion | (65) |
| *HAS2* | Dorsal migration of lateral cells | (66) | Invasion | (67) |
| *MMP14* | Convergence and extension | (68) | Invasion | (69) |
| *COX1* | Epiboly | (70) | Invasion | (71) |
| *PTGES* | Convergence and extension | (72) | Invasion | (73) |
| *SLC39A6* | Aterior migration | (74) | EMT | (75) |
| *GNA12 / 13* | Convergence and extension | (41) | Migration and Invasion | (76) |
| *OGT* | Epiboly | (77) | Migration and Invasion | (78) |
| *CCN1* | Cell Movement | (79) | Migration and Invasion | (80) |
| *TRPM7* | Convergence and extension | (81) | Migration | (82) |
| *MAPKAPK2* | Epiboly | (83) | Migration | (84) |
| *B4GALT1* | Convergence and extension | (85) | Invasion | (86) |
| *IER2* | Convergence and extension | (87) | Migration | (88) |
| *TIP1* | Convergence and extension | (89) | Migration and Invasion | (90) |
| *PAK5* | Convergence and extension | (91) | Migration | (92) |
| *MARCKS* | Convergence and extension | (93) | Migration and Invasion | (94) |

**Table S2.**
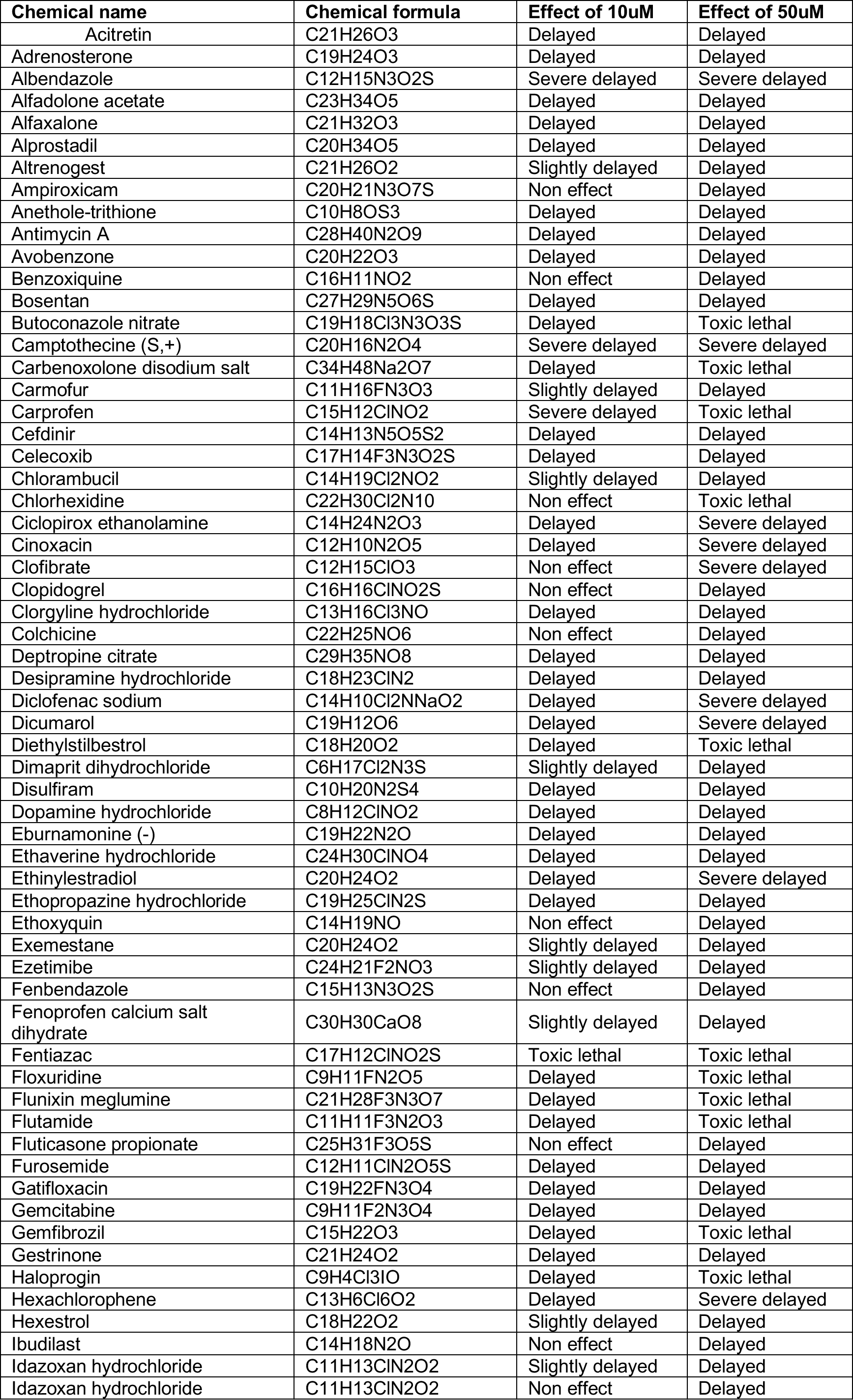
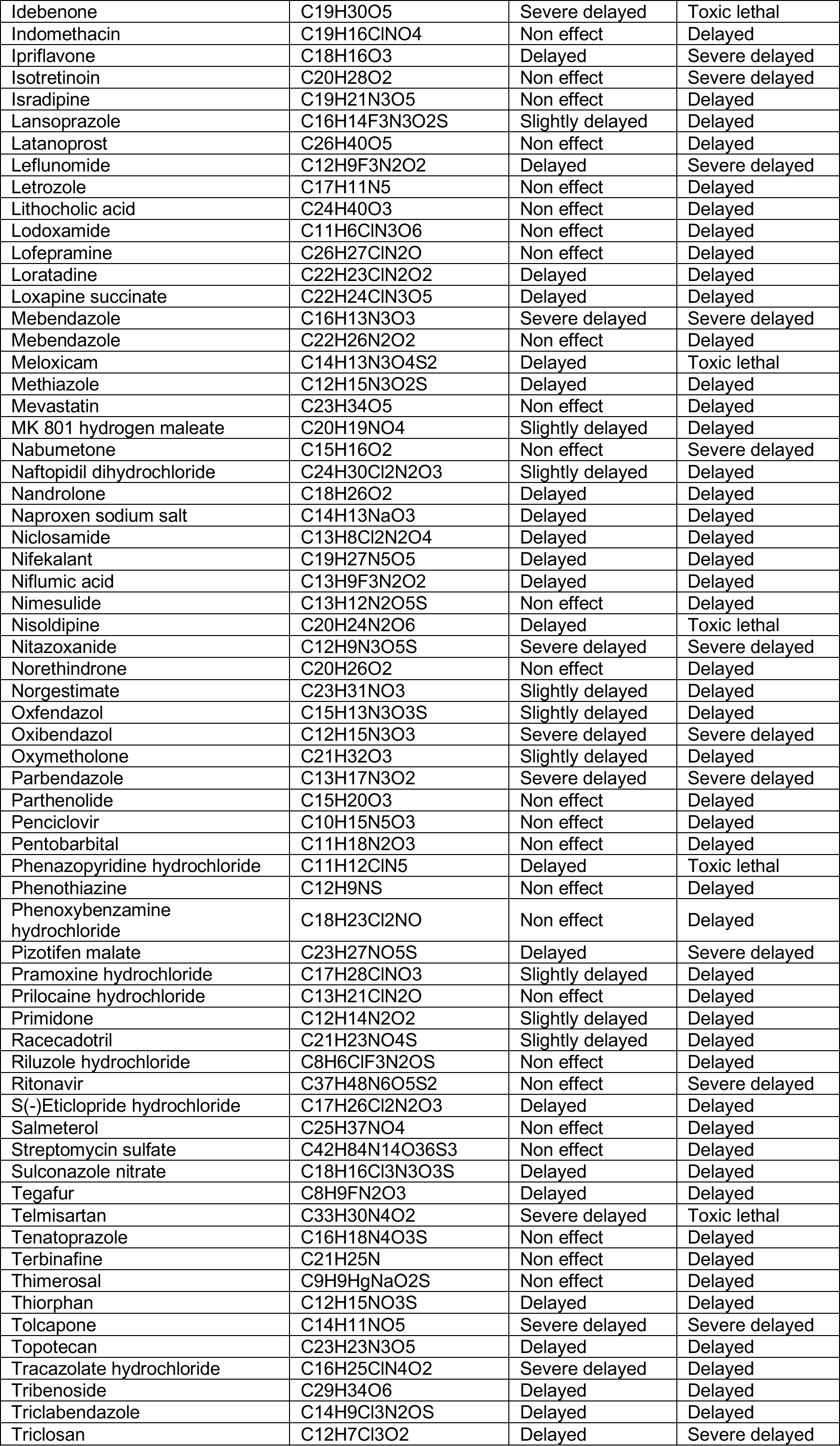
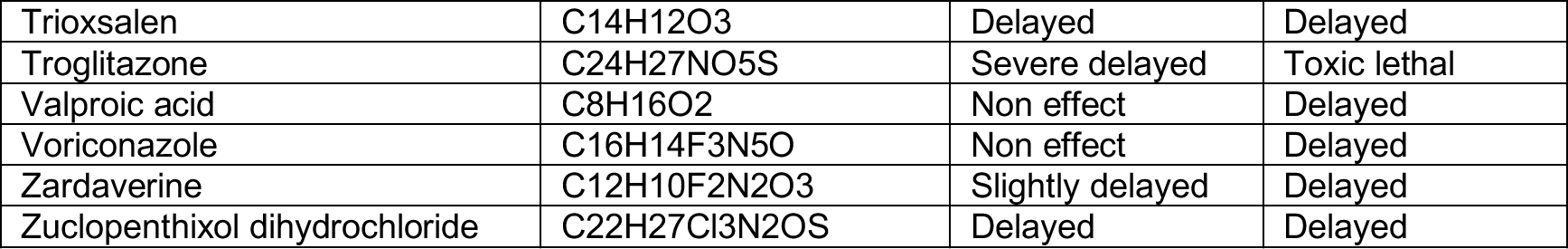
A list of the drugs that interfere with epiboly progression in zebrafish. Related to Figure 1. A list of positive “hit” drugs that interfered with epiboly progression. Gastrulation defects or status of each of the zebrafish embryos that were treated with either 10μM or 50μM concentrations, are indicated.

**Table S3.** Effects of pharmacological inhibition of HTR2C on metastatic dissemination of MDA-MB-231 cells in zebrafish xenografted models. Related to Figure 2D. The numbers and frequencies of the fish showing the dissemination patterns in vehicle or Pizotifen-treated group, were indicated. The fish showed both patterns of dissemination were redundantly counted in this analysis.

|  |  | Experiment #1 | Experiment #2 | Experiment #3 | Average of Experiments |
| --- | --- | --- | --- | --- | --- |
| Drug: Vehicle<br>Cell: MDA-MB-231 | Non-dissemination | 0%<br>n1=0/17 | 0%<br>n2=0/12 | 6.66%<br>n3=1/15 | 2.22±3.84% |
|  | Head | 58.82%<br>n1=10/17 | 91.66%<br>n2=11/12 | 6.66%<br>n3=1/15 | 72.38±17.15% |
|  | Trunk | 52.94%<br>n1=9/17 | 8.33%<br>n2=1/12 | 20%<br>n3=2/15 | 27.09±23.13% |
|  | End-tail | 100%<br>n1=17/17 | 100%<br>n2=12/12 | 86.66%<br>n3=13/15 | 95.55±7.69% |
| Drug: Pizotifen<br>Cell: MDA-MB-231 | Non-dissemination | 55%<br>n1=11/20 | 31.57%<br>n2=6/19 | 45.45 %<br>n3=10/22 | 44.01±11.77% |
|  | Head | 5%<br>n1=1/20 | 31.57%<br>n2=6/19 | 18.18%<br>n3=4/22 | 18.25±13.28% |
|  | Trunk | 5%<br>n1=1/20 | 10.52%<br>n2=2/19 | 4.45%<br>n3=1/22 | 6.69±3.32% |
|  | End-tail | 45%<br>n1=9/20 | 57.89%<br>n2=11/19 | 50%<br>n3=11/22 | 50.96±6.50% |

**Table S4.** Effects of pharmacological inhibition of HTR2C on metastatic dissemination of Mia-PaCa2 cells in zebrafish xenografted models. Related to Figure S3. The numbers and frequencies of the fish showing the dissemination patterns in vehicle or Pizotifen-treated group, were indicated. The fish showed both patterns of dissemination were redundantly counted in this analysis.

|  |  | Experiment #1 | Experiment #2 | Average of Experiments |
| --- | --- | --- | --- | --- |
| Drug: Vehicle<br>Cell: MIA-PaCa2 | Non-dissemination | 17.64%<br>n1=3/17 | 16.66%<br>n2=2/12 | 17.15+0.69% |
|  | Head | 82.35%<br>n1=14/17 | 66.66%<br>n2=8/12 | 74.50+11.09% |
|  | Trunk | 29.41%<br>n1=5/17 | 8.33%<br>n2=1/12 | 18.87+14.90% |
|  | End-tail | 70.58%<br>n1=12/17 | 83.33%<br>n2=10/17 | 76.96%+9.01 |
| Drug: Pizotifen<br>Cell: MIA-PaCa2 | Non-dissemination | 40%<br>n1=4/10 | 52.63%<br>n2=10/19 | 46.31+8.93% |
|  | Head | 20%<br>n1=2/10 | 10.52%<br>n2=2/19 | 15.26+6.69% |
|  | Trunk | 10%<br>n1=1/10 | 5.26%<br>n2=1/19 | 7.63+3.34% |
|  | End-tail | 40%<br>n1=4/10 | 42.05%<br>n2=8/19 | 41.4+1.48% |

**Table S5.** Effects of genetic inhibition of HTR2C on metastatic dissemination of MDA-MB-231 cells in zebrafish xenografted models. Related to Figure 2E. The numbers and frequencies of the fish showing the dissemination patterns in the zebrafish that were inoculated with either shLacZ or shHTR2C MDA-MB-231 cells, were indicated. The fish showed both patterns of dissemination were redundantly counted in this analysis.

|  |  | Experiment #1 | Experiment #2 | Average of Experiments |
| --- | --- | --- | --- | --- |
| shLacZ | Non-dissemination | 0%<br>n1=0/10 | 0%<br>n2=0/10 | 0% |
|  | Head | 60%<br>n1=6/10 | 100%<br>n2=10/10 | 80<br>±28.28% |
|  | Trunk | 30%<br>n1=3/10 | 10%<br>n2=1/10 | 20±14.14% |
|  | End-tail | 80%<br>n1=8/10 | 100%<br>n2=10/10 | 90%±14.14 |
| shHTR2C | Non-dissemination | 80%<br>n1=12/15 | 76.84%<br>n2=14/19 | 76.84±4.46<br>% |
|  | Head | 6.66%<br>n1=1/15 | 15.78%<br>n2=3/19 | 11.22±6.45% |
|  | Trunk | 6.66%<br>n1=1/15 | 5.26%<br>n2=1/19 | 5.96±0.99% |
|  | End-tail | 20%<br>n1=3/15 | 26.31%<br>n2=5/19 | 23.15±4.46% |

**Table S6.** Effects of HTR2C overexpression on metastatic dissemination of MCF7 cells in zebrafish xenografted models. Related to Figure 4E. The numbers and frequencies of the fish showing the dissemination patterns in the zebrafish that were inoculated with MCF7 cells expressing either VC or HTR2C, were indicated. The fish showed both patterns of dissemination were redundantly counted in this analysis.

|  |  | Experiment #1 | Experiment #2 | Average of Experiments |
| --- | --- | --- | --- | --- |
| VC | Non-dissemination | 46.15%<br>n1=6/13 | 40%<br>n2=4/10 | 43.07±4.35% |
|  | Head | 46.15%<br>n1=6/13 | 20%<br>n2=2/10 | 33.07±18.49% |
|  | Trunk | 0%<br>n1=0/13 | 0%<br>n2=0/10 | 0% |
|  | End-tail | 53.84%<br>n1=7/13 | 60%<br>n2=6/10 | 56.92±4.35 |
| HTR2C | Non-dissemination | 0%<br>n1=0/14 | 0%<br>n2=0/15 | 0% |
|  | Head | 100%<br>n1=14/14 | 93.33%<br>n2=14/15 | 96.66±4.71% |
|  | Trunk | 64.28%<br>n1=9/14 | 73.33%<br>n2=11/15 | 68.80±6.39% |
|  | End-tail | 85.71%<br>n1=12/14 | 93.33%<br>n2=14/15 | 89.52±5.38% |

## Source data and legend for source data

**Figure 2a_HRT2C_source data.**
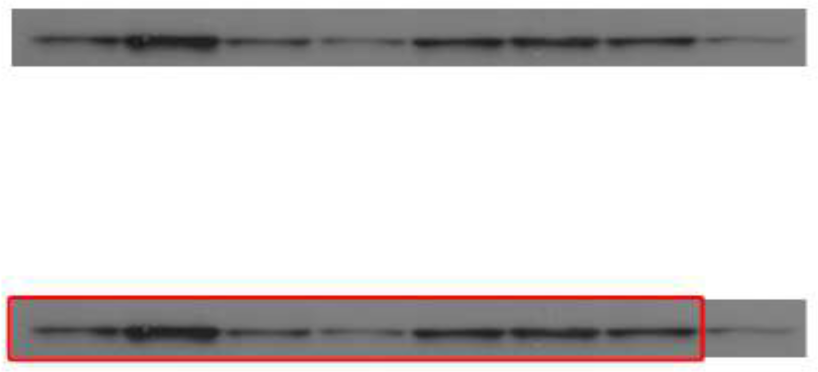
Western blot analysis of HRT2C protein levels in non-metastatic human cancer cell line, MCF7 (breast) and highly metastatic human cancer cell lines, MDA-MB-231 (breast), MDA-MB-435 (melanoma), MIA-PaCa2 (pancreas), SW620 (colon) and PC3 (prostate).

**Figure 2a_GAPDH_source data.**
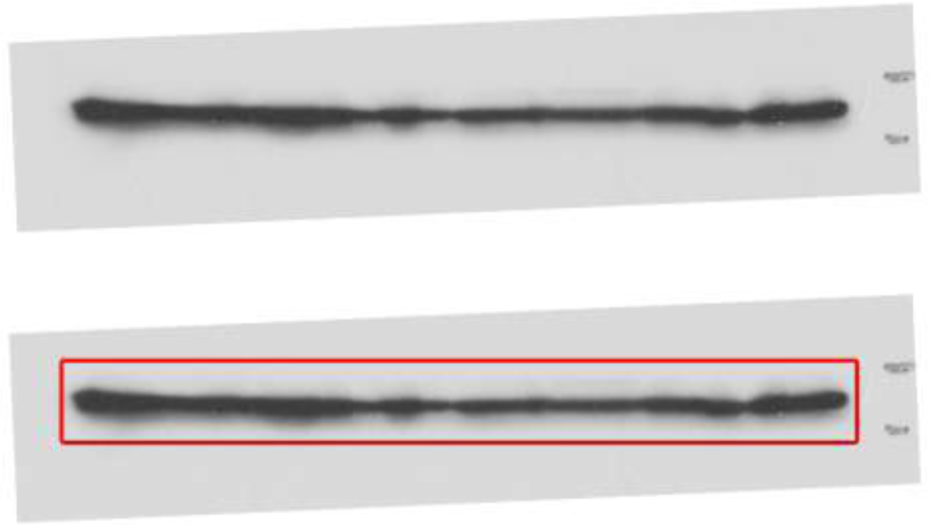
Western blot analysis of GAPDH protein levels in non-metastatic human cancer cell line, MCF7 (breast) and highly metastatic human cancer cell lines, MDA-MB-231 (breast), MDA-MB-435 (melanoma), MIA-PaCa2 (pancreas), SW620 (colon) and PC3 (prostate).

**Figure 4C_E-cadherin in MCF7_ source data.**
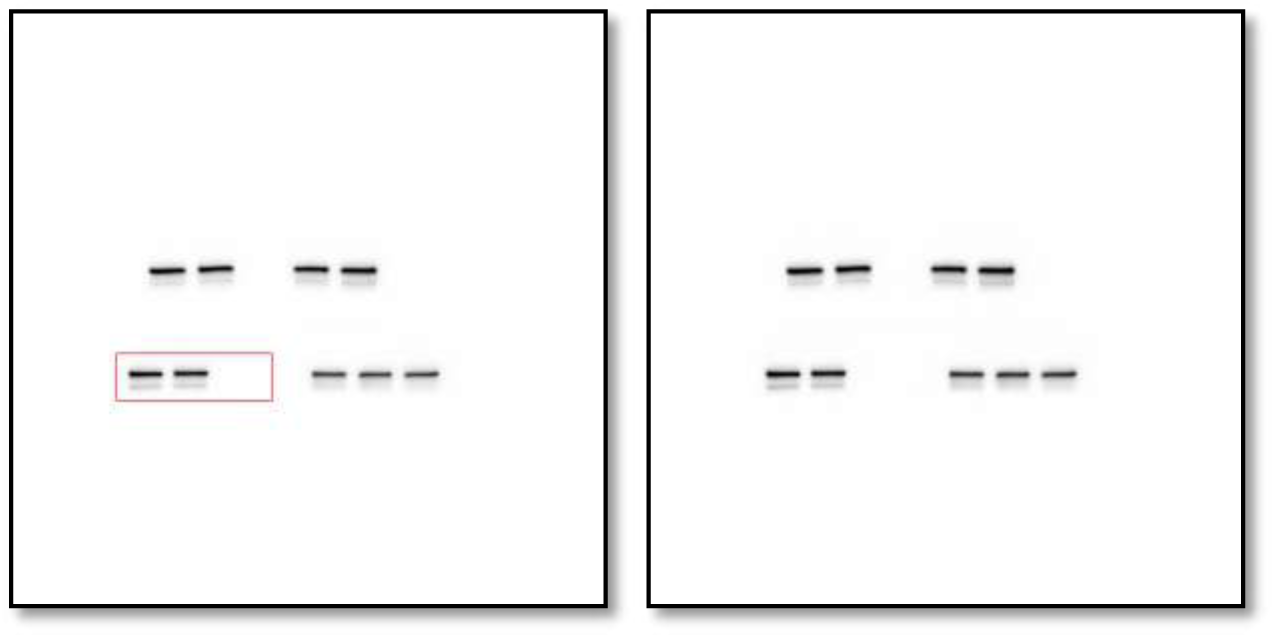
Western blot analysis of E-cadherin protein levels in MCF7 cells expressing either the control vector or HTR2C.

**Figure 4C_EpCAM in MCF7_ source data.**
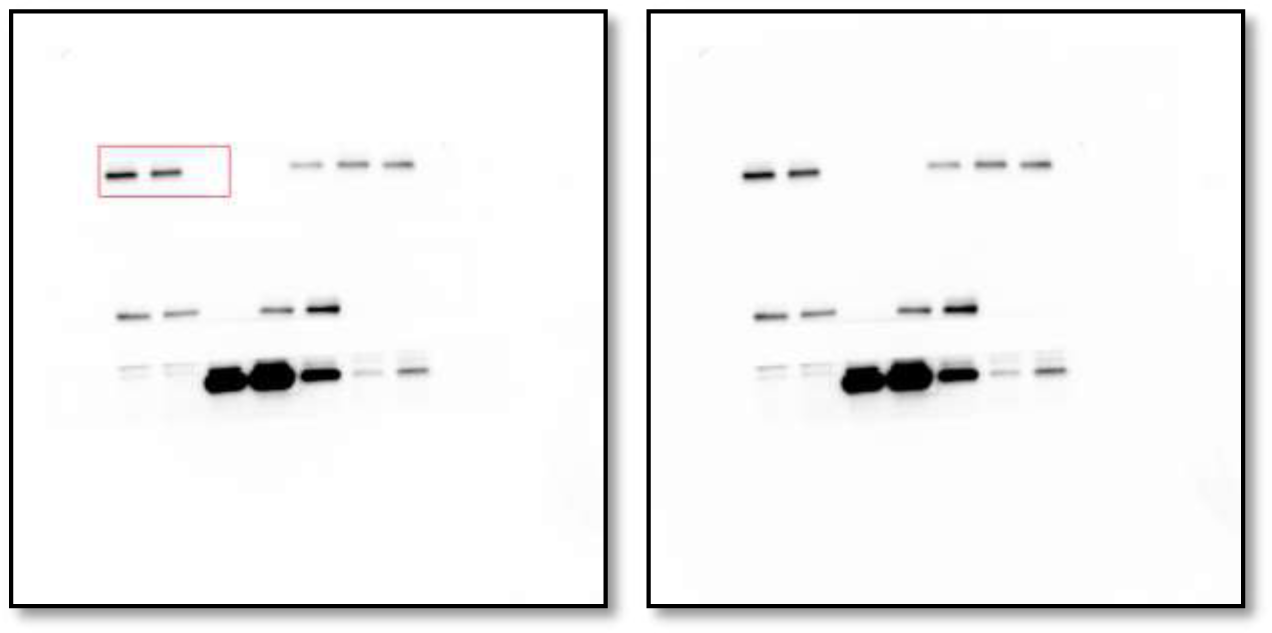
Western blot analysis of EpCAM protein levels in MCF7 cells expressing either the control vector or HTR2C.

**Figure 4C_Vimentin in MCF7_ source data.**
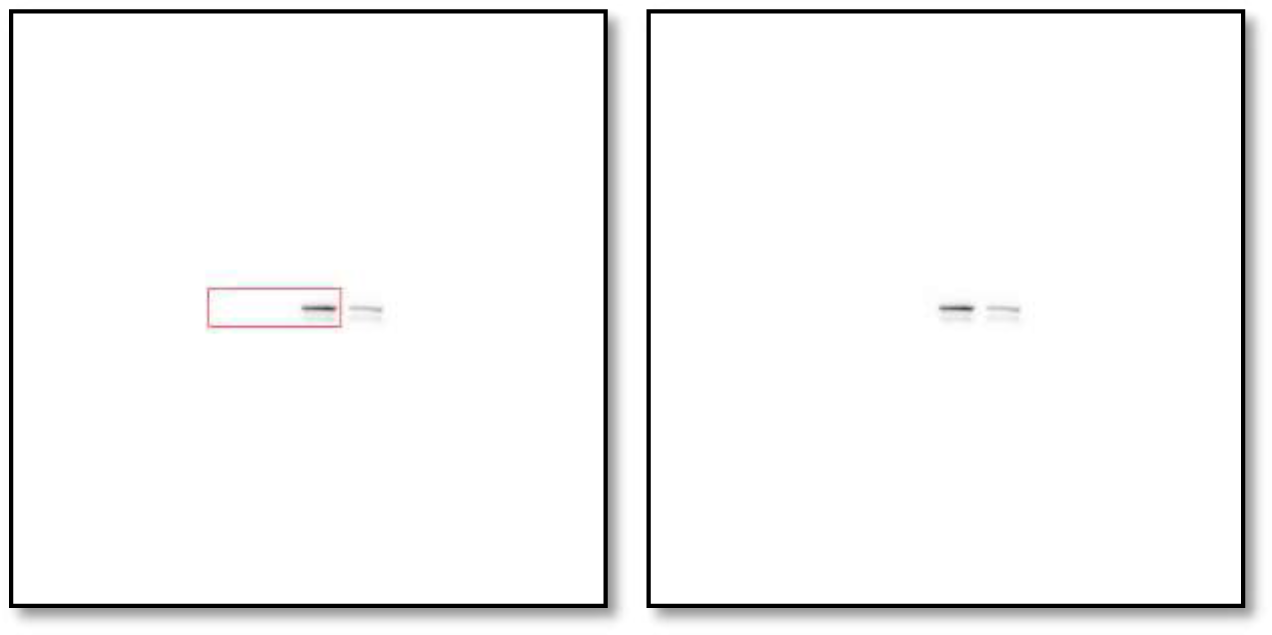
Western blot analysis of Vimentin protein levels in MCF7 cells expressing either the control vector or HTR2C.

**Figure 4C_N-cadherin in MCF7_ source data.**
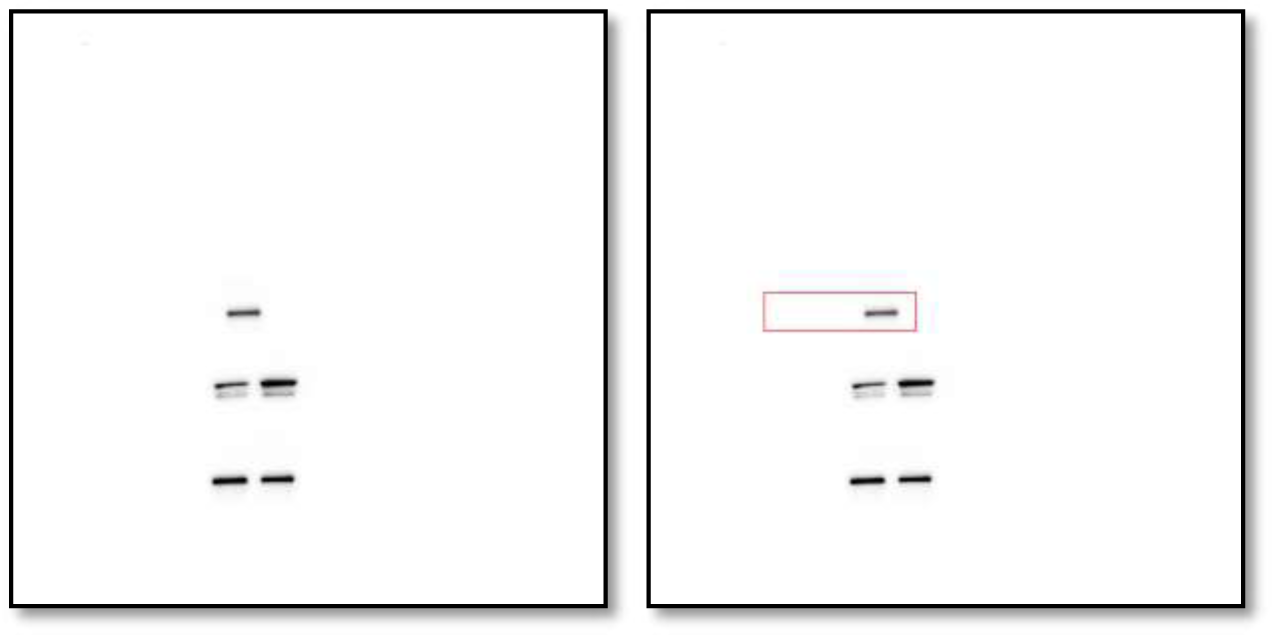
Western blot analysis of N-cadherin protein levels in MCF7 cells expressing either the control vector or HTR2C.

**Figure 4C_Zeb1 in MCF7_ source data.**
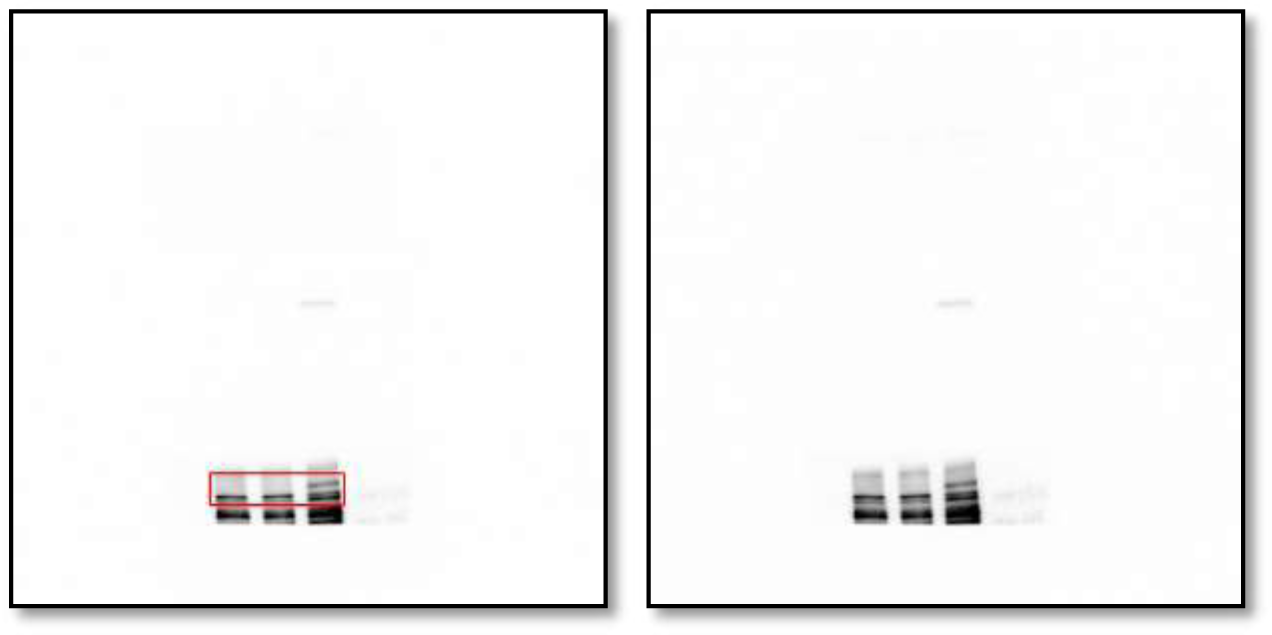
Western blot analysis of Zeb1 protein levels in MCF7 cells expressing either the control vector or HTR2C.

**Figure 4C_HRT2C in MCF7_ source data.**
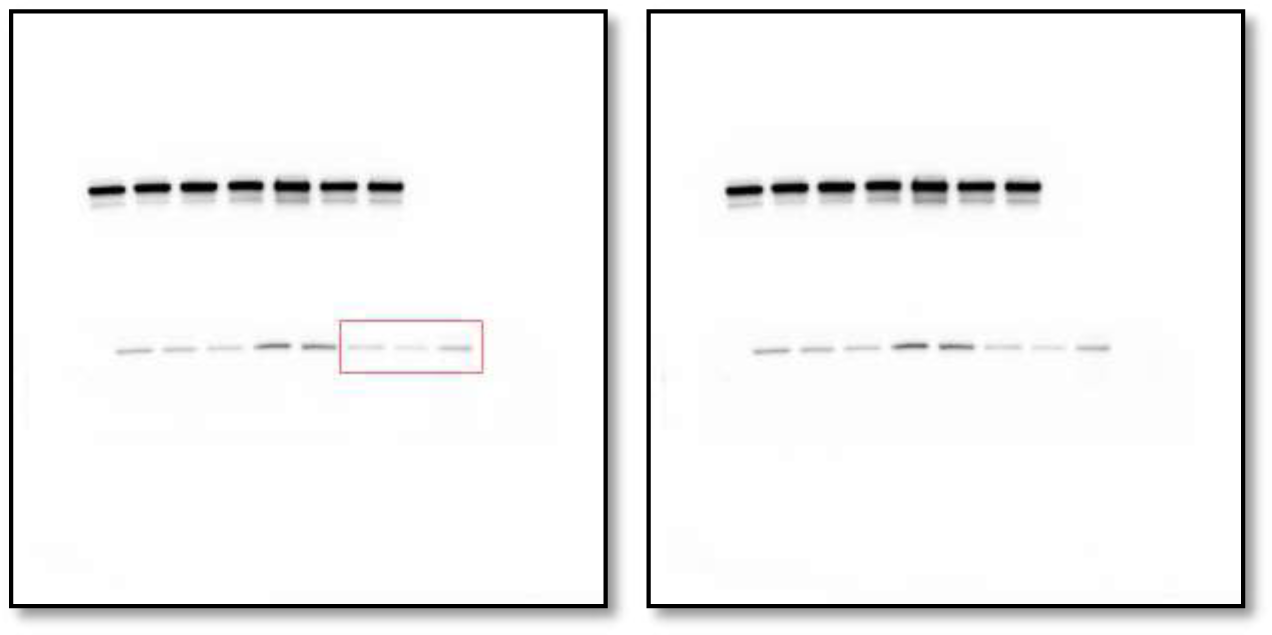
Western blot analysis of HRT2C protein levels in MCF7 cells expressing either the control vector or HTR2C.

**Figure 4C_GAPDH in MCF7_source data.**
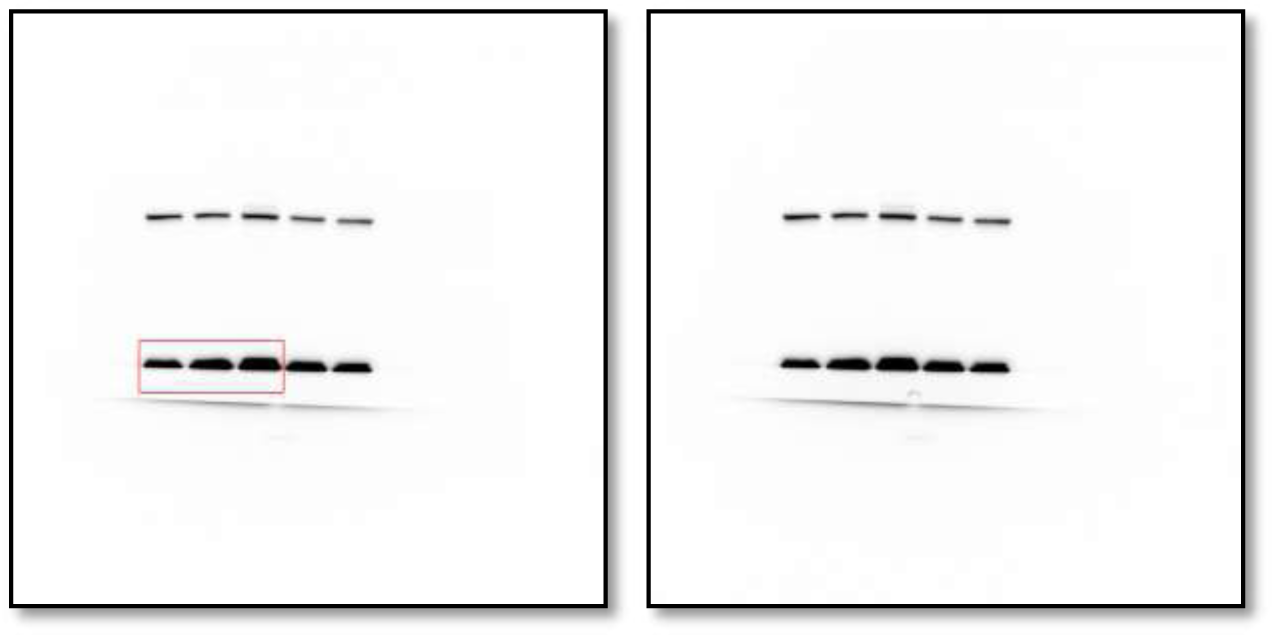
Western blot analysis of GAPDH protein levels in MCF7 cells expressing either the control vector or HTR2C.

**Figure 4C_E-cadherin in HaCaT_source data.**
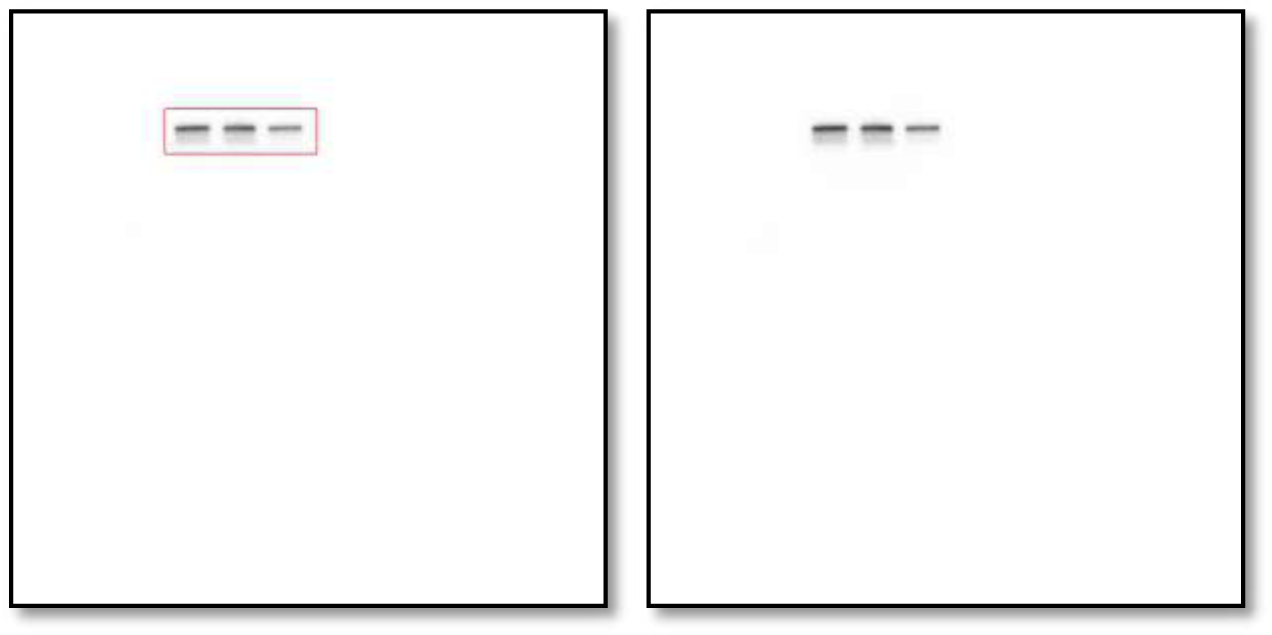
Western blot analysis of E-cadherin protein levels in HaCaT cells expressing either the control vector or HTR2C.

**Figure 4C_EpCAM in HaCaT_source data.**
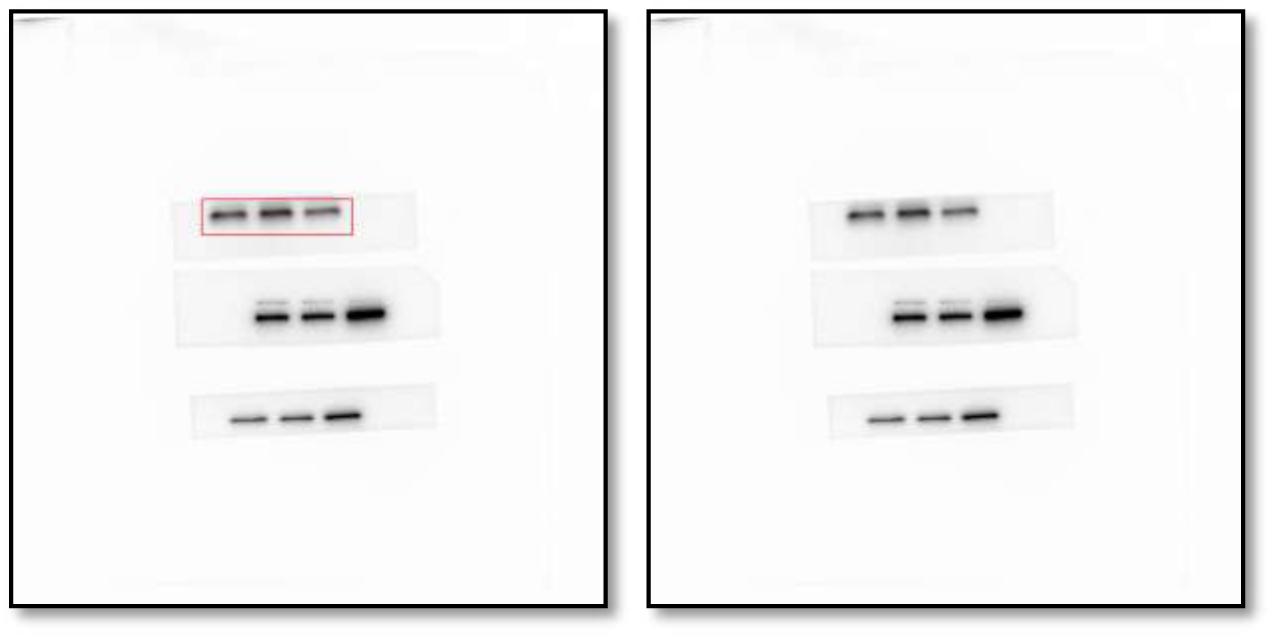
Western blot analysis of EpCAM protein levels in HaCaT cells expressing either the control vector or HTR2C.

**Figure 4C_Vimentin in HaCaT_source data.**
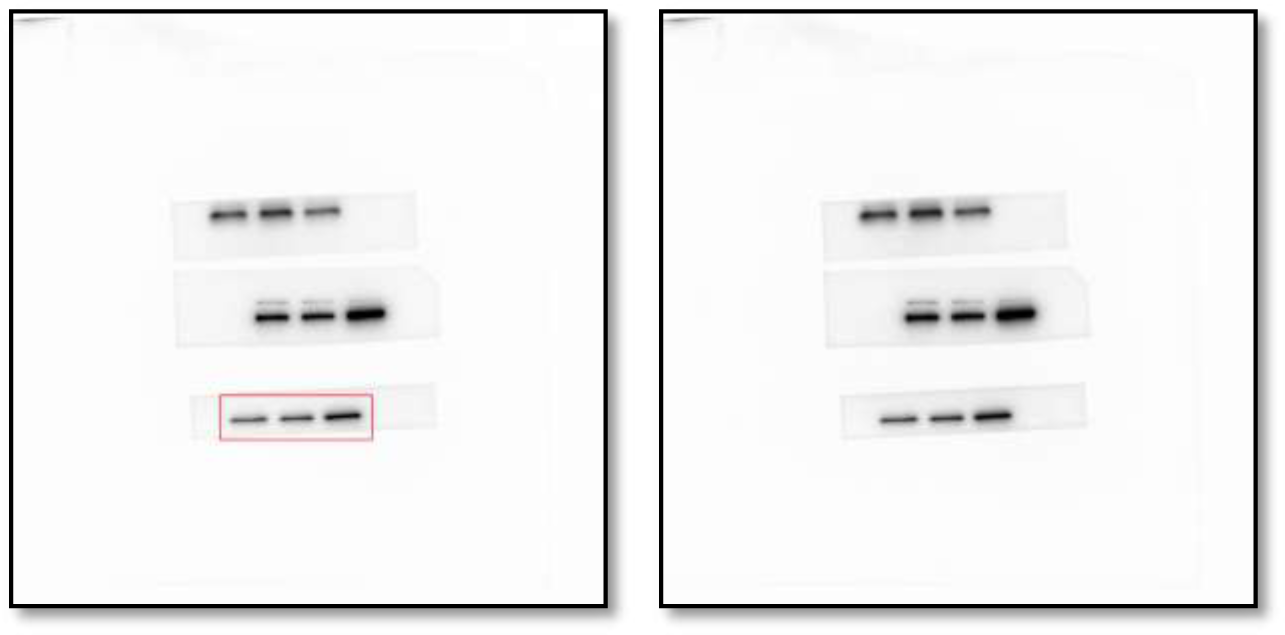
Western blot analysis of Vimentin protein levels in HaCaT cells expressing either the control vector or HTR2C.

**Figure 4C_N-cadherin in HaCaT_source data.**
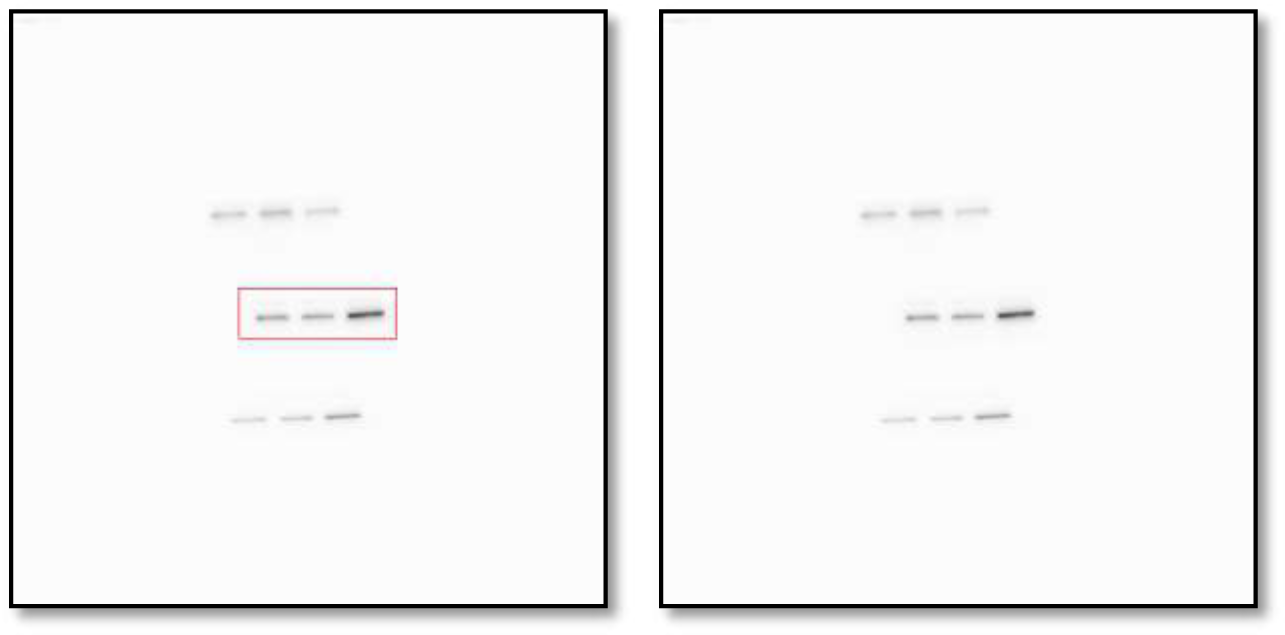
Western blot analysis of N-cadherin protein levels in HaCaT cells expressing either the control vector or HTR2C.

**Figure 4C_Zeb1 in HaCaT source data.**
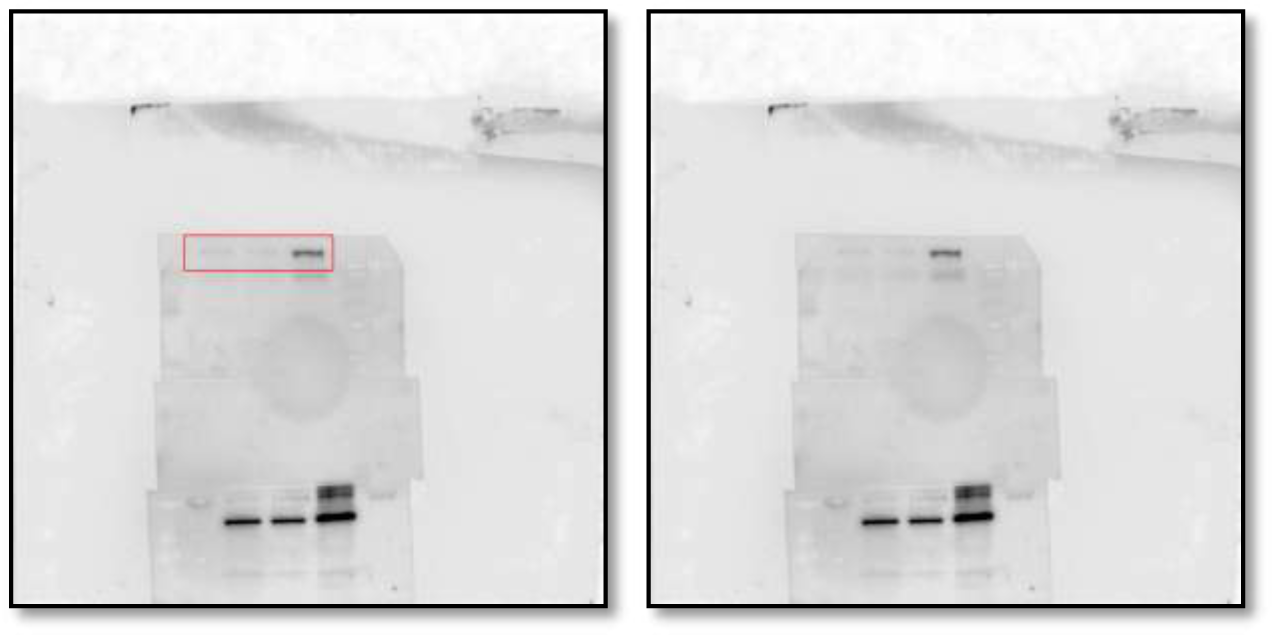
Western blot analysis of zeb1 protein levels in HaCaT cells expressing either the control vector or HTR2C.

**Figure 4C_HRT2C in HaCaT_source data.**
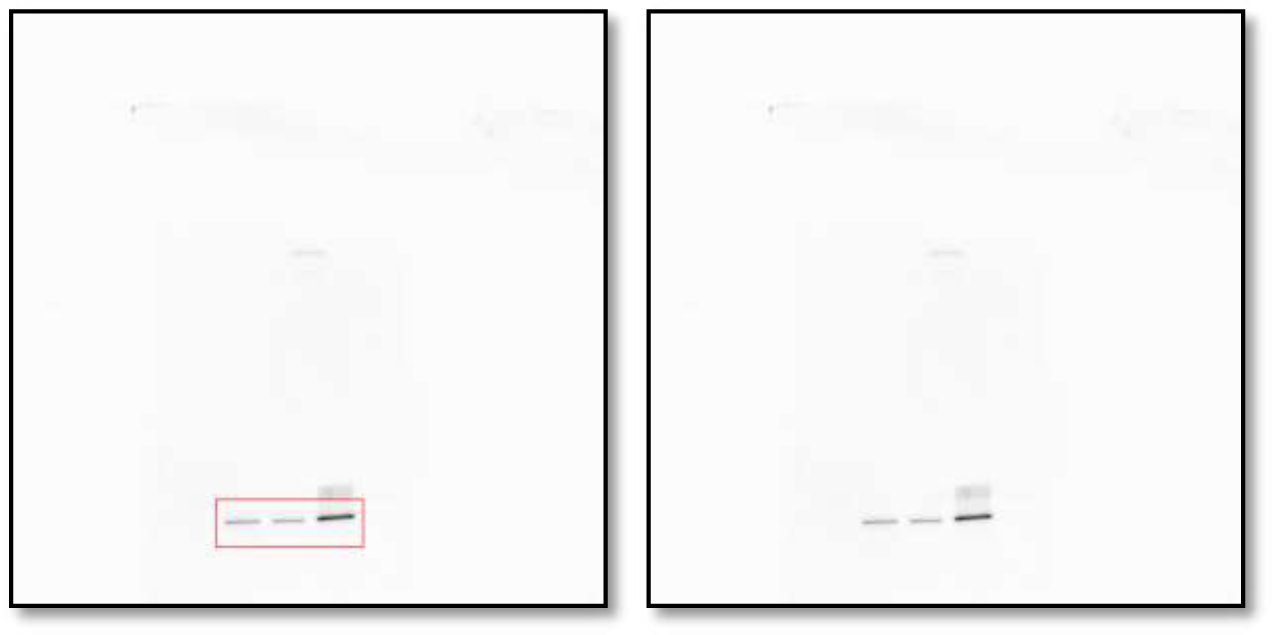
Western blot analysis of HRT2C protein levels in HaCaT cells expressing either the control vector or HTR2C.

**Figure 4C_GAPDH in HaCaT_source data.**
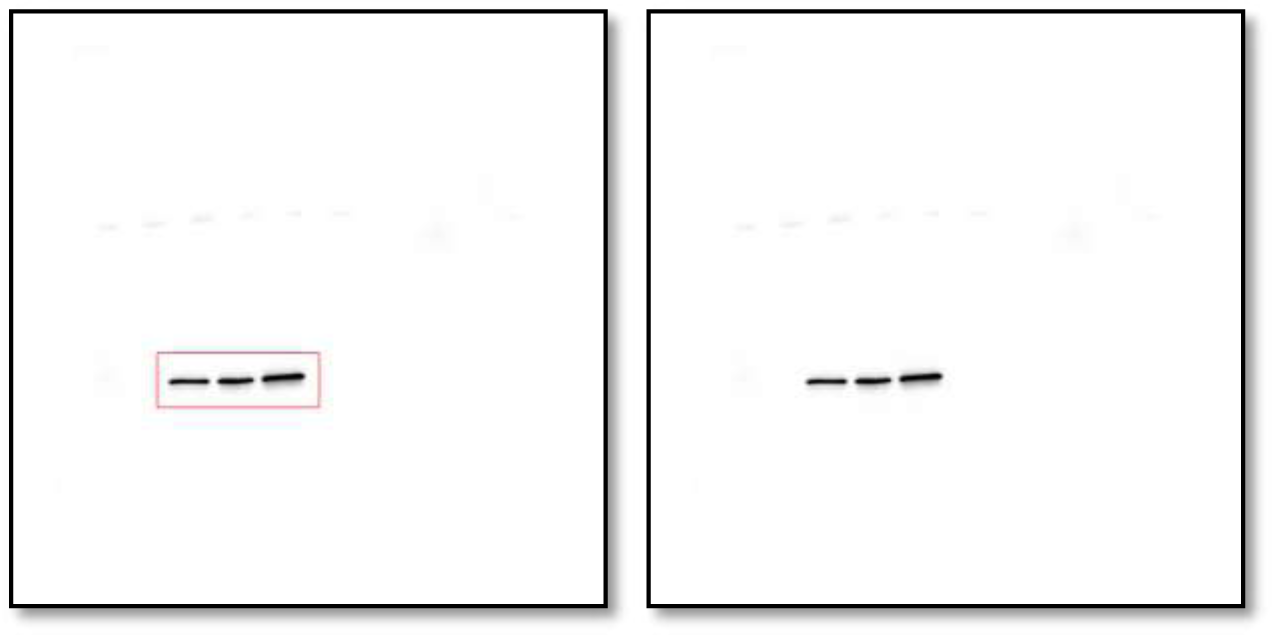
Western blot analysis of GAPDH protein levels in HaCaT cells expressing either the control vector or HTR2C.

**Figure 5C_E-cadherin_source data.**
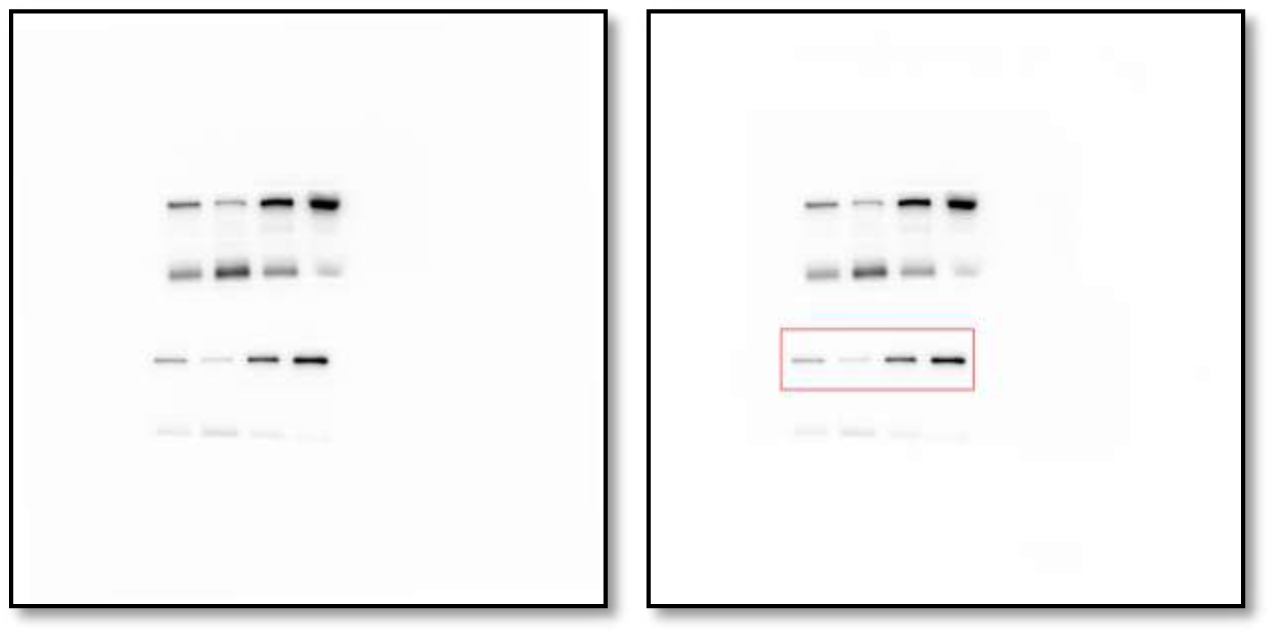
Western blot analysis of E-cadherin protein levels in whole cell lysate of 4T1 primary tumors from either vehicle or Pizotifen-treated mice.

**Figure 5C_Zeb1 _source data.**
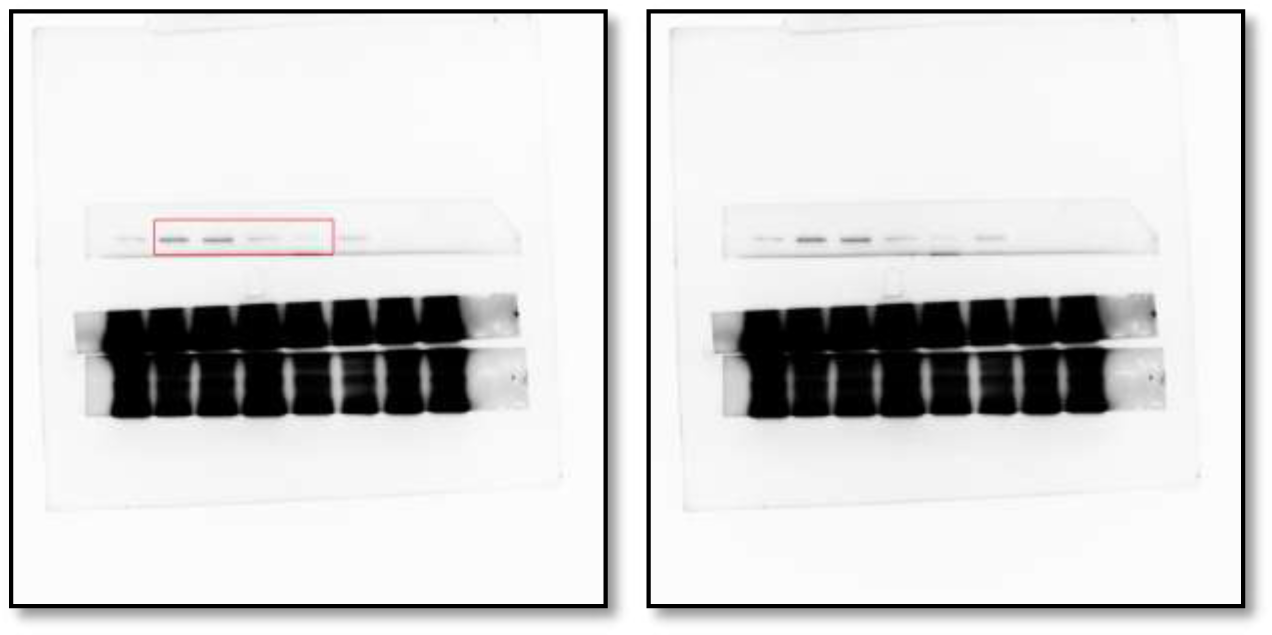
Western blot analysis of Zeb1 protein levels in whole cell lysate of 4T1 primary tumors from either vehicle or Pizotifen-treated mice.

**Figure 5C_ Phosphorylation of serine-9 in GSK3β_source data.**
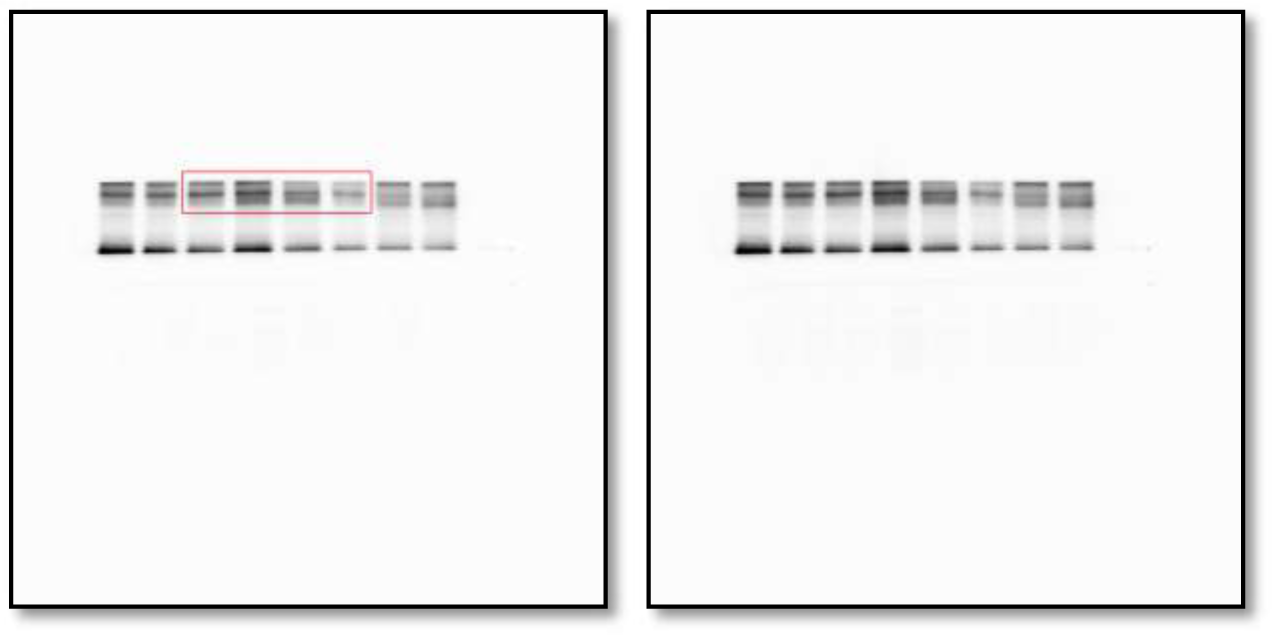
Western blot analysis of the protein levels of phosphorylation of serine-9 in GSK3β in whole cell lysate of 4T1 primary tumors from either vehicle or Pizotifen-treated mice.

**Figure 5C_ GSK3β_source data.**
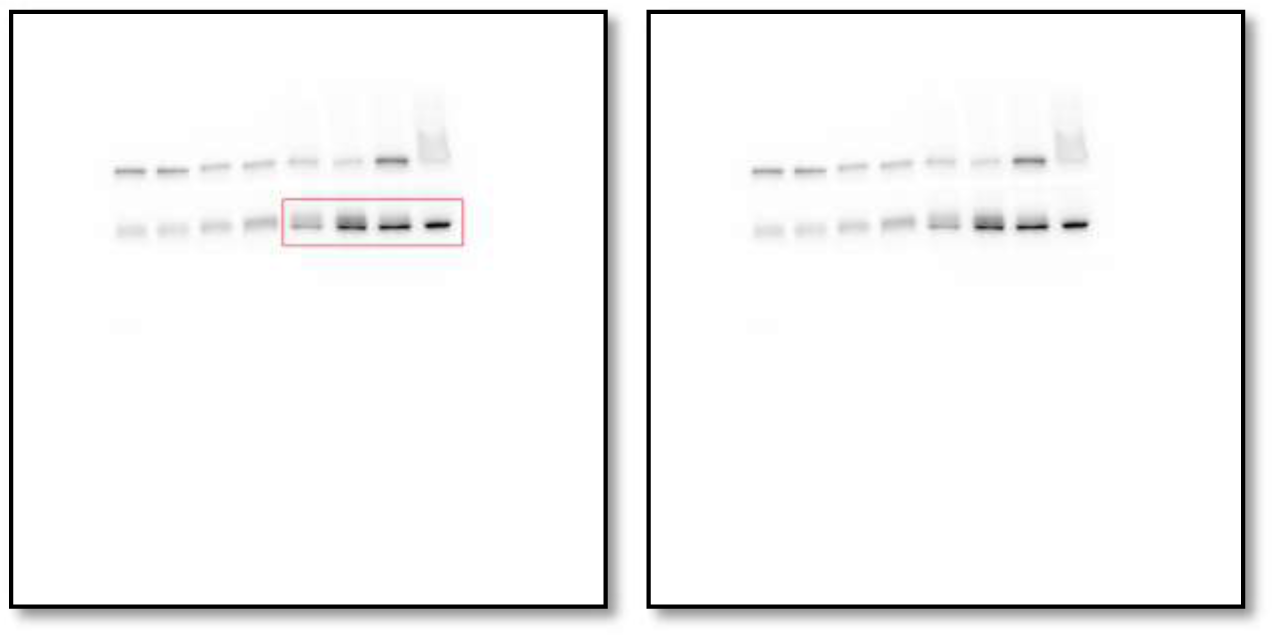
Western blot analysis of GSK3β protein levels in whole cell lysate of 4T1 primary tumors from either vehicle or Pizotifen-treated mice.

**Figure 5C_Luciferase_source data.**
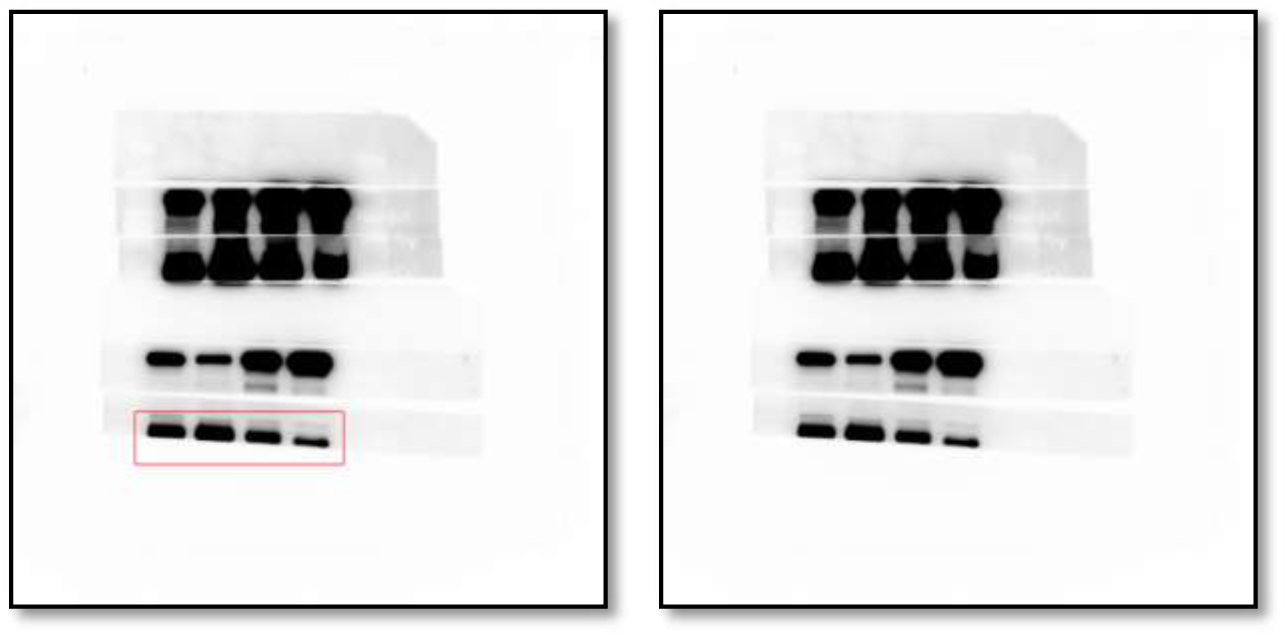
Western blot analysis of Luciferase protein levels in whole cell lysate of 4T1 primary tumors from either vehicle or Pizotifen-treated mice.

**Figure 5C_β-catenin in the nucleus_source data.**
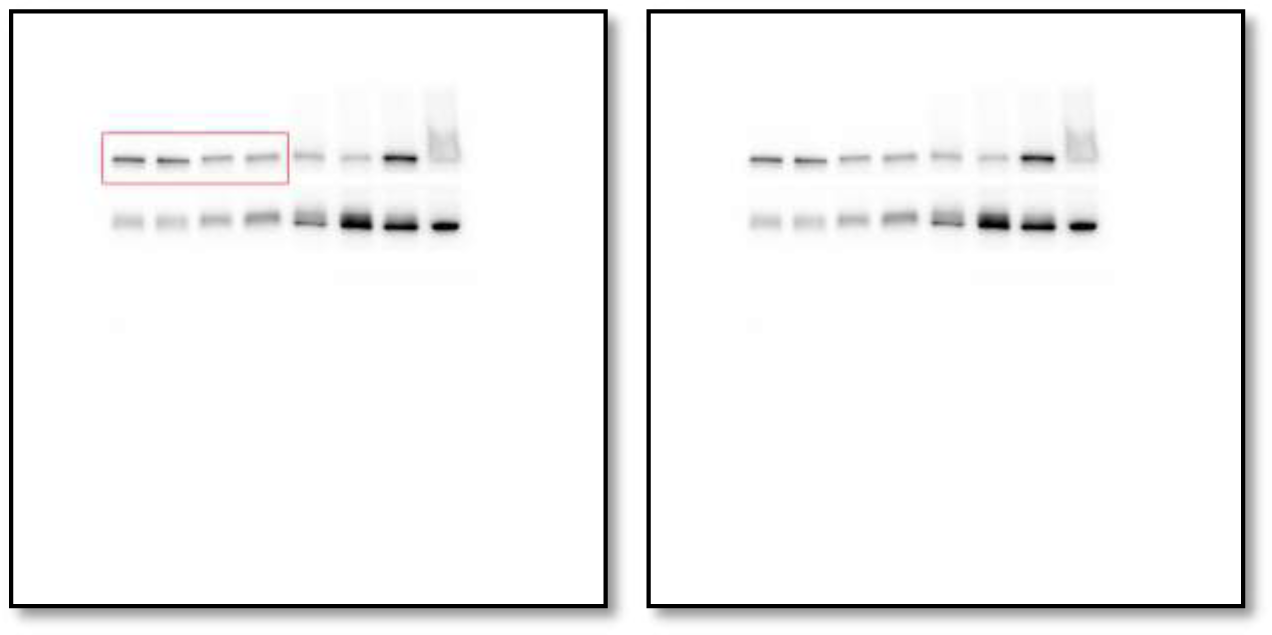
Western blot analysis of β-catenin protein levels in the nucleus of 4T1 primary tumors from either vehicle or Pizotifen-treated mice.

**Figure 5C_Histone H3 in the nucleus_source data.**
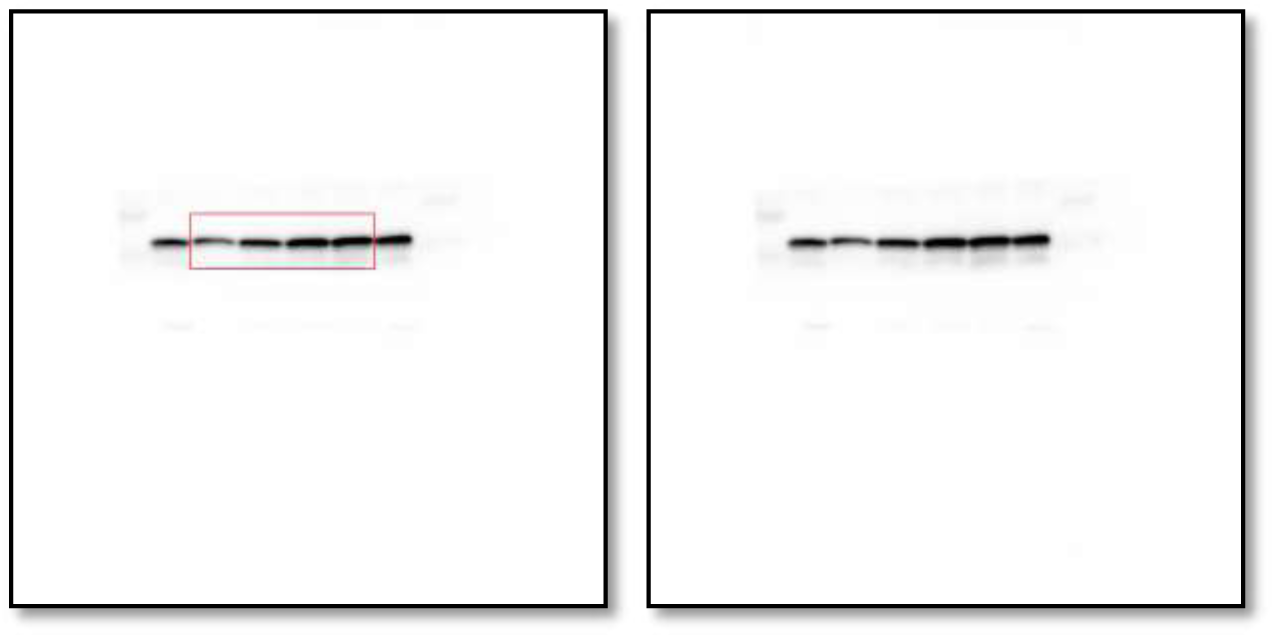
Western blot analysis of Histone H3 protein levels in the nucleus of 4T1 primary tumors from either vehicle or Pizotifen-treated mice.

**Figure 5C_β-catenin in the cytoplasm_source data.**
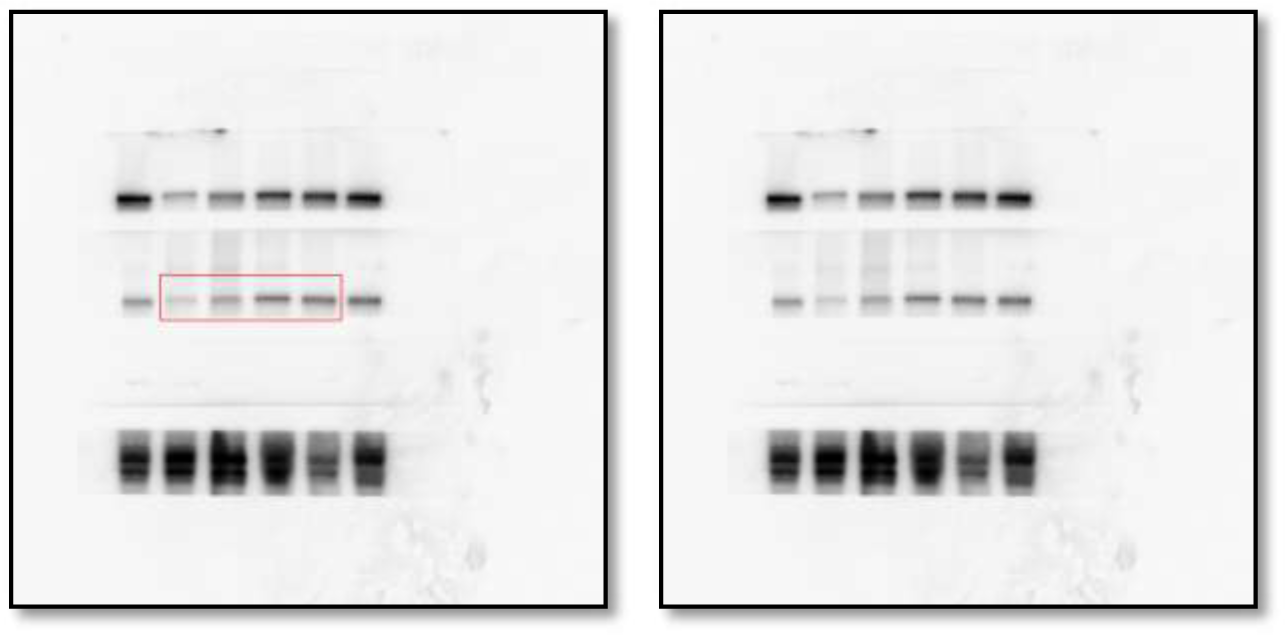
Western blot analysis of β-catenin protein levels in the cytoplasm of 4T1 primary tumors from either vehicle or Pizotifen-treated mice.

**Figure 5C_β-tubulin in the cytoplasm_source data.**
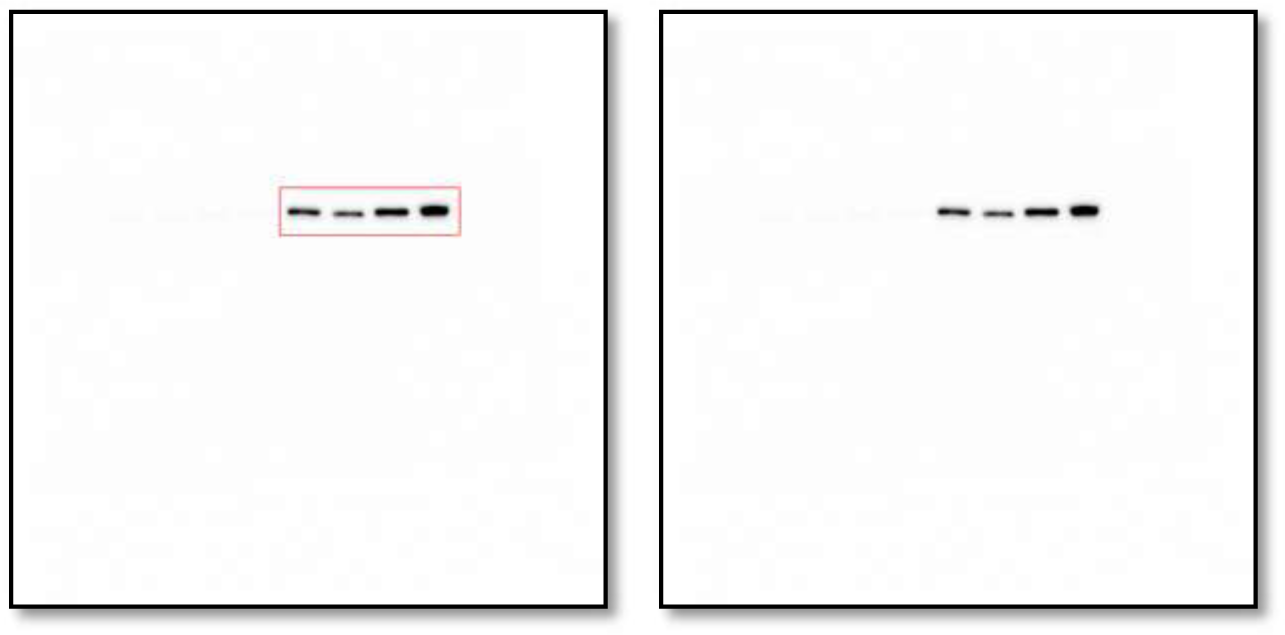
Western blot analysis of β-tubulin protein levels in the cytoplasm of 4T1 primary tumors from either vehicle or Pizotifen-treated mice.

**Figure 5D_E-cadherin_source data.**
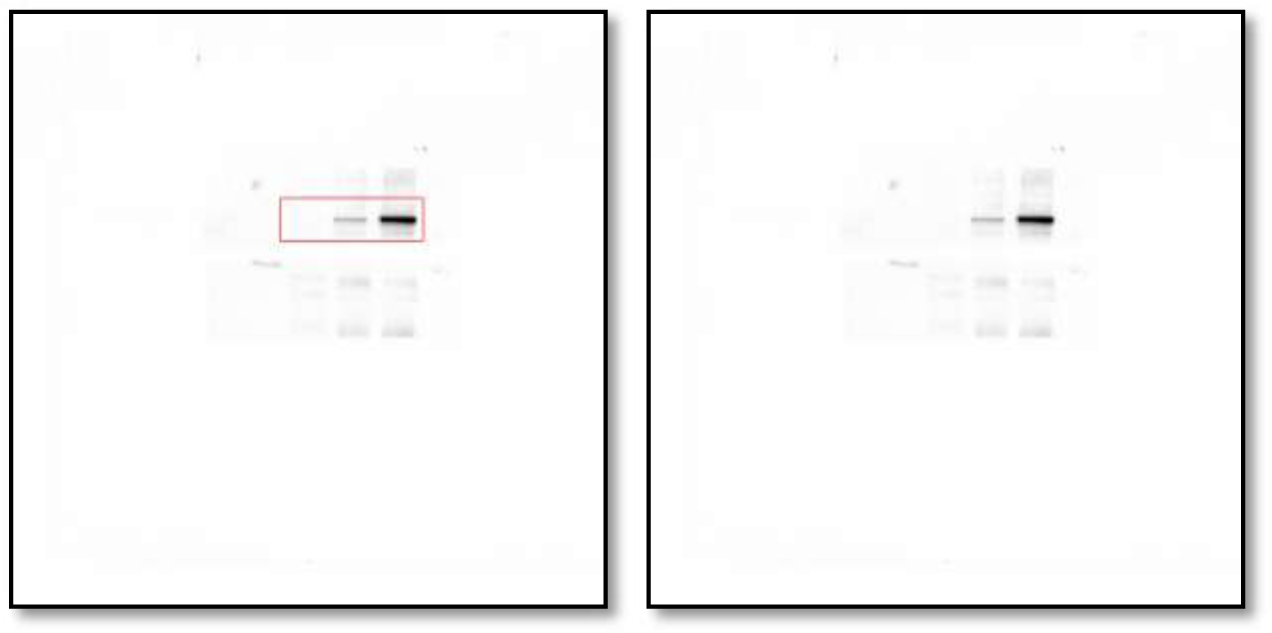
Western blot analysis of E-cadherin protein levels in either vehicle or Pizotifen-treated MDA-MB-231 cells or E-cadherin positive cells in Pizotifen-treated MDA-MB-231 cells.

**Figure 5D_EpCAM_source data.**
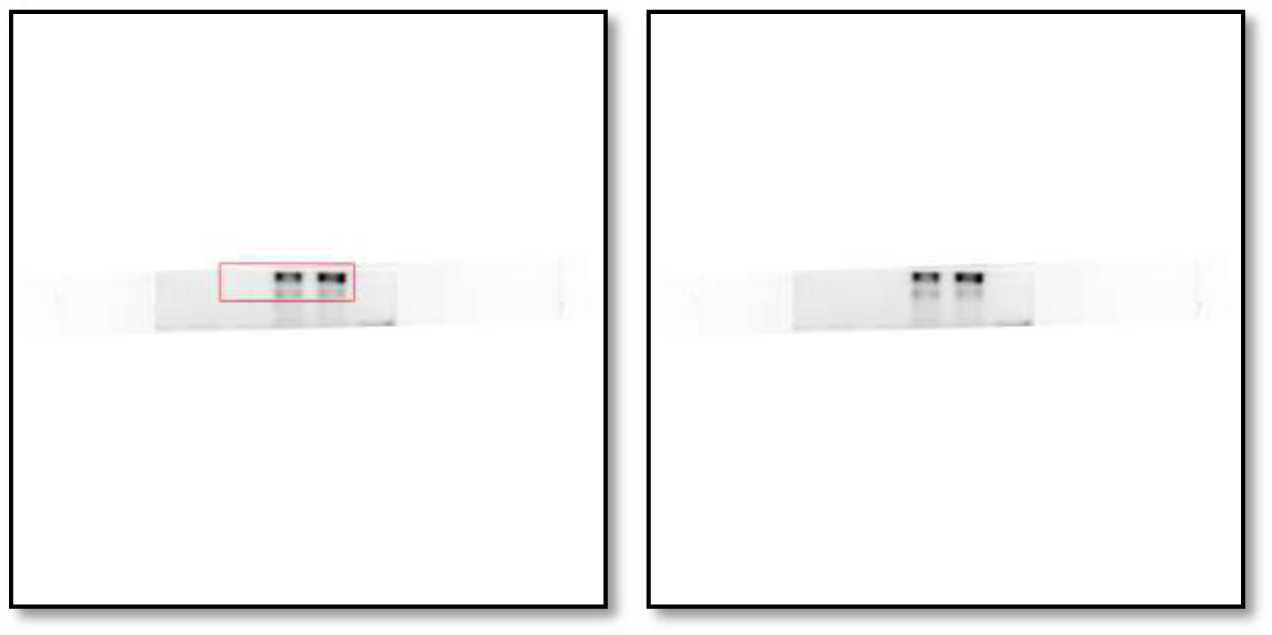
Western blot analysis of EpCAM protein levels in either vehicle or Pizotifen-treated MDA-MB-231 cells or E-cadherin positive cells in Pizotifen-treated MDA-MB-231 cells.

**Figure 5D_Keratin18 (KRT18)_source data.**
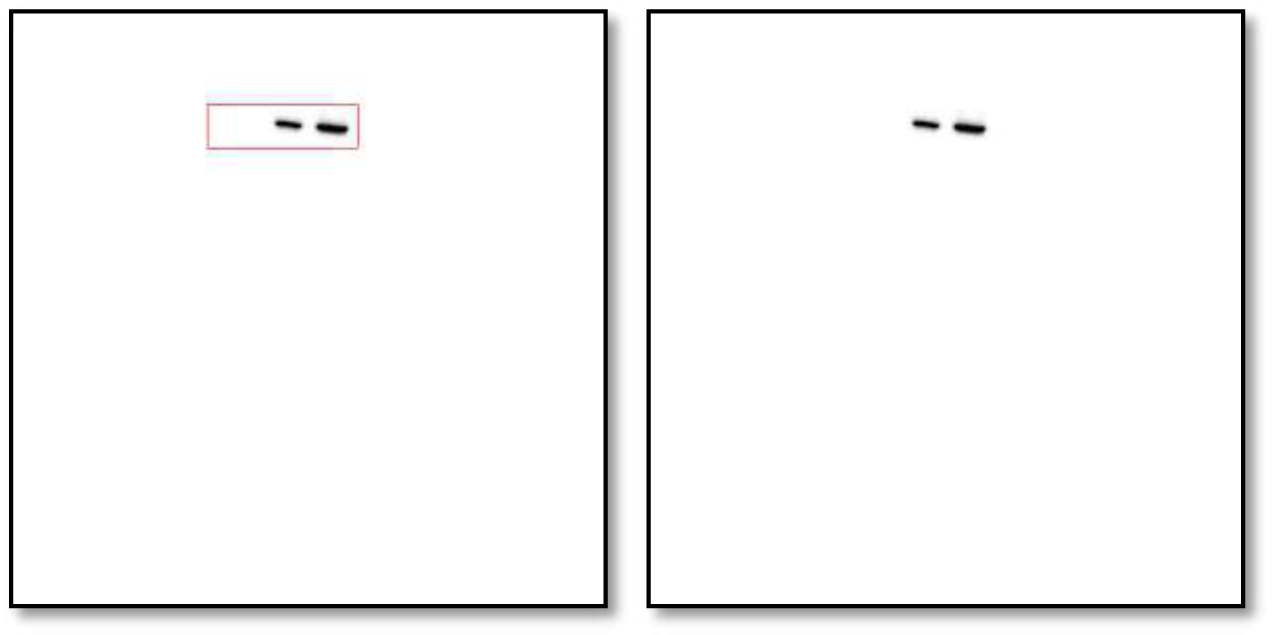
Western blot analysis of KRT18 protein levels in either vehicle or Pizotifen-treated MDA-MB-231 cells or E-cadherin positive cells in Pizotifen-treated MDA-MB-231 cells.

**Figure 5D_Keratin19 (KRT19)_source data.**
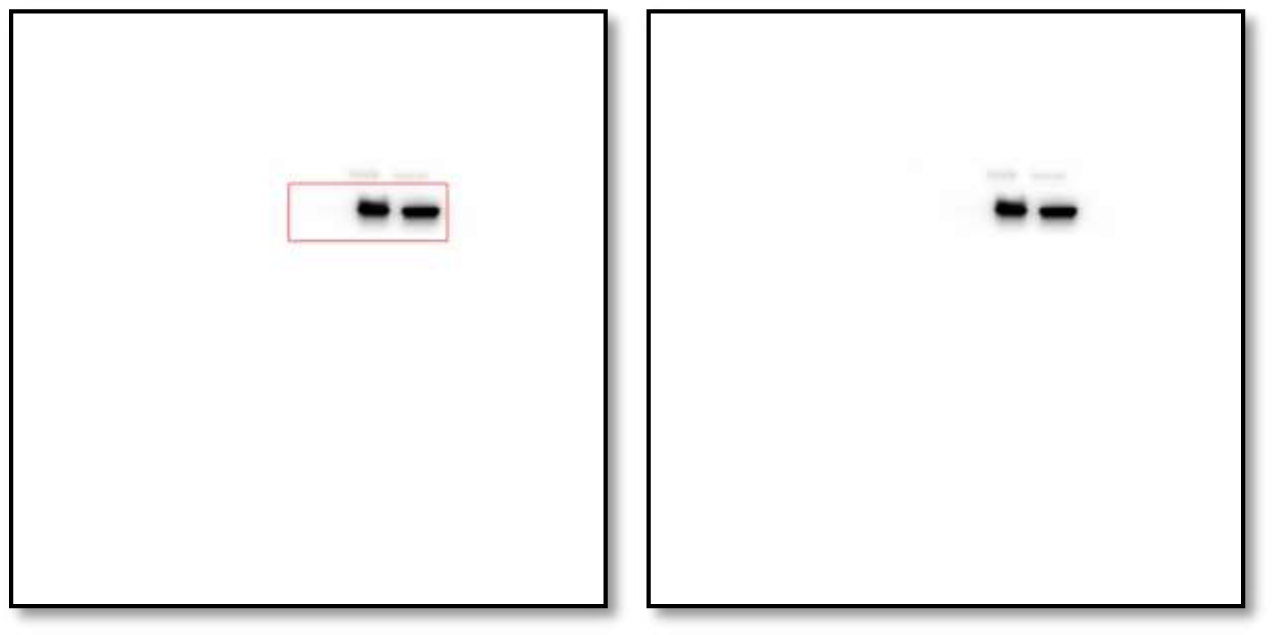
Western blot analysis of KRT19 protein levels in either vehicle or Pizotifen-treated MDA-MB-231 cells or E-cadherin positive cells in Pizotifen-treated MDA-MB-231 cells.

**Figure 5D_Vimentin_source data.**
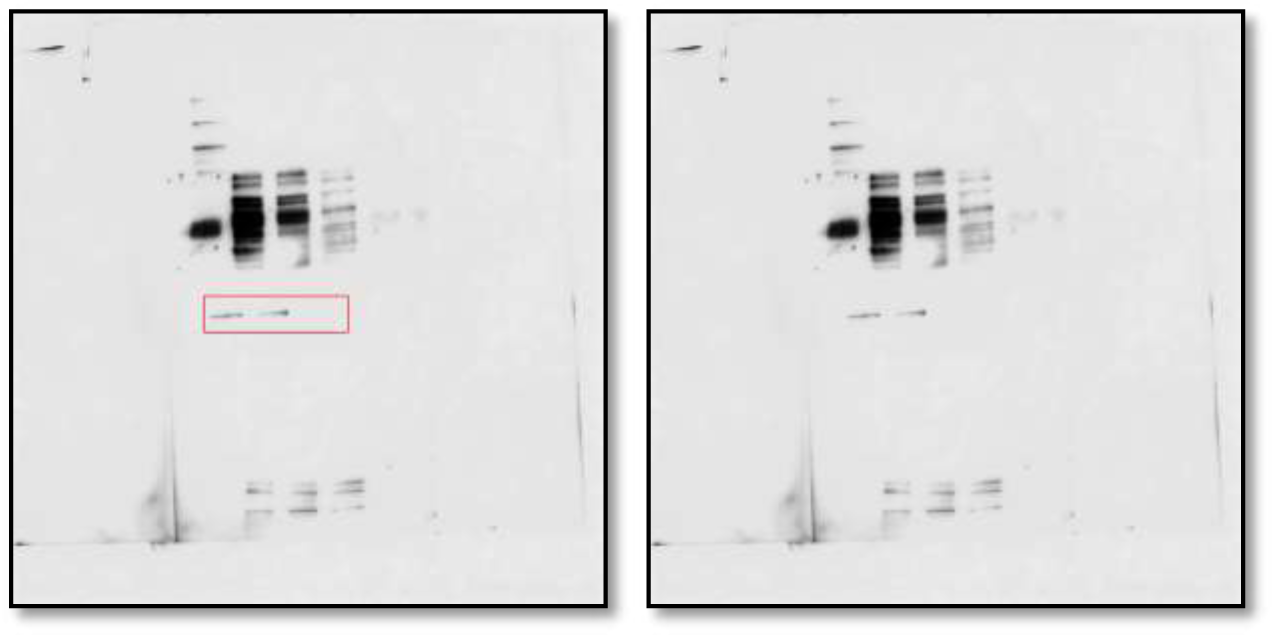
Western blot analysis of Vimentin protein levels in either vehicle or Pizotifen-treated MDA-MB-231 cells or E-cadherin positive cells in Pizotifen-treated MDA-MB-231 cells.

**Figure 5D_MMP1_source data.**
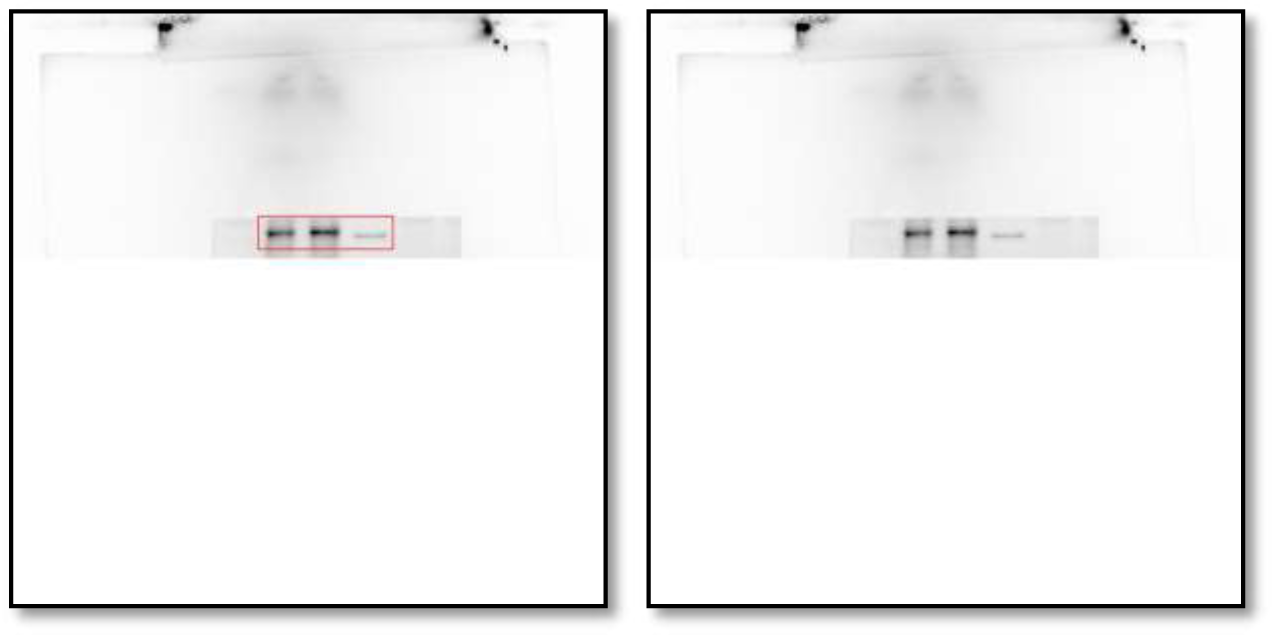
Western blot analysis of MMP1 protein levels in either vehicle or Pizotifen-treated MDA-MB-231 cells or E-cadherin positive cells in Pizotifen-treated MDA-MB-231 cells.

**Figure 5D_MMP3_source data.**
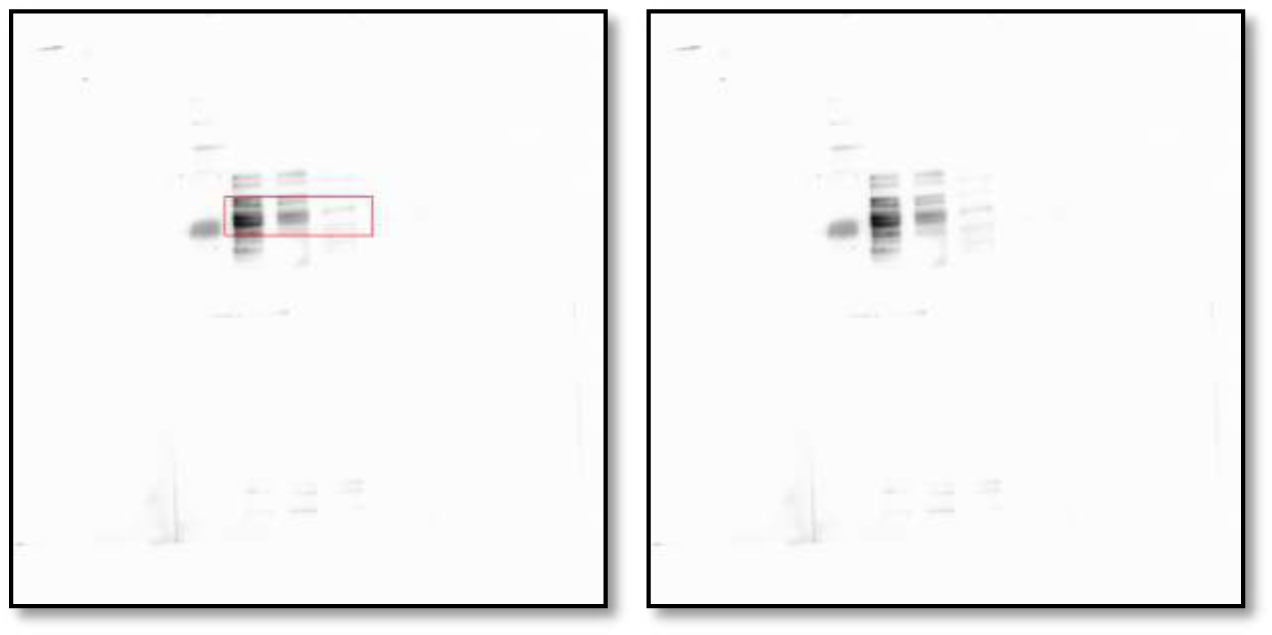
Western blot analysis of MMP3 protein levels in either vehicle or Pizotifen-treated MDA-MB-231 cells or E-cadherin positive cells in Pizotifen-treated MDA-MB-231 cells.

**Figure 5D_S100A4_source data.**
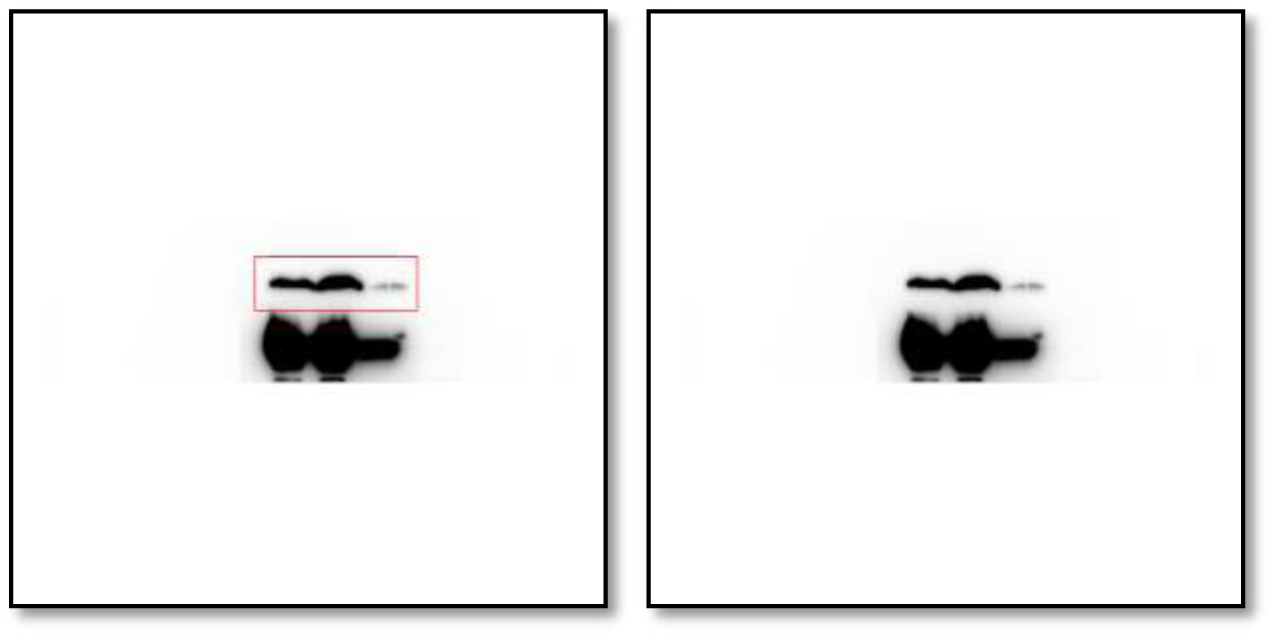
Western blot analysis of S100A4 protein levels in either vehicle or Pizotifen-treated MDA-MB-231 cells or E-cadherin positive cells in Pizotifen-treated MDA-MB-231 cells.

**Figure 5D_Zeb1_source data.**
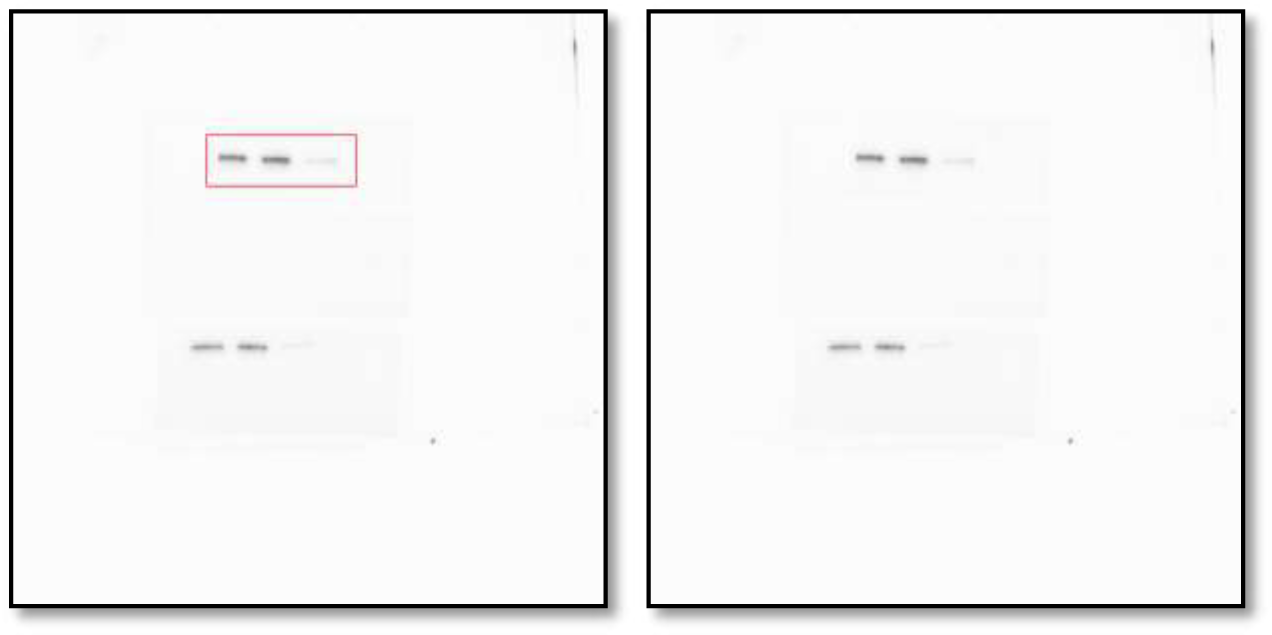
Western blot analysis of Zeb1 protein levels in either vehicle or Pizotifen-treated MDA-MB-231 cells or E-cadherin positive cells in Pizotifen-treated MDA-MB-231 cells.

**Figure 5D_GAPDH_source data.**
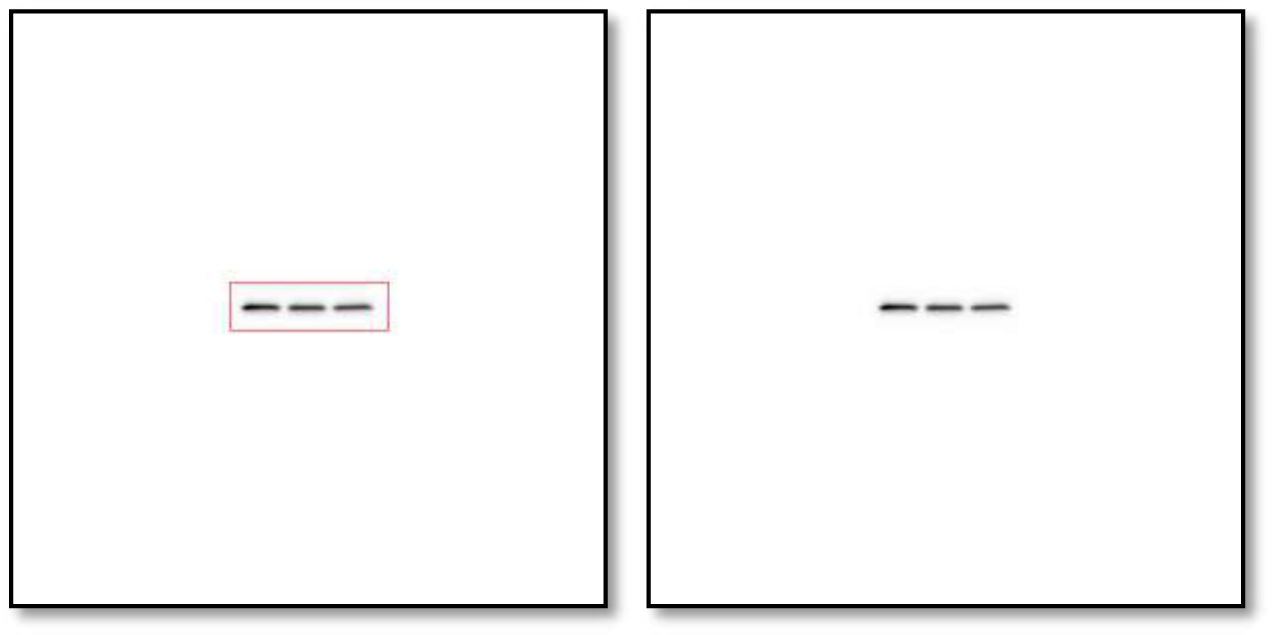
Western blot analysis of GAPDH protein levels in either vehicle or Pizotifen-treated MDA-MB-231 cells or E-cadherin positive cells in Pizotifen-treated MDA-MB-231 cells.

**Figure 5F_β-catenin in the nucleus_source data.**
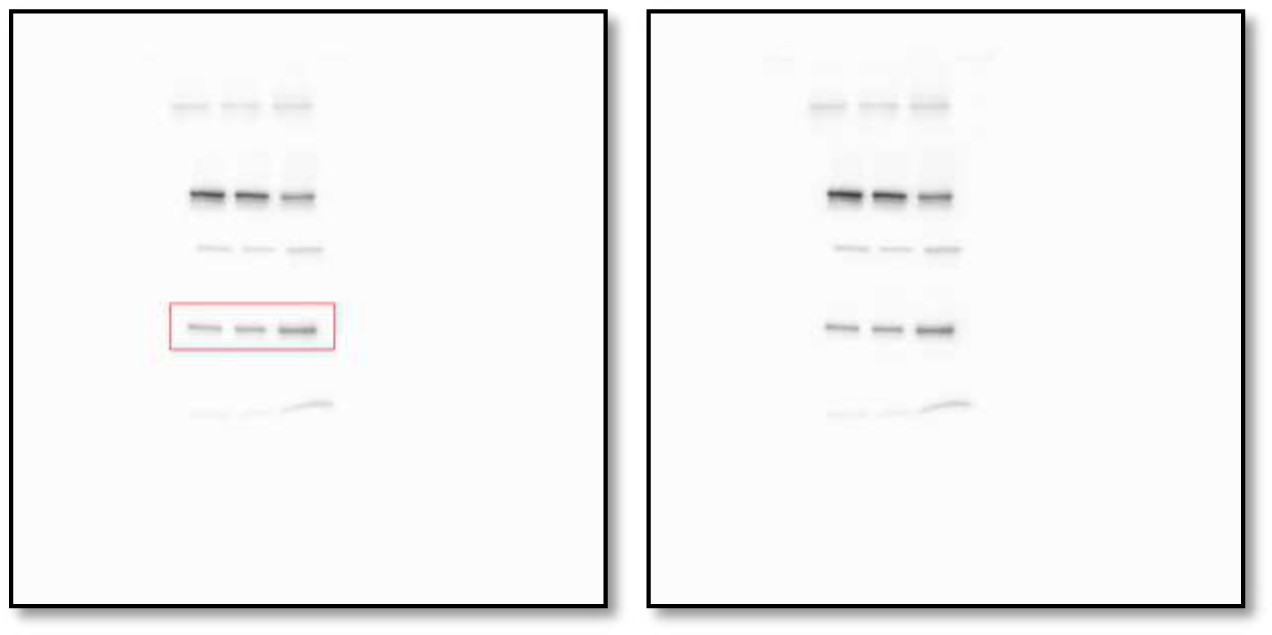
Western blot analysis of β-catenin protein levels in the nuclear of MCF7 cells expressing either the control vector or HTR2C.

**Figure 5F_Histone H3 in the nucleus _source data.**
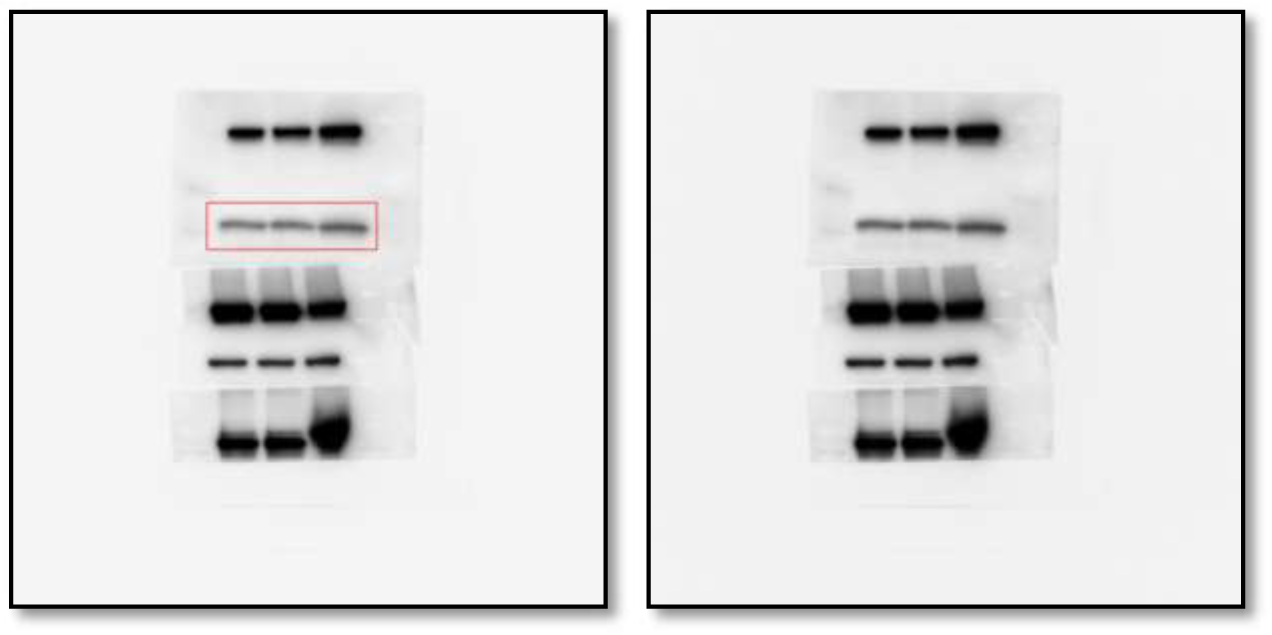
Western blot analysis of Histone H3 protein levels in the nuclear of MCF7 cells expressing either the control vector or HTR2C.

**Figure 5F_β-catenin in the cytoplasm_source data.**
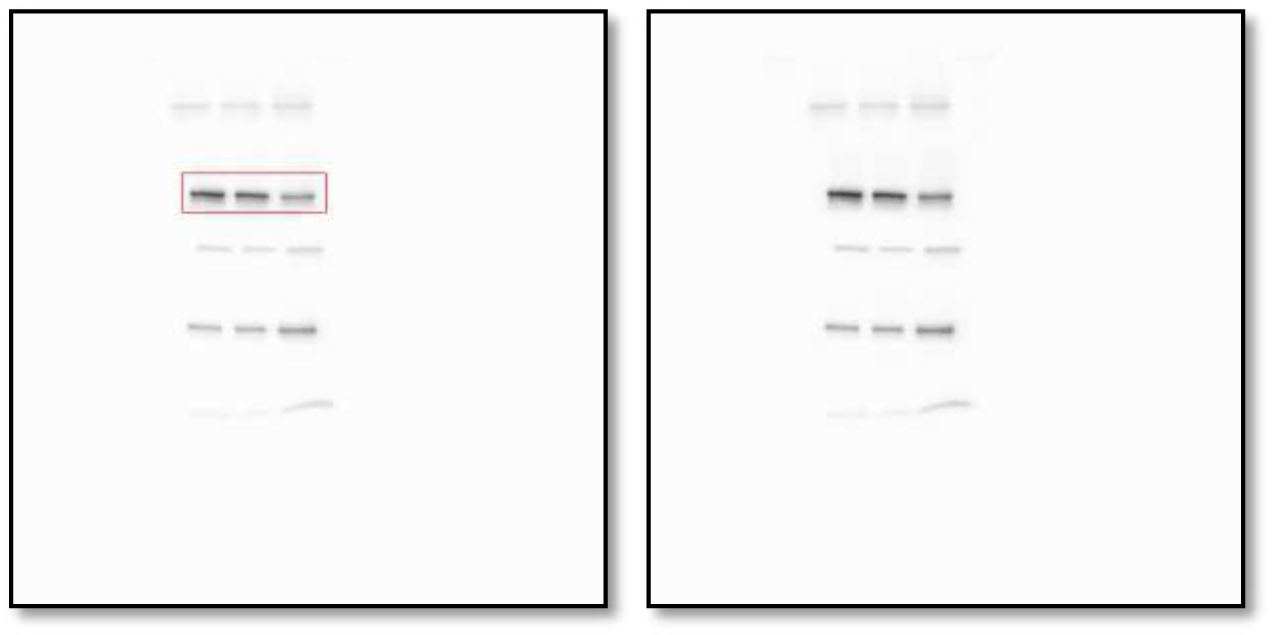
Western blot analysis of β-catenin protein levels in the cytoplasm of MCF7 cells expressing either the control vector or HTR2C.

**Figure 5F_β-tubulin in the cytoplasm_source data.**
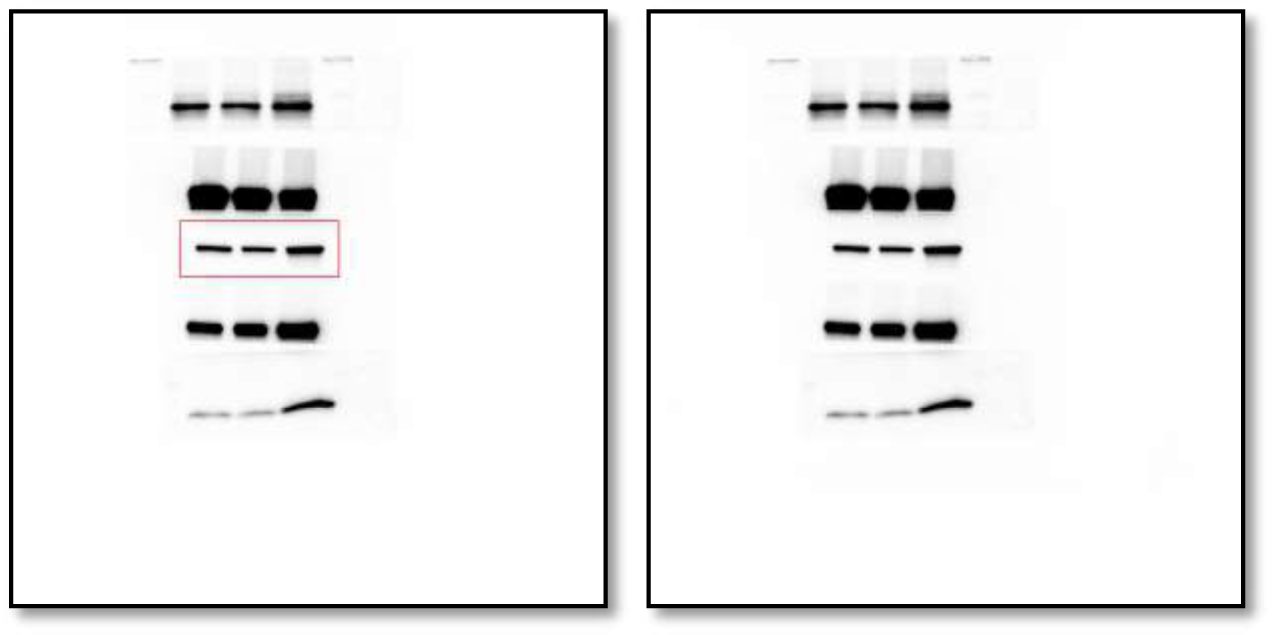
Western blot analysis of β-tubulin protein levels in the cytoplasm of MCF7 cells expressing either the control vector or HTR2C.

**Figure 5F_Phosphorylation of serine-9 in GSK3β_source data.**
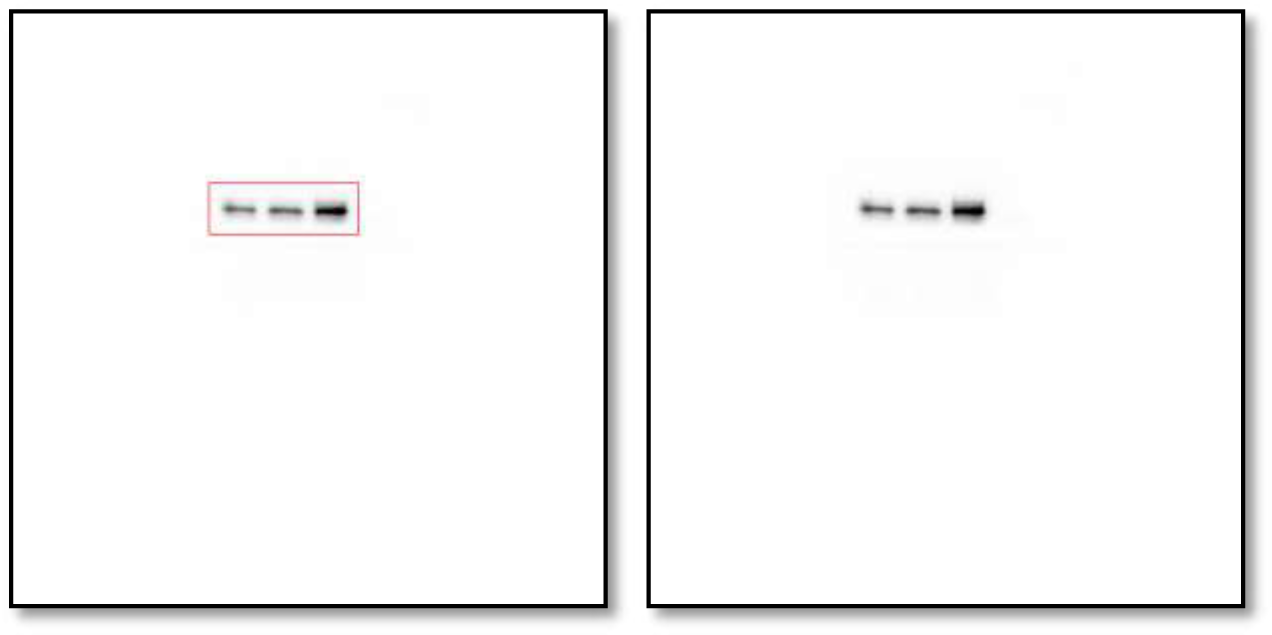
Western blot analysis of the protein levels of phosphorylation of serine-9 in GSK3β in whole cell lysate of MCF7 cells expressing either the control vector or HTR2C.

**Figure 5F_GSK3β_source data.**
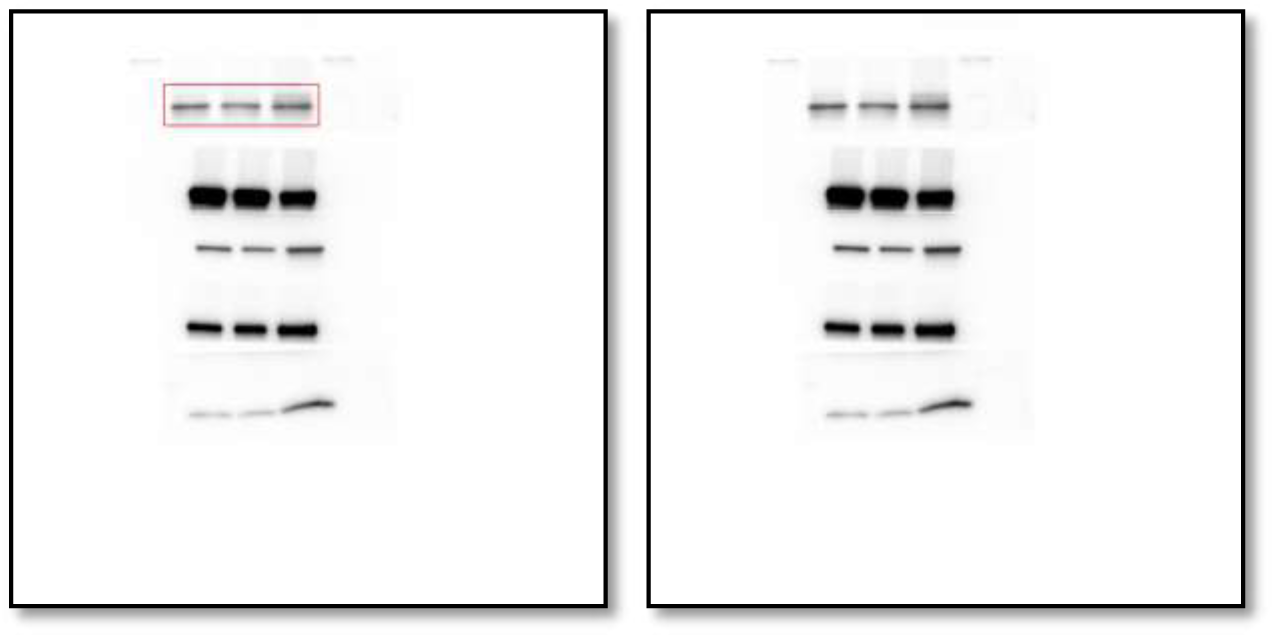
Western blot analysis of GSK3β protein levels in whole cell lysate of MCF7 cells expressing either the control vector or HTR2C.

**Figure 5F_GAPDH_source data.**
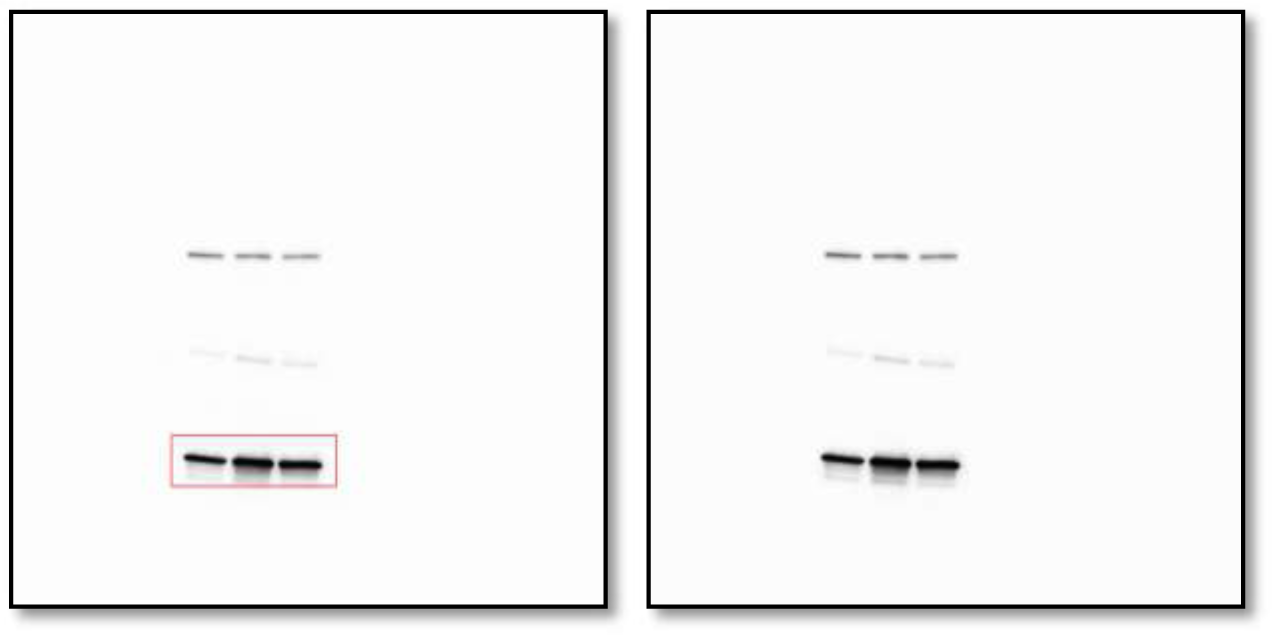
Western blot analysis of GAPDH protein levels in whole cell lysate of MCF7 cells expressing either the control vector or HTR2C.

**Figure 5H_β-catenin in the nucleus_source data.**
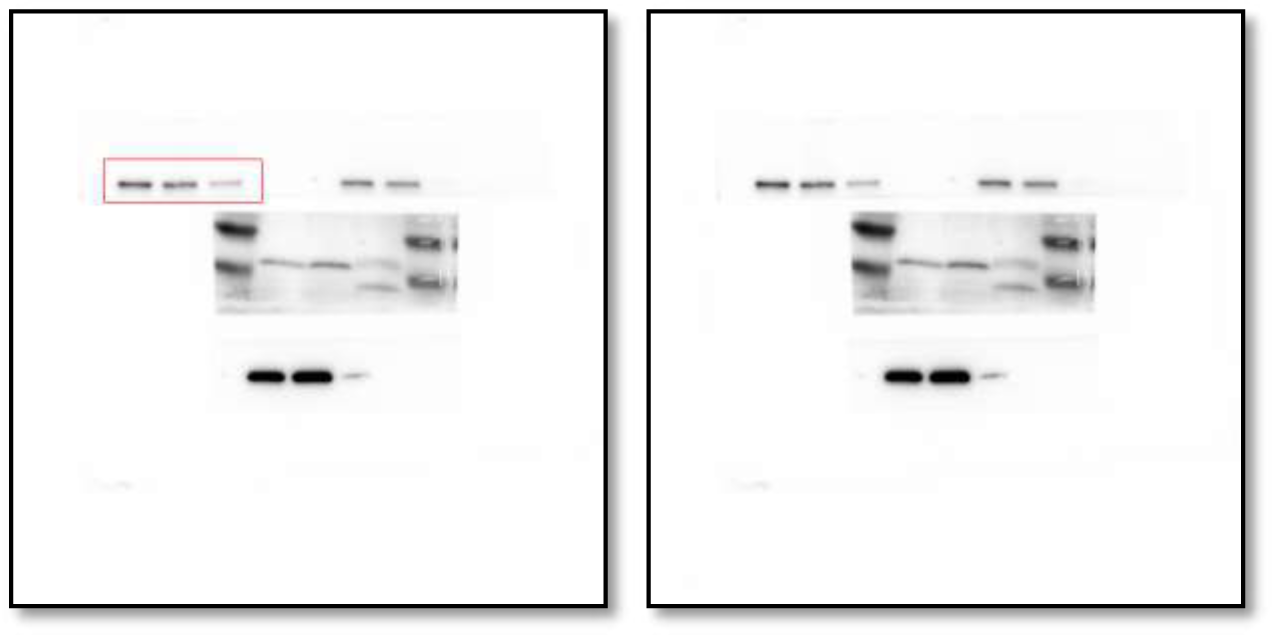
Western blot analysis of β-catenin protein levels in the nucleus of either vehicle or Pizotifen-treated MDA-MB-231 cells or E-cadherin positive cells in Pizotifen-treated MDA-MB-231 cells.

**Figure 5H_Histone H3 in the nucleus _source data.**
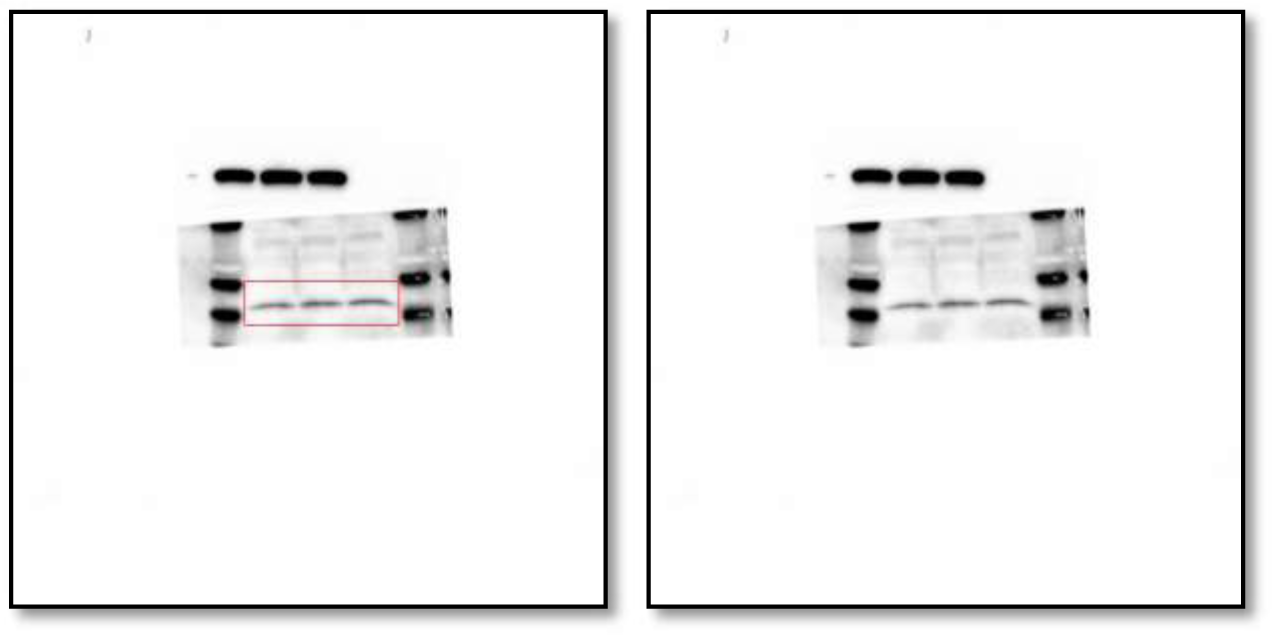
Western blot analysis of Histone H3 protein levels in the nucleus of either vehicle or Pizotifen-treated MDA-MB-231 cells or E-cadherin positive cells in Pizotifen-treated MDA-MB-231 cells.

**Figure 5H_β-catenin in the cytoplasm_source data.**
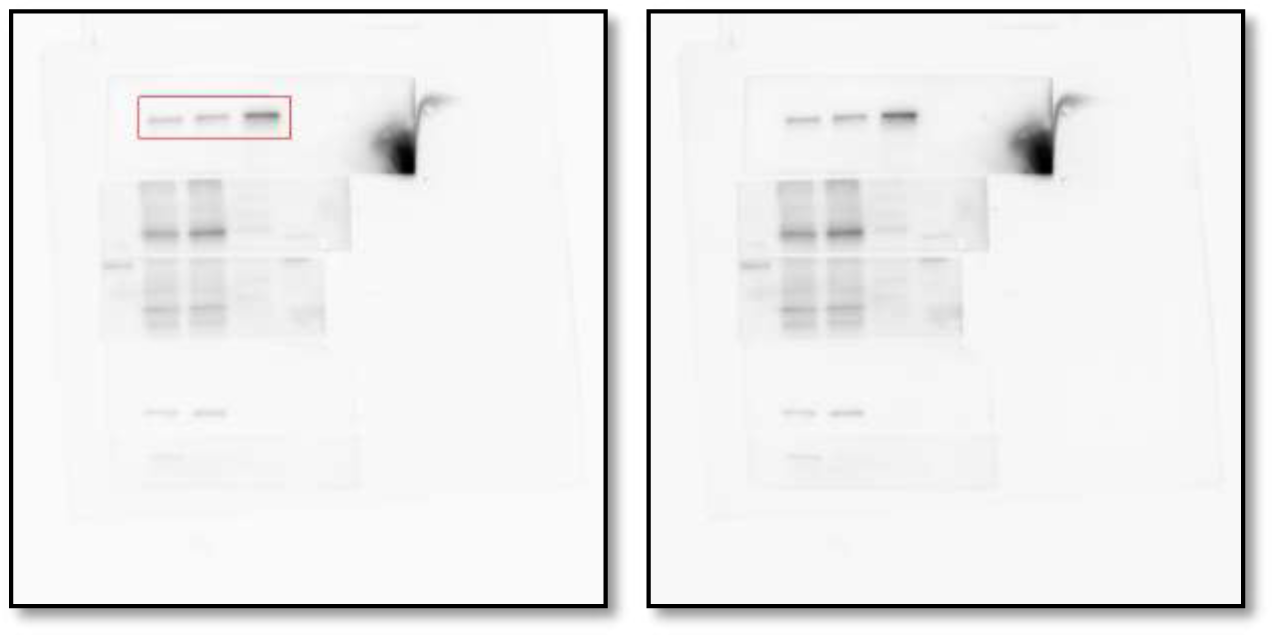
Western blot analysis of β-catenin protein levels in the cytoplasm of either vehicle or Pizotifen-treated MDA-MB-231 cells or E-cadherin positive cells in Pizotifen-treated MDA-MB-231 cells.

**Figure 5H_β-tubulin in the cytoplasm_source data.**
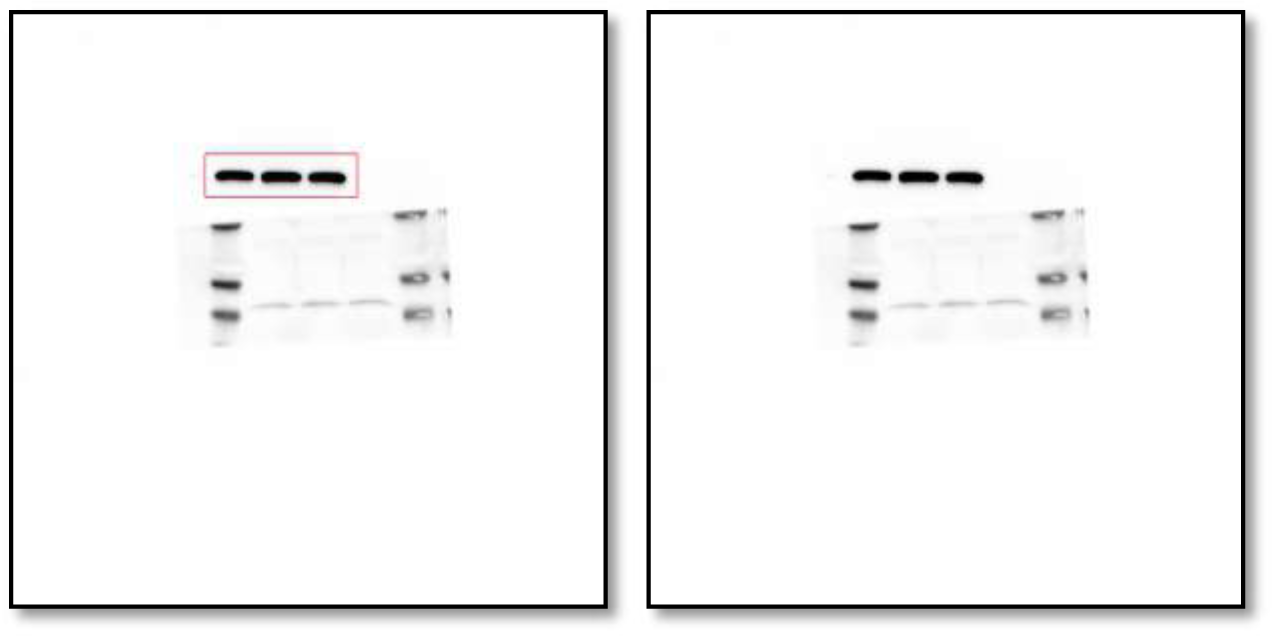
Western blot analysis of β-tubulin protein levels in the cytoplasm of either vehicle or Pizotifen-treated MDA-MB-231 cells or E-cadherin positive cells in Pizotifen-treated MDA-MB-231 cells.

**Figure 5H_Phosphorylation of serine-9 in GSK3β_source data.**
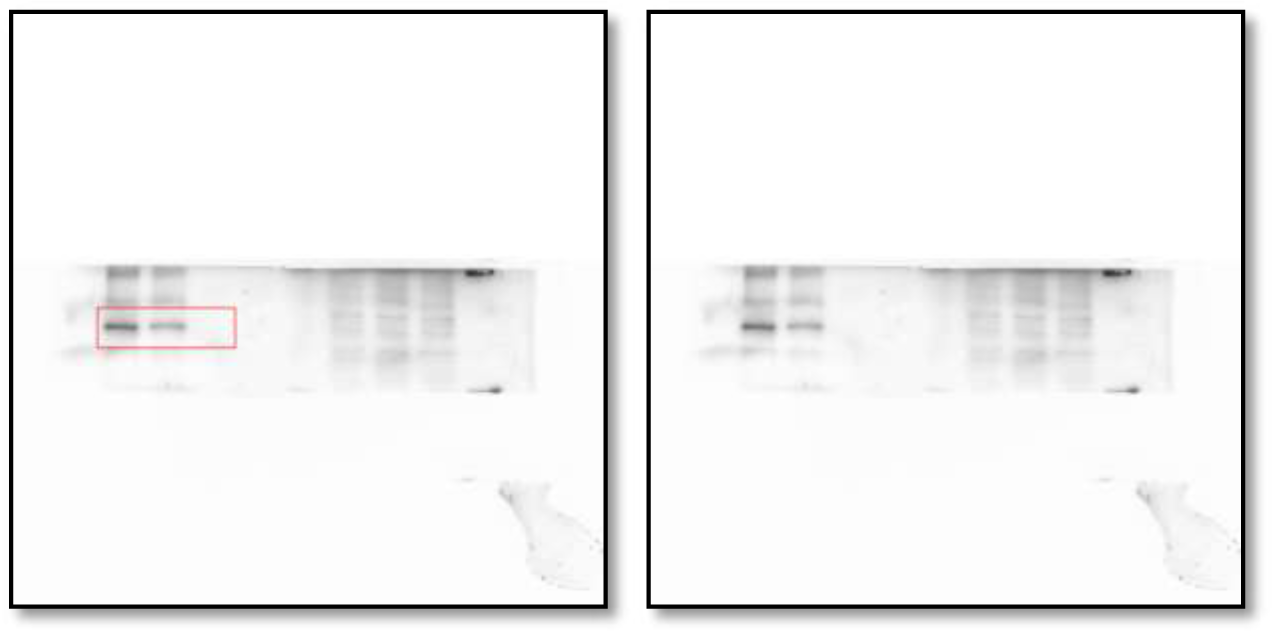
Western blot analysis of the protein levels of phosphorylation of serine-9 in GSK3β in whole cell lysate of either vehicle or Pizotifen-treated MDA-MB-231 cells or E-cadherin positive cells in Pizotifen-treated MDA-MB-231 cells.

**Figure 5H_GSK3β_source data.**
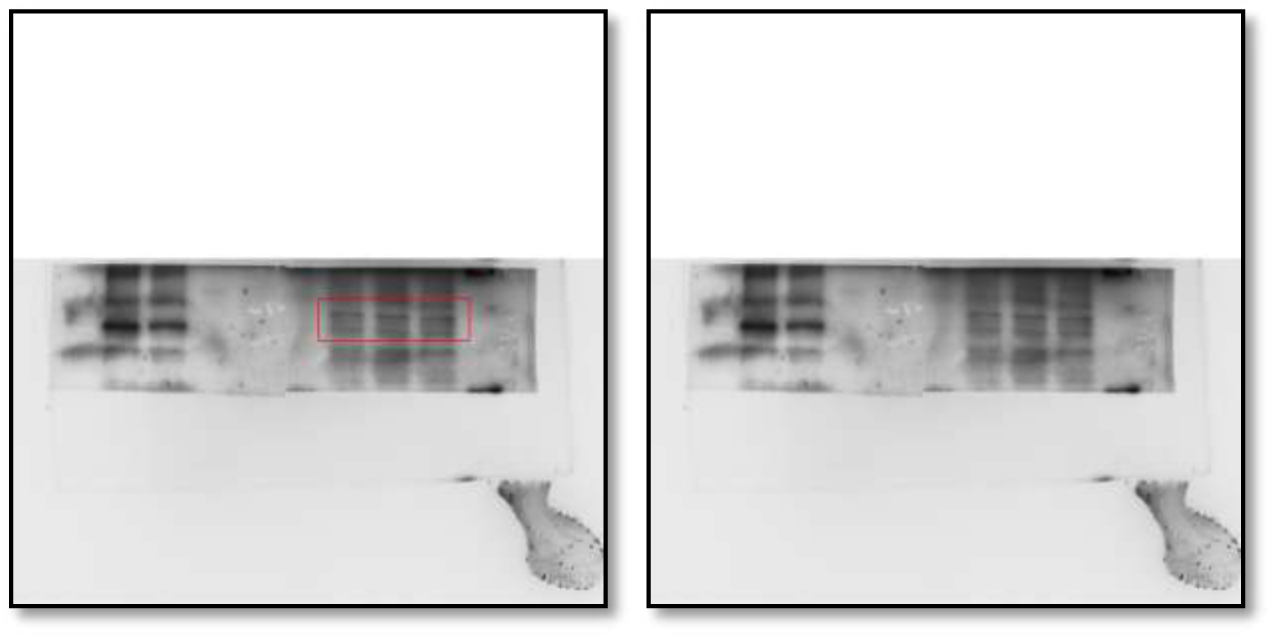
Western blot analysis of GSK3β protein levels in whole cell lysate of either vehicle or Pizotifen-treated MDA-MB-231 cells or E-cadherin positive cells in Pizotifen-treated MDA-MB-231 cells.

**Figure 5H_GAPDH_source data.**
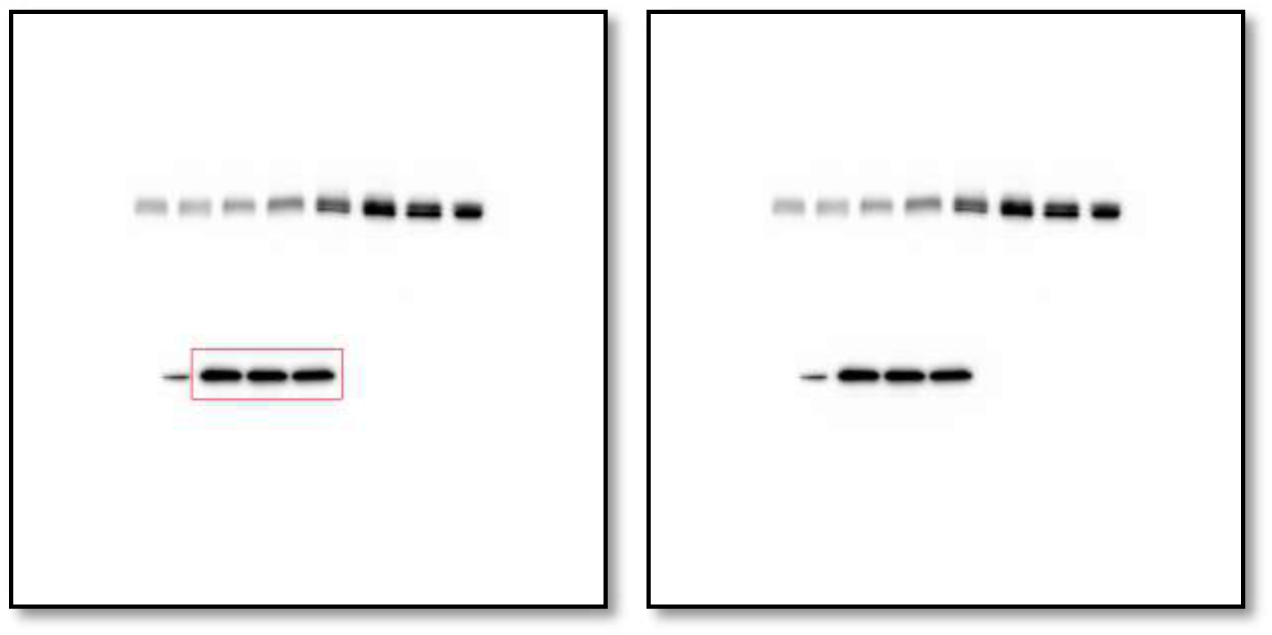
Western blot analysis of GAPDH protein levels in whole cell lysate of either vehicle or Pizotifen-treated MDA-MB-231 cells or E-cadherin positive cells in Pizotifen-treated MDA-MB-231 cells.

**Figure S1A_ PRMT1_source data.**
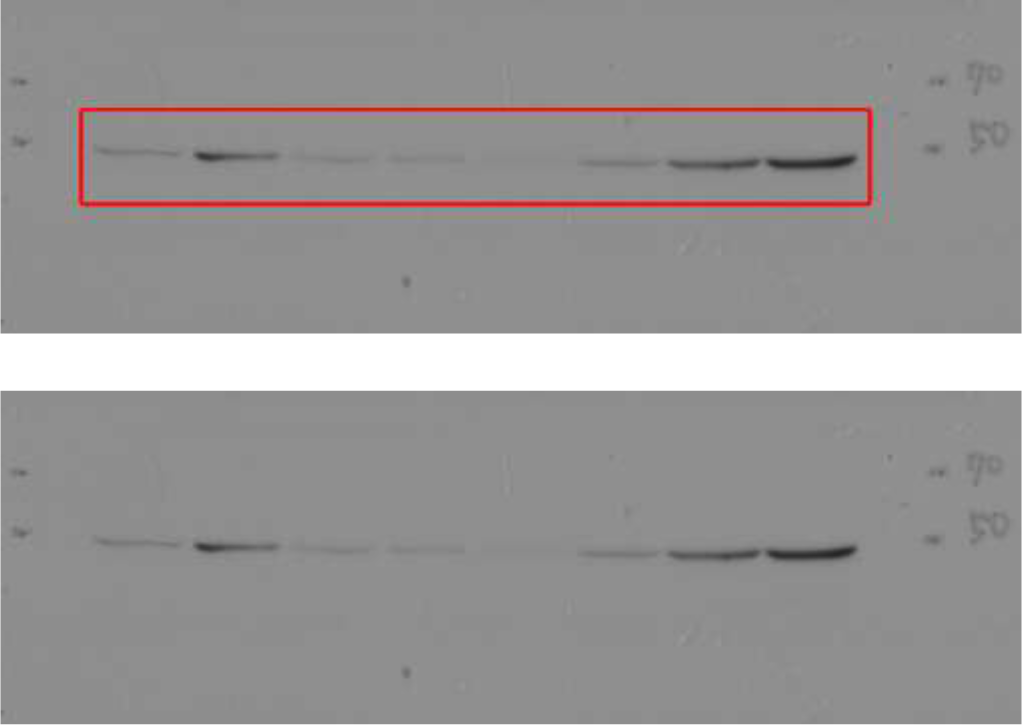
Western blot analysis of PRMT1 protein levels in non-metastatic human cancer cell line (MCF7) and highly metastatic human cancer cell lines (MDA-MB-231, MDA-MB-435, MIA-PaCa2, PC9, HCCLM3, SW620 and PC3).

**Figure S1A_ CYP11A1_source data.**
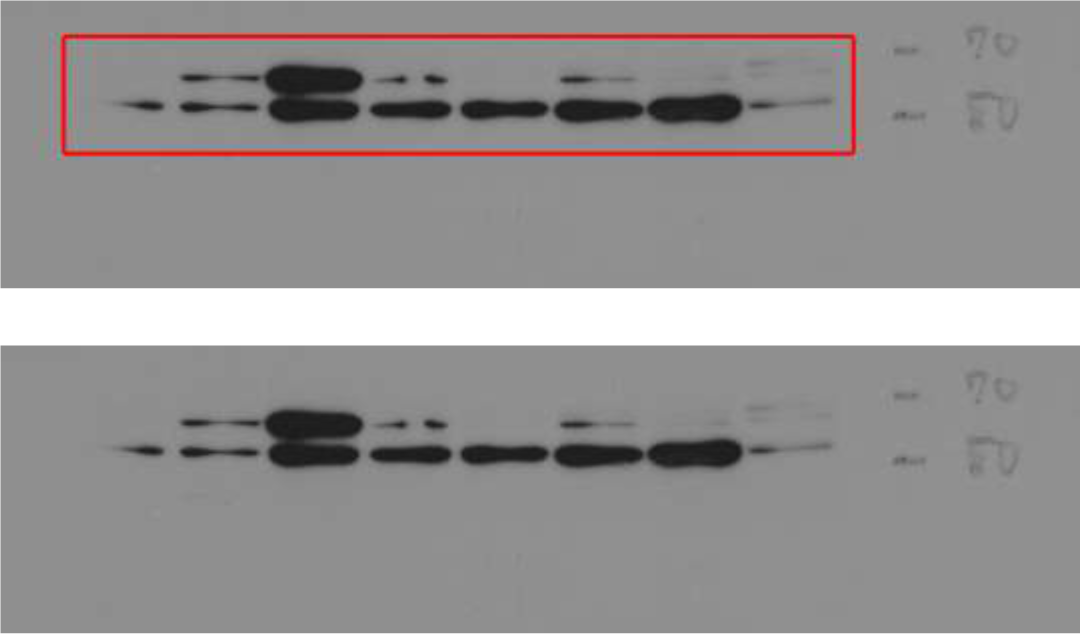
Western blot analysis of CYP11A1 protein levels in non-metastatic human cancer cell line (MCF7) and highly metastatic human cancer cell lines (MDA-MB-231, MDA-MB-435, MIA-PaCa2, PC9, HCCLM3, SW620 and PC3).

**Figure S1A_ β-actin_source data.**
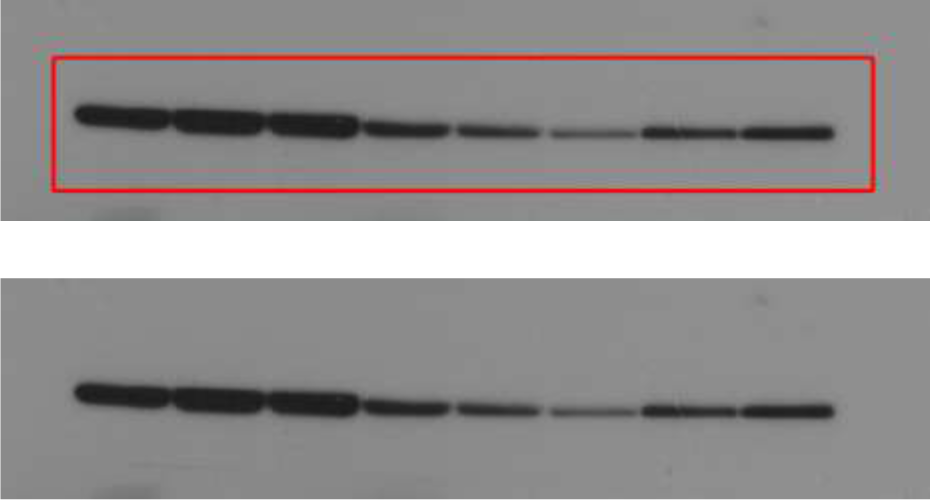
Western blot analysis of β-actin protein levels in non-metastatic human cancer cell line (MCF7) and highly metastatic human cancer cell lines (MDA-MB-231, MDA-MB-435, MIA-PaCa2, PC9, HCCLM3, SW620 and PC3).

**Figure S1A_ PRMT1_source data.**
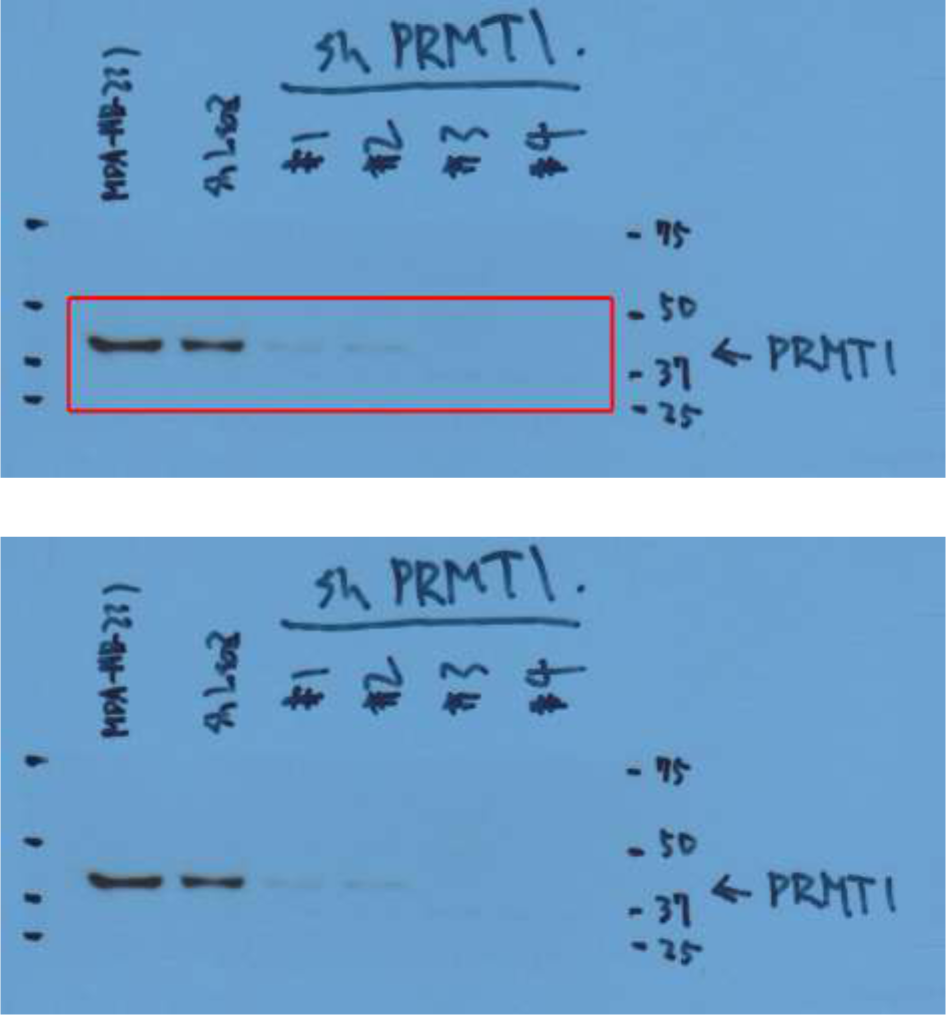
Western blot analysis of PRMT1 protein levels in sub-clones of MBA-MB-231 cells which were transfected with either a control shRNA targeting LacZ or one of four independent shRNAs targeting PRMT1 (clone #1 to #4).

**Figure S1A_ β-actin _source data.**
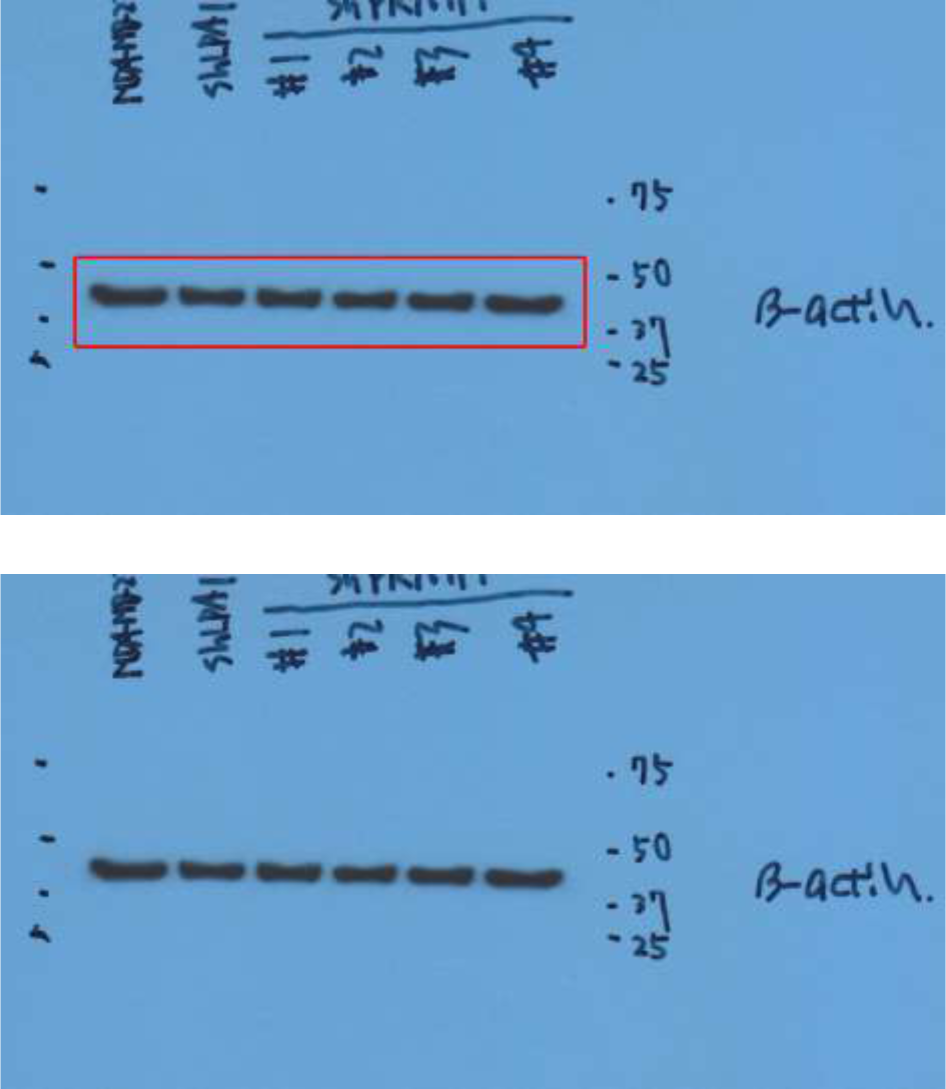
Western blot analysis of β-actin protein levels in sub-clones of MBA-MB-231 cells which were transfected with either a control shRNA targeting LacZ or one of four independent shRNAs targeting PRMT1 (clone #1 to #4).

**Figure S1A_CYP11A1_source data.**
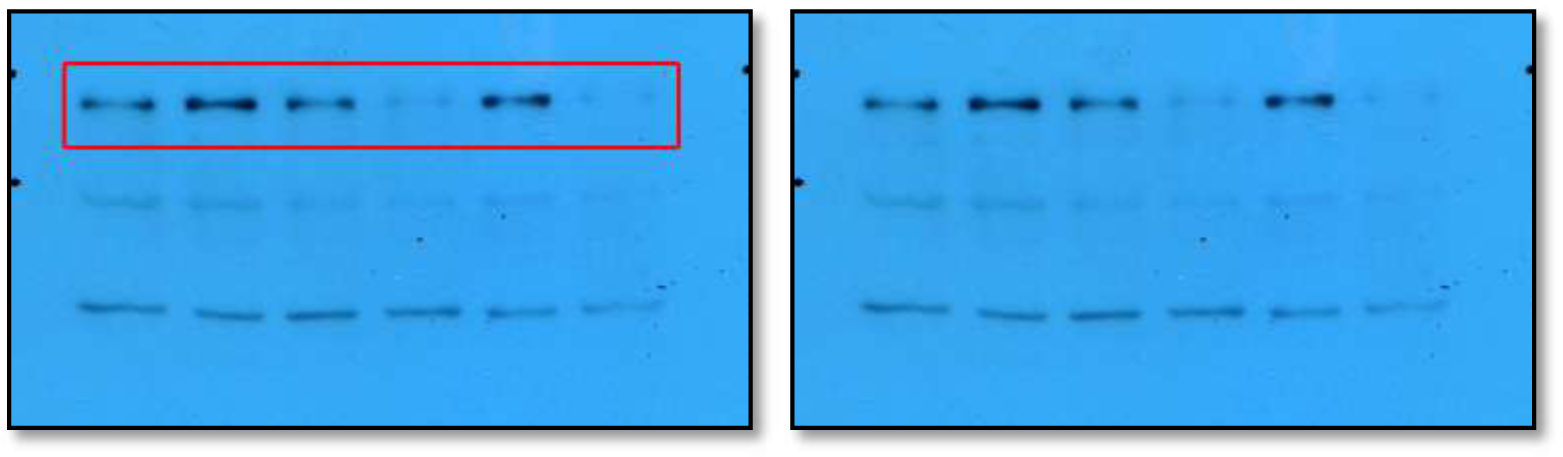
Western blot analysis of CYP11A1 protein levels in sub-clones of MBA-MB-231 cells which were transfected with either a control shRNA targeting LacZ or one of four independent shRNAs targeting CYP11A1 (clone #1 to #4).

**Figure S1A_ β-actin_source data.**
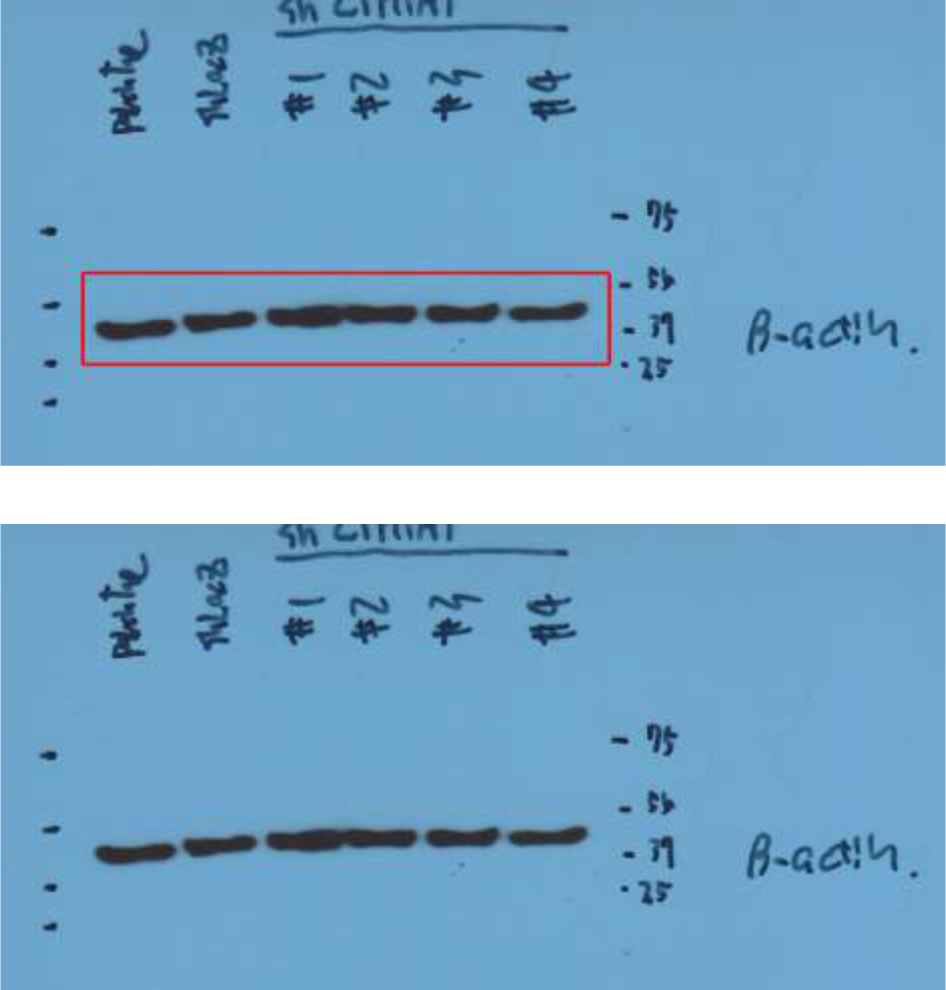
Western blot analysis of β-actin protein levels in sub-clones of MBA-MB-231 cells which were transfected with either a control shRNA targeting LacZ or one of four independent shRNAs targeting CYP11A1 (clone #1 to #4).

**Figure S2A_DRD2_source data.**
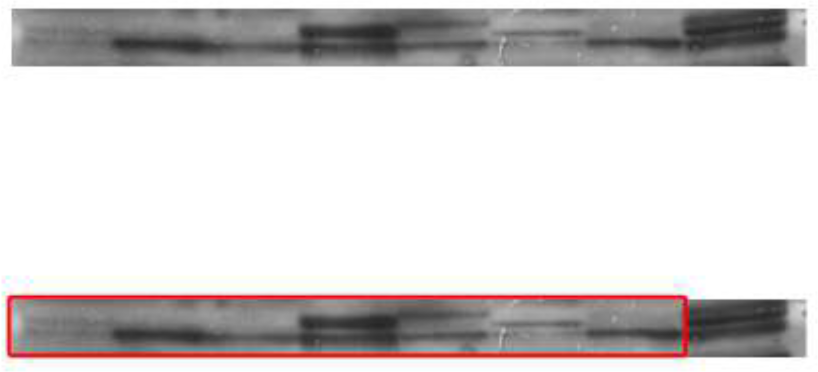
Western blot analysis of GAPDH protein levels in non-metastatic human cancer cell line, MCF7 (breast) and highly metastatic human cancer cell lines, MDA-MB-231 (breast), MDA-MB-435 (melanoma), MIA-PaCa2 (pancreas), PC3 (prostate) and SW620 (colon)

**Figure S2A_DRD2_source data.**
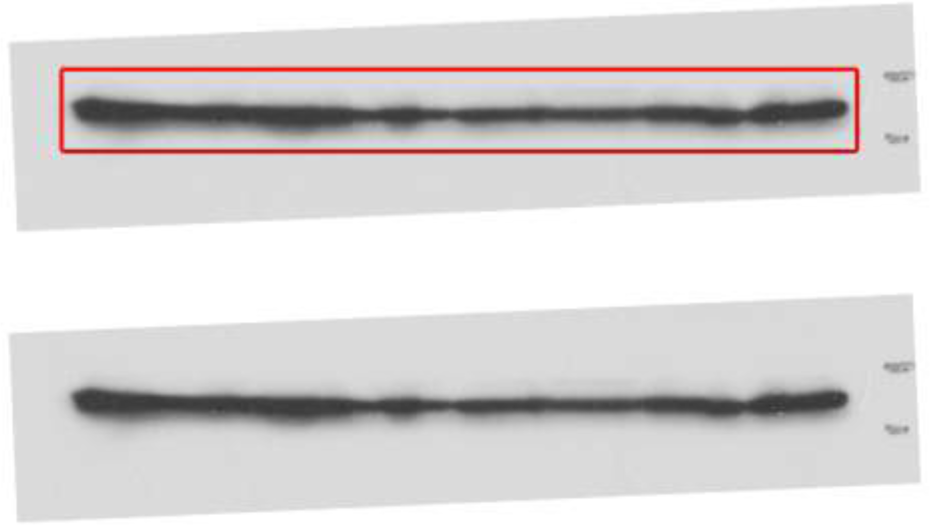
Western blot analysis of DRD2 protein levels in non-metastatic human cancer cell line, MCF7 (breast) and highly metastatic human cancer cell lines, MDA-MB-231 (breast), MDA-MB-435 (melanoma), MIA-PaCa2 (pancreas), PC3 (prostate) and SW620 (colon)

